# Thille, a chromosome-level type B avocado genome, and the structural and allelic variation at candidate flowering-type loci

**DOI:** 10.64898/2026.10.12.756161

**Authors:** Edwin Solares, Edward Tang, Brianna Sanchez, Gavin Simmons, Kathryn Chen, Zachary Hom, Nichole Mora, Rosa Figueroa-Balderas, Eric Focht, Dario Cantu, Mary L. Arpaia

**Affiliations:** Department of Computer Science, University of California, Davis, Davis, CA, 95616; Department of Computer Science and Engineering, University of California San Diego, La Jolla, CA 92093; ESB AI Lab, San Diego, CA, 92116; Botany and Plant Sciences Department, University of California, Riverside, CA 92521; Department of Viticulture and Enology, University of California, Davis, Davis, CA, 95616; Genome Center, University of California, Davis, Davis, CA, 95616

## Abstract

Avocado (Persea americana Mill.) is heterodichogamous: each flower opens first as female and then as male, and cultivars fall into two complementary flowering types, A and B, determined by the transcription factor SDMYB, whose dominant allele confers type A. Published avocado genomes come from type A cultivars or from accessions of unreported flowering type. Here we report a chromosome-level genome of Thille, a type B cultivar used in breeding. From 72.2 Gb of PacBio HiFi reads (81x coverage) and Dovetail Omni-C data, the assembly places 885.4 Mb, 98.8% of the expected 896 Mb genome, in 12 chromosomes with an NG50 of 84.5 Mb, a consensus quality of QV 70.02, and 99.7% complete BUSCO genes; repeats occupy 64.0% of the assembly, and 36,349 protein-coding genes were predicted. We compared Thille with the type A cultivars Hass and Puebla, a Guatemalan x Mexican hybrid, and with Unnamed Lowland, an accession of the Lowland botanical group (formerly West Indian) of unreported flowering type. The assemblies differ by large rearrangements, including a 22.6 Mb inversion on chromosome 12 specific to Hass which is most likely due to scaffolding error. Using phased reads and haplotype assemblies, we revisited and reconstructed the alleles of candidate flowering genes from the type A versus type B Fst scan from our previous work. Both Unnamed Lowland haplotypes carry the recessive SDMYB allele found in Thille; based on this study and the previous study of SDMYB, we predict Unnamed Lowland to be type B. We also observe five other genes at which two type A cultivars (Puebla and Gwen) share an allele absent from Thille and Unnamed Lowland, a pattern that requires further investigation. The genome and HiFi reads have been submitted to NCBI under BioProject PRJNA1142556.

**Summary:** The avocado (Persea americana) is known for its synchronous dichogamous flowering system, in which cultivars are classified as Type A or Type B based on when their female and male reproductive organs mature, a timing offset that encourages cross-pollination (Stout, 1927; Davenport, 1986). While the Type A genome has been extensively studied, primarily through the widely cultivated Hass variety, this study presents the first nearly complete genome of a documented Type B avocado, derived from the Thille variety. Due to Type B being homozygous recessive (Groh et al., 2026), a Type B assembly carries only the recessive allele at the flowering-type locus. Assembling this genotype therefore avoids the allelic heterozygosity expected at that locus in Type A individuals, where the dominant and recessive haplotypes must either be phased or risk collapsing into a single consensus sequence, and it provides an unambiguous reference for the recessive haplotype. The assembly reaches 98.8% completeness, with the 12 largest scaffolds approximating pseudomolecule length and an NG50 of 84.5 Mb. Our analysis reveals structural variants, including inversions and deletions, between the Thille and Hass genomes, particularly on chromosome 10, which houses key flowering markers. This genome and its HiFi sequencing data enrich the genomic resources available for avocado, supporting breeding and further research into this agriculturally significant and resource-intensive crop (Solares et al., 2023).

## Introduction

The avocado (Persea americana Mill.) has a synchronous dichogamous flowering system, heterodichogamy, in which each flower opens twice, first as functionally female and then as functionally male, and cultivars fall into two complementary flowering types, A and B, whose female and male phases alternate within the same day, a timing offset that encourages cross-pollination between types (Stout, 1927; Davenport, 1986; Renner, 2001). Flowering type behaves as a single locus (Groh et al., 2026). Linkage mapping placed a quantitative trait locus for flowering type on linkage group 10 (Ashworth et al., 2019), and Solares et al. (2023) listed the gene as a candidate flowering gene among the top 1% Fst genes in their supplementary tables (Chr10.ver1.g56860, EOBI-like). Groh et al. (2026) mapped the locus by GWAS and narrowed it to SDMYB using shared heterozygosity among resequenced A-type cultivars, including those of Solares et al. (2023); SDMYB is a MYB transcription factor on chromosome 10 whose dominant allele confers type A, so that type A individuals are heterozygous, and on rare occasions homozygous for the dominant allele, and type B individuals are homozygous for the recessive allele. Khan et al. (2026) independently associated flowering type with a 20 kb region on chromosome 10 containing an R2R3 MYB-like transcription factor, PaDSPD. Genomic resources for avocado have been built almost entirely from type A cultivars. Hass, the type A cultivar that dominates commercial production, has assemblies of increasing contiguity (Rendón-Anaya et al., 2019; Nath et al., 2022), and the genome of Gwen, a close relative of Hass, was used to study the domestication of the three botanical groups of avocado, Mexican, Guatemalan, and Lowland (formerly West Indian), and to compare type A and type B accessions (Solares et al., 2023). Haplotype-resolved assemblies of the type A cultivar Puebla were produced to characterize the flowering-type locus (Groh et al., 2026), and a telomere-to-telomere assembly of an unnamed accession of the Lowland group of unreported flowering type (Unnamed Lowland) has also been published (Yang et al., 2024). To our knowledge, no genome of a documented type B cultivar has been published; in this study, Unnamed Lowland, the Lowland T2T accession, we find to also be homozygous for the recessive SDMYB allele, which we predict to be a flower type B. A type B genome is useful beyond its rarity: because type B is homozygous recessive at SDMYB (Groh et al., 2026), a type B assembly carries only the recessive allele at the flowering-type locus, avoiding the heterozygosity that a type A assembly must either phase or collapse into a single consensus, and it provides an unambiguous reference for the recessive haplotype. Here we report a chromosome-level assembly of the type B cultivar Thille, a variety planted from an open pollinated Hass seed in 1946 by J.N. Thille in Santa Paula, California (Delphey, 1960). The assembly places 98.8% of the expected 896 Mb genome in the 12 largest scaffolds, which correspond to the 12 avocado chromosomes, with an NG50 of 84.5 Mb. We compare Thille with the Hass, Puebla, and Lowland assemblies to describe structural variants between them, including a large inversion on chromosome 12, and examine the read coverage of Thille HiFi, Gwen CLR, and Lowland HiFi reads at candidate flowering genes from the type A versus type B Fst scan of the Gwen study. Using phased reads and haplotype assemblies, we reconstruct the alleles of these genes in Thille, Unnamed Lowland, and three type A cultivars, determine the SDMYB genotype of Unnamed Lowland, which we also attempt to place in a flowering type at several other gene loci, and report the alleles that the type A cultivars share. We provide this genome and its HiFi sequencing data to the research and breeding community as a resource for this agriculturally significant and resource-intensive crop (Solares et al., 2023).

## Methods

### Material Collection

Young leaves and meristems for PacBio HiFi sequencing were collected from a Thille tree in field 22, row 32, tree 10, at the UC Agriculture and Natural Resources (UCANR) South Coast Research and Extension Center (SCREC) in Irvine, California. Leaves were placed in sealed freezer bags, packed in dry ice, and transported by E.S. to the University of California, Davis, where they were stored at −80°C in the Ross-Ibarra laboratory. Samples were subsequently transferred to the Cantu laboratory (R. Figueroa-Balderas) and held at −80°C until DNA extraction. For Omni-C library preparation, young leaves were collected from the same Thille tree by E. Focht, packed in dry ice, and shipped to Dovetail Genomics (Scotts Valley, California) for chromatin extraction and Dovetail Omni-C library construction; Omni-C is a variant of the Hi-C chromatin conformation capture method.

### DNA Extraction, Purification and Sequencing

High Molecular Weight genomic DNA (gDNA) was extracted from young leaves using the method described by Solares et al. (2023), based on Chin et al. (2016). DNA purity was assessed with a Nanodrop spectrophotometer (Thermo Scientific, IL, USA), and DNA concentration was quantified using the DNA High Sensitivity Kit on a Quibit 2.0 Fluorometer (Life Technologies, CA, USA). DNA Integrity was evaluated using an Agilent FEMTO Pulse system (Agilent, CA, USA). HiFi sequencing libraries were prepared following the manufacturer’s protocol for the SMRTbell Express Template Prep Kit 3.0 (Pacific Biosciences, CA, USA). Libraries were size-selected using 3.1 X v/v of 35% Ampure PB beads (Pacific Biosciences, CA, USA). Concentration and final size distribution of the library were evaluated using a Qubit and FEMTO respectively. Libraries were sequenced on four SMRT cells on a PacBio Sequel IIe at the DNA Technologies Core, University of California, Davis, and read statistics were generated with seqkit v2.13.0 (Shen et al., 2016).

### Genome Assembly Methods

Genome assembly was performed using PacBio HiFi reads with a per-read quality of at least Q20 (Figure 1B), combined from the four SMRT cells into one FASTQ file (81x coverage). The reads were assembled with hifiasm v0.25.0 (Cheng et al., 2021, 2022) using default HiFi and Hi-C parameters (where pertinent), both with and without Omni-C reads during assembly. Dovetail Omni-C reads were aligned to each primary assembly using BWA-MEM v0.7.19 (Li, 2013) with the −5SP options, and the output was piped to SAMtools view v1.21 (Danecek et al., 2021) to generate BAM files. Omni-C scaffolding was performed using YaHS v1.2.2 (Zhou et al., 2023) on both primary assemblies with the BWA-MEM BAM files as input. Previously published scaffolded avocado assemblies (Rendón-Anaya et al., 2019; Nath et al., 2022; Solares et al., 2023) were aligned to each scaffolded assembly using minimap2 v2.30 (Li, 2018) to verify assembly structure. The most contiguous assembly with the largest Nxsome and total size was retained as the final assembly for all downstream analyses. The Nxsome metric, the percentage of the expected genome size contained in the *x* largest scaffolds where *x* is the haploid chromosome count, follows the measure described in HapSolo (Solares et al., 2021) and the Gwen genome (Solares et al., 2023): scaffolds are sorted by size, and the largest 12 (for avocado) are summed and divided by the expected genome size of 896 Mb. Cumulative assembly length was computed for the contig and Omni-C scaffold assemblies with an in-house Python script (cumulativesum.py) by sorting sequences by descending length and adding them one at a time (Figure 1). The 12 largest scaffolds were assigned to chromosomes 1 to 12 by alignment to the Hass UoQ reference (GenBank GCA_029852735.1). Chromosome 8 was then curated manually as described below. The curated scaffolds were named by alignment to Unnamed Lowland telomere-to-telomere assembly with minimap2 v2.30 (-x asm5): each of the 12 chromosome-scale scaffolds placed 93.7 to 99.7% of its length on a single Unnamed Lowland chromosome, one to one, and took that chromosome number, and all other scaffolds were labeled unplaced. The resulting 12-chromosome FASTA (assembly v4) was used in all subsequent analyses of this study.

**Figure 1:**
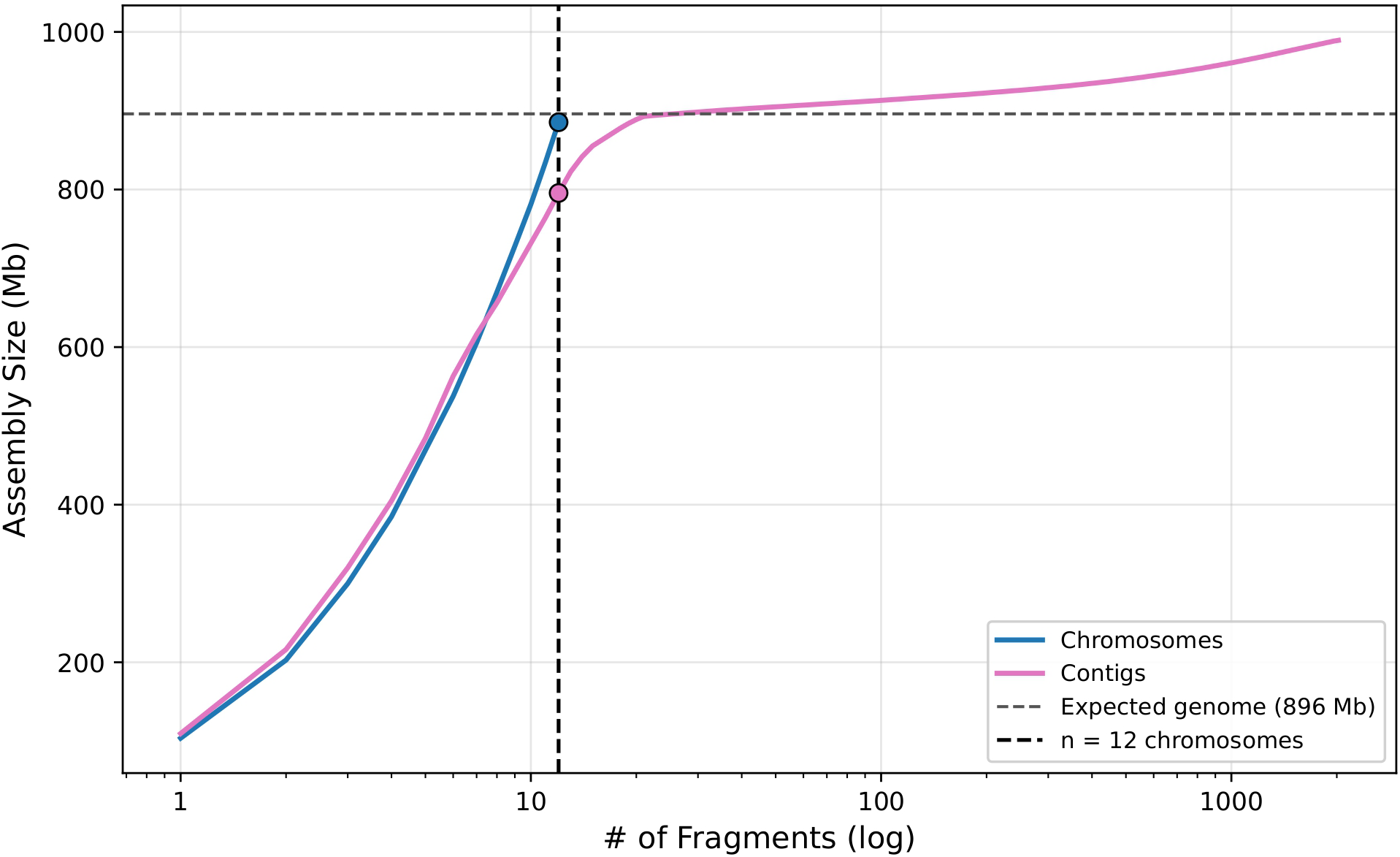
Cumulative length of the Thille genome assembly. Cumulative length of the 12 chromosomes (885.4 Mb) and of the 2,021 hifiasm contigs (989.3 Mb), largest first. The dashed lines mark the expected genome size (896 Mb) and 12 chromosomes.

### Manual Curation of Chromosome 8

YaHS did not join chromosome 8 into a single scaffold: one arm was carried by scaffold_12 (42.9 Mb, the first 42.85 Mb of the hifiasm contig ptg000004l) and the other was left as an unplaced 39.7 Mb scaffold (scaffold_13, the complete contig ptg000011l). The facing ends of the two scaffolds were aligned to each other with minimap2 v2.30 (-c -x asm5), and Thille HiFi reads were aligned to the complete YaHS assembly (minimap2 -ax map-hifi –secondary=no) and read depth was computed with SAMtools v1.24 depth -a and summarized in 1 Mb windows relative to the median of each scaffold. Following the manual curation route documented for YaHS (Zhou et al., 2023) and used in reference genome projects (Howe et al., 2021; Rhie et al., 2021), a curation contact map was generated from the YaHS output with juicer pre -a (YaHS v1.2.2) and juicer_tools v2.20.00 pre without matrix normalization (-n), since genome-wide normalization failed on the single assembly-mode pseudo-chromosome, and the join was edited in Juicebox Assembly Tools v2.17.00 (Durand et al., 2016; Dudchenko et al., 2018). The edited assembly was converted to AGP and FASTA with juicer post, and the conversion was verified by recovering every hifiasm contig segment, in its recorded orientation, exactly once from the curated FASTA.

### Assembly Quality Assessment

Assembly completeness was benchmarked using BUSCO v6.1 (Manni et al., 2021; Tegenfeldt et al., 2025) with the embryophyta_odb12.2 database for both the 12-chromosome assembly (genome mode with miniprot) and the predicted proteins (protein mode) (Figure 3B), and an in-house script was used to extract assembly statistics. Base-level accuracy and k-mer completeness were assessed with Merqury v1.4.1 and Meryl v1.4.1 (Rhie et al., 2020) at k = 21, chosen from the Merqury best_k.sh estimate of 19.8 for an 874 Mb genome at a collision rate of 0.001. A read k-mer database was built from all HiFi reads with meryl count, and merqury.sh was run against the 12-chromosome assembly with default parameters. Consensus quality (QV) was derived from k-mers present only in the assembly, k-mer completeness from the fraction of reliable read k-mers present in the assembly, and the copy-number spectrum from the number of distinct read k-mers at each multiplicity, stratified by their copy number in the assembly (Figure 3A).

### Genome Annotation

A de novo repeat library was built from the full Omni-C scaffolded assembly using RepeatModeler v2.0.9 (Flynn et al., 2020) with the -LTRStruct option. Transposable element (TE) annotation and soft masking of the 12-chromosome assembly were then performed using RepeatMasker v4.2.4 (Smit et al., 2013–2015) in sensitive mode with rmblastn v2.17.1+ against that library, and BEDTools v2.30.0 (Quinlan and Hall, 2010) was used to apply the RepeatMasker annotation as a soft mask. RepeatModeler and RepeatMasker were run in Singularity containers (Kurtzer et al., 2017). Gene annotation was performed by extracting peptide FASTA sequences using GffRead (Pertea and Pertea, 2020) from the Gwen genome (Solares et al., 2023) and the Hass genome (Nath et al., 2022). These peptide sequences were used as protein evidence for gene prediction on the soft-masked Thille genome using a deep learning approach, Tiberius v2.0.7 (Gabriel et al. 2024; https://github.com/Gaius-Augustus/Tiberius), run on a single NVIDIA RTX A6000 GPU. The fraction of the assembly assigned to each repeat class, and the fraction occupied by gene models, were summarized with an in-house script (Figure 2). Gene and TE density were computed per chromosome with an in-house Python script and summarized together with the centromere positions described below (Figure 5C).

**Figure 2:**
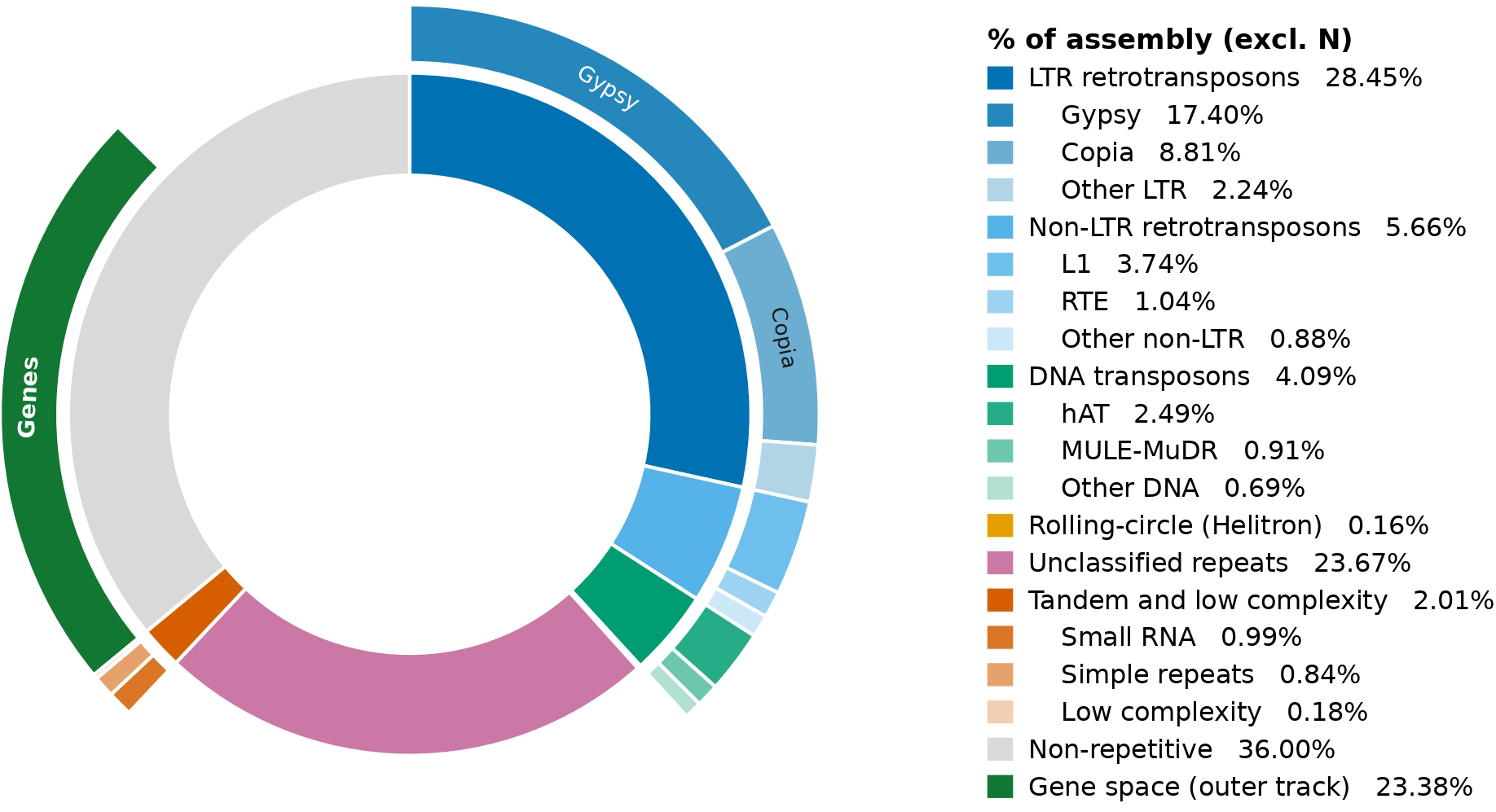
Genome annotation composition. Repeat classes (inner ring), their subfamilies (outer ring), and gene space (outer track) as percentages of the 885.4 Mb assembly (GC 39.7%), of which 64.0% is repetitive.

### Omni-C contact maps

Omni-C contacts were exported from the YaHS output with juicer pre (default minimum mapping quality of 10) using the AGP of the curated assembly, so that pairs were reported in chromosome coordinates, and pairs with both ends on chromosomes 1 to 12 were retained. Pairs were binned at 50 kb with cooler v0.10.4 (Abdennur and Mirny, 2020), aggregated to 100, 250, 500, and 1,000 kb with cooler zoomify, and matrix-balanced at each resolution by iterative correction (Imakaev et al., 2012) with the cooler defaults (Figure 4). A .hic file for viewing in Juicebox was built from the same pairs with juicer_tools v2.20.00 pre.

### Centromere and Telomere Identification

Centromeres were identified on each chromosome by combining four lines of evidence (Omni-C contacts, satellite repeats, telomeric repeats, and gene density) into candidate regions, which were then resolved by elimination with an in-house Python script (resolve_centromeres.py) using cooler v0.10.4 (Abdennur and Mirny, 2020), NumPy v1.26.4 (Harris et al., 2020), and SciPy v1.17.1 (Virtanen et al., 2020). Tandem repeats were identified on each chromosome with Tandem Repeats Finder (TRF) v4.10.0 (Benson, 1999) using the parameters recommended in the TRF documentation (https://github.com/Benson-Genomics-Lab/TRF; match 2, mismatch 7, indel 7, match probability 80, indel probability 10, minimum score 50), a maximum period of 2,000 bp, and the -h and -ngs options. Omni-C candidates were derived from the balanced chromosome 1 to 12 contact map, on the assumption that centromeres cluster in the nucleus and therefore share inter-chromosomal contacts (Varoquaux et al., 2015; Mascher et al., 2017). For each chromosome, every inter-chromosomal block was divided by its mean non-zero value and smoothed with a Gaussian filter (*σ* = 1 Mb; SciPy ndimage), and a contact profile was computed as the sum, over all other chromosomes, of the strongest contact made by each position anywhere on that chromosome. The whole chromosome was searched from coarse to fine resolution: local maxima of the profile at 1 Mb resolution that exceeded the median by at least half the difference between the maximum and the median were retained, and each was refined at 250 kb and then 100 kb resolution within 1 Mb on either side. A joint fit of all 12 centromere positions, following the approach of Centurion (Varoquaux et al., 2015) but implemented here with random initializations, was tested first and rejected, since 200 random starts converged to 22 distinct solutions and the highest-scoring solution placed every centromere at a chromosome end. Satellite candidates were clusters of TRF arrays with periods of at least 100 bp, merged when within 50 kb of each other and retained when at least 200 kb long. Overlapping candidates were merged, and a Omni-C peak was assigned to a candidate region when it fell within 1 Mb of it. The centromeric satellite family was chosen from the data rather than assumed, since centromeric satellites vary widely in sequence and monomer length among species (Melters et al., 2013; Naish et al., 2021), and the most abundant tandem repeat is typically centromeric (Melters et al., 2013). TRF periods of at least 100 bp were grouped into families of periods within 5% of each other, and the family with the most sequence within the gene-poorest 10% of 500 kb windows genome-wide was designated centromeric. Candidates on each chromosome were then eliminated in four ordered steps: (1) candidates within 2 Mb of a chromosome end carrying at least 1 kb of the plant telomeric repeat (TTTAGGG or CCCTAAA; Richards and Ausubel 1988) within 100 kb of the end were classified as telomeres; (2) candidates with less than 100 kb of the centromeric satellite family were removed, a step skipped when no candidate on the chromosome carried the family; (3) candidates with a gene density above the chromosome mean were removed, since centromeres are gene-poor and dominated by satellites and retrotransposons, as in Arabidopsis (Naish et al., 2021), a step skipped when it would remove every candidate; and (4) the surviving candidates were ranked by centromeric satellite content, with the Omni-C peak height as a tiebreaker. The centromere position was placed at the Omni-C peak when it lay within 1 Mb of the satellite cluster, and otherwise at the midpoint of the cluster. As a check, the ranking in step 4 was repeated with the Omni-C peak height as the first key. Each chromosome was classified from the ratio of its long arm to its short arm at the centromere position as metacentric (1.0 to 1.7), submetacentric (more than 1.7 to 3.0), acrocentric (more than 3.0 to 7.0), or telocentric (more than 7.0) (Levan et al., 1964) (Figure 5C).

### Comparative assembly analysis

Whole-genome alignments were generated with minimap2 v2.30 using the assembly-to-reference preset (-x asm5). For Thille versus Hass, the 14 largest Thille scaffolds were aligned to the complete Hass UoQ assembly (482 sequences, 912.9 Mb) with –secondary=no. For Thille versus Gwen, the Gwen primary assembly (Solares et al. 2023; 989 contigs and 1,032.1 Mb as counted in this study) was aligned to Thille chromosomes 1 to 12 with base-level alignment (-c). Gwen contigs were placed on the Thille chromosome contributing the most matched bases among alignments with a mapping quality of at least 20 and ordered by the match-weighted midpoint of their alignments; contigs with no qualifying alignment were assigned to an unplaced group. For display and strand assignment, alignment blocks of at least 10 kb were retained. Any Hass chromosome or Gwen contig whose matched bases in these blocks lay mostly on the minus strand was reverse-complemented, following the mummerplot –layout convention, so that blocks on the opposite strand from the rest of their sequence mark internal inversions (Supplementary Figure S2). Whole-genome sequence identity between Thille and Hass was estimated with ModDotPlot v0.9.9 (Sweeten et al., 2024) in static comparison mode (–compare-only) with default parameters (k = 21, 1,000 windows, delta 0.5, identity threshold 86%). Chromosomes 1 to 12 of each assembly were concatenated in numerical order with 100 kb of N between chromosomes. Hass chromosomes that aligned to Thille end to end on the opposite strand, identified by a negative Pearson correlation between Thille and Hass positions across same-chromosome windows of at least 95% identity and confirmed by the minimap2 strand assignment above, were reverse-complemented before concatenation; this reverse complementation reflects the strand chosen for each assembly rather than an inversion (Supplementary Figure S2).

### Additional Reference Assemblies

Two further chromosome-level avocado assemblies were included in the comparisons: the telomere-to-telomere assembly of a single Lowland avocado accession (Persea americana var. americana, commonly called West Indian; variety not reported; Unnamed Lowland in this study), whose two haplotypes both carry the recessive SDMYB allele found in Thille, so that it is predicted type B (see Results) (Yang et al. 2024; GenBank GCA_051132755.1, XTBG_Pame_1.0, 12 chromosomes, 841.6 Mb) and the assembly of the A-type cultivar Puebla, a predominantly Guatemalan hybrid with 82% Guatemalan ancestry (Chen et al. 2009; Groh et al. 2026; GenBank GCA_060308345.1, ASM6030834v1, WGS project JBTHJO000000000, BioProject PRJNA1392526; chromosomes CM191109.1 to CM191120.1), whose chromosomes were scaffolded with RagTag against the Lowland T2T assembly (Groh et al., 2026). The Unnamed Lowland assembly and its GenBank annotation (40,252 protein-coding genes) were downloaded from NCBI and checked against the supplied md5 check-sums, and its PacBio HiFi and ONT ultra-long reads were downloaded from Science Data Bank (doi:10.57760/sciencedb.07602, version V3) and checked for size and gzip integrity. Each assembly was aligned to Thille chromosomes 1 to 12 with minimap2 v2.30 (-x asm5; base-level alignment with -c for Unnamed Lowland), the Thille chromosome receiving the most matched bases was recorded for each query chromosome, and query chromosomes whose matched bases in blocks of at least 10 kb lay mostly on the minus strand were reverse-complemented, as for Hass. Whole-genome sequence identity between Thille and each of Unnamed Lowland and Puebla was estimated with ModDotPlot v0.9.9 using the settings applied to Hass.

### Structural Rearrangements

Structural rearrangements and large insertions and deletions between Thille and each chromosome-level assembly were identified with SyRI v1.7.0 (Goel et al., 2019). Following the SyRI documentation, query chromosomes were renamed chr1 to chr12 and placed on the same strand as Thille, and each query was aligned to Thille with minimap2 v2.30 (-ax asm5 –eqx); alignments were converted to BAM with SAMtools v1.24 and passed to SyRI (-F B). Thille served as the reference for Hass, Unnamed Lowland, and Puebla. For multi-genome visualization, two further SyRI comparisons were run with the same settings: Unnamed Lowland as reference with Thille as query, and Hass as reference with Puebla as query. SyRI annotations of at least 100 kb (inversions, translocations, duplications, copy gains and losses, deletions, highly diverged regions, and sequence without alignment in the other genome) were converted to BED format in the coordinates of each assembly, mapping reverse-complemented query coordinates back to the original GenBank sequences, and merged into a single table of events with the coordinates of each genome side by side (Supplementary Table S1). Events shared by the comparisons with Unnamed Lowland and Puebla were identified on Thille coordinates as events of the same class (inversion, translocation, duplication, deletion, insertion, copy gain, copy loss, or Thille-only sequence) of at least 50 bp with a reciprocal overlap of at least 50%, or, for insertions, which have no Thille span, with breakpoints within 100 bp; the same rule was used to ask whether Hass carries each shared event. Each event with a Thille interval was annotated with the Tiberius genes it overlaps, its TE fraction (union of RepeatMasker intervals), and its distance to the Thille centromere call. Rearrangements and unaligned sequence of at least 100 kb are shown in Figure 5A, and synteny across Unnamed Lowland, Thille, Hass, and Puebla assemblies in Figure 5B. Because the Gwen assembly is contig-level, it was not analyzed with SyRI. Instead, Thille intervals of at least 100 kb covered by no Gwen contig, or by two or more distinct Gwen contigs, were identified from primary alignments of at least 10 kb with a mapping quality of at least 20, and each interval was tested against Gwen CLR read depth on Thille (SAMtools v1.24 bedcov) relative to the median of 1 Mb windows on that chromosome. Intervals were classed with the coverage boundaries described under Raw Read Mapping as absent (at most 0.25 of the median), half (above 0.25 to 0.75), two copies (above 0.75 to 1.25), or more than two copies (above 1.25), which separates sequence missing from one or both Gwen haplotypes from sequence absent only from the Gwen assembly.

### Chromosome 3 and Chromosome 10 Loci

Two loci flagged by these comparisons, Thille chromosome 3 from 9.10 to 10.20 Mb and chromosome 10 from 11.56 to 13.60 Mb, were examined in Unnamed Lowland, Thille, and Hass. The corresponding sequence in each genome was taken as the interval between the SyRI syntenic blocks flanking the Thille locus. Tandem repeats in each interval and in a window extending at least 1.5 Mb on either side were identified with TRF v4.10.0 using the parameters described above. TEs in the Hass and Unnamed Lowland windows were annotated with RepeatMasker v4.2.4 using the Thille RepeatModeler library and the settings used for the Thille genome (-s -lib consensi.fa.classified), so that the three TE tracks are directly comparable, and TE classes were assigned from the library family names. Read depth across each window was taken from BEDTools genomecov -d output for every raw read alignment to that genome (Thille and Unnamed Lowland HiFi reads, and Gwen CLR reads), reduced to the mean depth of 10 kb bins and divided by the median depth of the chromosome over all positions. Candidate flowering genes from the top 1% Fst windows that fell within each window were located in each genome as described below.

### Read Support at Inversion Breakpoints

Every inversion longer than 100 bp in the SyRI comparisons of Thille with Hass (UoQ), Unnamed Lowland, and Puebla was evaluated with raw reads: 152 inversions (37, 52, and 63, respectively). Reads were aligned with minimap2 v2.30 (-ax map-hifi for HiFi reads and -ax map-pb for continuous long reads, –secondary=no) and sorted and indexed with SAMtools v1.23.1. Thille HiFi, Unnamed Lowland HiFi, and Gwen continuous long reads were aligned to the Thille, Hass (UoQ), and Unnamed Lowland chromosomes; because chromosome-scale scaffolds of the Hass (UoQ) assembly appeared to be misplaced in places, Puebla HiFi reads (SRR36605778; Groh et al. 2026), Thille HiFi, Unnamed Lowland HiFi, and Gwen continuous long reads were also aligned to the Puebla assembly (all 527 sequences), and Puebla HiFi reads to the Thille chromosomes. All alignments used the same settings. Query coordinates, which SyRI reports on chromosomes renamed and placed on the Thille strand, were converted back to the original assembly and strand; for all 152 inversions, the Thille interval aligned to the converted interval in the orientation expected for an inversion (minimap2 -x asm5). Each inversion was examined in windows of exactly 100 kb on each genome: one window centered on the inversion when it spanned at most 100 kb, and two windows centered on its breakpoints otherwise (18 inversions), for 169 Thille windows and 168 windows on the other assemblies. For every window and read set, a pileup of all reads was generated with SAMtools v1.24 mpileup (-aa -B -Q 0 -q 0 -s, with the reference sequence), without mapping or base quality filtering, so that poorly placed reads remained visible; depth was summarized in 500 bp bins for all reads and for reads with a mapping quality of at least 20 and expressed relative to the chromosome median depth of that read set (median of 30 randomly placed 10 kb windows). At each breakpoint, primary alignments with a mapping quality of at least 20 were classified with pysam v0.24.1 as spanning (aligned continuously across the breakpoint with at least 5 kb on both sides), clipped (alignment end within 1 kb of the breakpoint with a clip of at least 500 bp), or split (a clipped read whose supplementary alignment lies within 10 kb of the other breakpoint of the same inversion, on the same chromosome and the opposite strand, the signature of an inversion). Summed over the two breakpoints, a read set was scored as following the assembly (at least 3 spanning and fewer than 3 split reads), carrying the other orientation (at least 3 split and fewer than 3 spanning reads), carrying both orientations (at least 3 of each, as expected where the read donor is heterozygous for the inversion), or uninformative. Inversions shorter than the 5 kb flank can be crossed by a single read, so for these a spanning read indicates only that no split alignment was made. Below the read tracks, the breakpoint figures show candidate flowering genes, all gene models (Thille Tiberius models and the Hass (UoQ) and Unnamed Lowland GenBank annotations; no gene annotation is available for Puebla), and transposable elements, annotated in the Hass, Unnamed Lowland, and Puebla windows with RepeatMasker v4.2.4 and the Thille library as for the chromosome 3 and chromosome 10 loci.

### Thille-guided Scaffolds of Gwen and Hass (UoQ)

Because Thille is the mother of Gwen and a child of Hass, the Gwen contigs and the Hass (UoQ) assembly were also ordered and oriented on the Thille chromosomes with RagTag v2.1.1 scaffold (Alonge et al. 2022; minimap2 v2.30 -x asm5, 100 bp gaps of unknown size, no correction step, so no contig was broken). The Gwen primary contigs (989 contigs, 1,032.1 Mb) and alt contigs (the HapSolo secondary set, 4,133 contigs, 417.8 Mb; Solares et al. 2021; Solares et al. 2023) were scaffolded separately, and the 12 Hass (UoQ) chromosomes were first split at their 36 gaps into 48 contigs. Every input contig was recovered exactly from each output through its AGP, and completeness was assessed with BUSCO v6.1 in genome mode on chromosomes 1 to 12.

### Raw Read Mapping

Raw sequence reads were aligned with minimap2 v2.30 (–secondary=no), using the HiFi preset (-ax map-hifi) for Thille, Unnamed Lowland, and Puebla HiFi reads (Puebla: SRR36605778; Groh et al. 2026) and the CLR preset (-ax map-pb) for Gwen PacBio CLR subreads (Solares et al., 2023). Thille HiFi, Gwen CLR, and Unnamed Lowland HiFi reads were aligned to the 12 Thille chromosomes (curated assembly, v4) and to the 12 Hass (UoQ) chromosomes, Puebla HiFi reads to the Thille chromosomes, and Thille HiFi reads also to all Thille Omni-C scaffolds and all Hass (UoQ) sequences (Table 3); the alignments to the Puebla assembly are described under Read Support at Inversion Breakpoints. The Hass HiFi reads available from NCBI were a fraction of those reported (Nath et al., 2022) and were not used. SAM output was sorted, indexed, and split into per-chromosome BAM files that retained the full sequence header with SAMtools v1.23.1 (sorting and splitting) and v1.24 (final sorting and indexing). Per-position read coverage was computed for each alignment from the sorted, indexed BAM files using BEDTools v2.30.0 genomecov with the -d option, which reports depth at every position in 1-based coordinates, including positions with zero depth. For comparison, BEDTools coverage was also run with the -d option using the sorted BAM and a BED file spanning each chromosome, generated from the assembly sequence lengths using awk with the columns sequence ID, start, and end. The genomecov output was used for all subsequent classification and plotting. Before use, each chromosome was checked to contain every position from 1 to its full length, in order and without gaps. Coverage was classified at single-nucleotide resolution relative to the median depth of the source chromosome. For chromosome c, the median depth Mc was computed over all positions of the chromosome, including zero-depth positions, and truncated to an integer. Each position with depth d was assigned to one of five classes: low (0 *≤* d *≤* 0.25 Mc), half (0.25 Mc < d *≤* 0.75 Mc), medium (0.75 Mc < d *≤* 1.25 Mc), high (1.25 Mc < d *≤* 1.5 Mc), and outlier (d > 1.5 Mc). The class boundaries lie midway between the depth ratios expected for 0, 0.5, and 1 copies of a haplotype relative to the chromosome median, so that the low and half classes mark sequence absent from, or present on only one haplotype of, the aligned cultivar, and the high and outlier classes mark sequence present at more than the diploid copy number, such as duplications, copy number variants, or collapsed repeats. Coverage was examined at two scales. Genome-wide, the per-position genomecov output was read per chromosome into a pandas data frame with three fields (sequence ID, position, and coverage) and compared with the chromosome median (Figure 5C). At each candidate locus, the classes were evaluated for every position in a fixed 100 kb window centered on the gene model, with the window start rounded to the nearest kilobase so that every gene was examined at the same scale; alignments to Thille and to Hass were each evaluated in the coordinates of their own reference (Figure 7; Supplementary Figures S4 and S13). For each candidate gene, the fraction of the gene-model span in each class was recorded for each alignment.

### Allelic Distribution Graphs of Candidate Genes

Alleles were reconstructed for each candidate gene with a Thille gene model, for the controls KCS11 and MIK2 on chromosome 5, and for the pigment genes SGRL and LCYB. For each gene, the coding sequence (CDS) exons of the Tiberius model (first transcript) were taken in transcript order, and the analysis window was the gene span with 10 kb of flanking sequence on each side. Read-based haplotypes were obtained for Thille HiFi, Unnamed Lowland HiFi, and Gwen CLR reads aligned to Thille with minimap2 v2.30 (primary alignments; SAMtools v1.21). Heterozygous sites were positions within the gene span and 2 kb of flanking sequence covered by at least 10 reads at which the second most frequent allele was carried by at least 25% of reads; for Gwen CLR reads, random sequencing error was masked by restricting heterozygous sites to substitutions at which the two most frequent bases accounted for at least 80% of non-gap reads and the second base for at least 30%. Reads covering at least three heterozygous sites were assigned to two haplotypes by two-means clustering of their alleles (Hamming distance over shared sites), seeded by the two most different reads sharing at least 5 sites (HiFi) or 20 sites (CLR); for SDMYB, this assignment agreed, read for read, with one seeded by the coding substitutions that distinguish the two Gwen alleles (88 of 88 reads). At every position of the window, the allele of each haplotype was the allele carried by at least 50% of its reads (at least 3 reads, gaps excluded), or, when no allele reached 50%, its single most frequent allele. A read set in which no two reads shared enough heterozygous sites was treated as one diploid pool: the most frequent allele was assigned to the first haplotype and the second most frequent to the second haplotype when it was carried by at least 25% of reads, and otherwise both haplotypes received the majority allele, whose observed frequency was reported. Alleles were never inferred from another cultivar or from the reference, and positions without at least 3 reads of a haplotype were recorded as missing. Insertions and deletions of at least 2 bp were clustered across reads by type, position (within 10 bp), and length (within 2 bp or 10%), retained when carried by at least 5 reads and by at least 25% of all spanning reads or 50% of the reads of one haplotype, and assigned to haplotypes by the same rules. The CDS of each haplotype was rebuilt from the majority base of its reads at each coding position, including indels supported by most of its reads, and coding positions without read support were recorded as unknown. Assembled haplotypes were taken from the haplotype-resolved assemblies of Puebla (GenBank GCA_060308345.1 and GCA_060308325.1; Groh et al. 2026) and from the principal and alternate pseudohaplotypes of Hass (MH2022; GCA_031208685.1 and GCA_031208655.1). Each Thille window was mapped to each assembly with minimap2 (-x asm20 –secondary=no); alignments on the homologous chromosome were chained around the block with the most matching bases, and the assembly region spanning the chain with 2 kb of flanks was aligned back to the Thille window (minimap2 -c –cs=long -x asm20). A window was considered placed when the aligned blocks covered at least 80% of it. Substitutions, insertions and deletions of at least 2 bp, and unaligned stretches between consecutive blocks were recorded as variants, and the CDS was rebuilt from the aligned bases, including 1 bp indels. The single-sequence Hass (UoQ) assembly was compared with both pseudohaplotypes over every window. Alleles were named with superscript letters: GENE^a^ is the Thille reference CDS, and every distinct CDS received the next letter, so that haplotypes share a letter only when their CDS are identical and any coding difference defines a new allele; the SDMYB alleles are also described as recessive (SDMYB^a^) and dominant (SDMYB^b^), following Groh et al. (2026). Allele frequency (AF) within each read set was the fraction of reads carrying the allele of a haplotype, pooled over CDS positions at which its two haplotypes differed and counting only the two most frequent alleles at each position; intronic and flanking variation was not used. Two phased haplotypes with identical CDS were reported at 100%, an unphased read set with a single allele at its observed majority-allele frequency, and assembled haplotypes at 50% or 100%. To test for the dominant SDMYB allele without read alignment, all 21-mers overlapping ten coding substitutions that distinguish the two alleles were counted exactly in the raw reads of each read set. Read sets in which one phased haplotype carried 60 to 80% of the reads were examined for structural causes: the depth of all reads over the gene was compared with that over flanks 2 to 6 kb away (an additional copy on one of two haplotypes is expected near 1.5 times the flank depth, and the loss of one haplotype near 0.5), and soft- or hard-clipped alignments of at least 200 bp were clustered within 100 bp, with clusters of at least 3 reads reported (Figure 6; Supplementary Figure S3).

### Candidate Flowering Gene Identification and Alignment

Candidate flowering genes were extracted from Gwen genes located within the top 1% of 20 kb windows ranked by Fst between flowering types A and B (Solares et al., 2023). Two Fst analyses of the same data were combined: the top 1% windows of the contig-level analysis (517 of 51,701 windows) and of the scaffold-level analysis (508 of 50,775 windows) were each intersected with the Gwen gene annotation, and the union of the two gene sets, 944 genes (246 in both sets, 463 in the contig set only, and 235 in the scaffold set only), was extracted for annotation. SwissProt and UniRef90 matches of these genes were collected from blastx and blastp searches of the contig-level and scaffold-level gene sets (BLAST+ v2.2.30; e-value at most 1e-5) and from Blast2GO. The SwissProt and UniRef90 matches reported for these genes in the Gwen study were re-annotated against UniProtKB release 2026_03 (UniProt Consortium, 2025) through the UniProt REST API (Ahmad et al., 2025). Flowering-associated genes were then identified from protein names, functional descriptions, Gene Ontology (GO) biological process terms, and membership in the FLOR-ID flowering-time gene database (Bouché et al., 2016). The keyword search was performed with a large language model (Claude Opus 5.5, Anthropic) following author-defined instructions for validation and verification, and each gene identified by the search was verified manually. Of the 944 genes, 36 Gwen gene models met at least one criterion: membership in FLOR-ID (7), a flowering-related GO biological process term (21), or a flowering-related protein name or function keyword (29). After manual checking and filtering, 24 gene models (20 genes) were retained as candidate flowering genes and 12 were set aside as indirectly annotated; four of the latter (FBD/RNI-like, RER3, RPT3, and PBL34) were added after review, for a final set of 28 gene models representing 24 genes (Table 4). Gene sequences were extracted from the Gwen genome in FASTA format with BEDTools v2.29.2 getfasta (-s) (Quinlan and Hall, 2010) using the Gwen gene annotation as input, aligned to the Thille and Hass genomes with minimap2 v2.30 (Li, 2018) using the spliced alignment preset (-ax splice), converted to sorted, indexed BAM files with SAMtools v1.24 (Danecek et al., 2021) as described above, and converted to BED format with bamToBed from BEDTools v2.29.2. The Hass reference was the University of Queensland (UoQ) assembly of cultivar Hass (GenBank GCA_029852735.1; chromosomes CM056809.1 to CM056820.1), and the Thille reference was the Omni-C scaffolded Thille assembly (12 chromosomes, 859,826,806 bp). In addition to the spliced alignments, every candidate flowering gene, including SDMYB, SPA1, SPA1-like, and the two loci listed as EOBI in the Gwen study (Tables 4 and 6), was located in the Hass (UoQ) genome with BLAST+ v2.2.30 (Camacho et al., 2009). Gene sequences were searched with blastn (-task megablast, -evalue 1e-20), and translated protein sequences were searched with tblastn (-evalue 1e-5) as an independent protein-level check. Since a small -max_target_seqs value can cause BLAST+ to report hits other than the highest-scoring ones (Shah et al. 2019; see also the response by Madden et al. 2019), -max_target_seqs was set to 50, exceeding the number of chromosome sequences in each target assembly, and the number of high-scoring segment pairs (HSPs) per subject was left unrestricted (-max_hsps 0) so that multi-exon genes were recovered as complete loci. Output was written in tabular format with query and subject lengths appended (-outfmt “6 qseqid sseqid pident length mismatch gapopen qstart qend sstart send evalue bitscore qlen slen”). For each gene, HSPs on the same chromosome and strand were grouped into loci, the locus with the highest summed bitscore was retained, and its coordinates were converted to BED format with custom scripts. Each locus was then transferred to the Thille assembly as three sequences extracted from the Hass genome with BEDTools getfasta: the 1 kb upstream flank, the gene, and the 1 kb downstream flank. Each sequence was aligned independently to Thille with minimap2 (-a), and a locus was accepted only when the gene and both flanking alignments tiled contiguously on the same chromosome and strand. Placements for three genes (PAR1, MIK2 on chromosome 12, and KCS11) were corrected manually after these checks. One gene model per candidate was then assigned in each assembly. In Thille, the Tiberius (Gabriel et al., 2024) gene model overlapping the placed locus whose protein best matched the SwissProt entry named for the gene was selected, preferring entries named for the gene family; this also separated Gwen models that spanned more than one Thille gene (for example, EMF1 Chr3.ver1.g154150, which covers a GOGC5 golgin and EMF1). In Hass, where no UoQ protein models were used for selection, the chosen Thille protein was searched against the span of each overlapping UoQ gene model with tblastn, and the model with the highest bitscore was retained; the largest overlap was used when no protein match was found. Each selected Thille protein was aligned with blastp to the SwissProt entries (UniProtKB release 2026_03) named in the gene’s evidence. The resulting BED files report, for each gene and assembly, the full span of the selected gene model (BED6, 0-based). The selected Hass models were then checked against the UoQ protein set (GenBank GCA_029852735.1; 42,442 proteins, one per gene model): the protein of each selected UoQ model was aligned with blastp (BLAST+ v2.2.30, -evalue 1e-5) to the SwissProt entry named for the gene and to the selected Thille protein, and each selected Thille protein was searched against the complete UoQ proteome (blastp, -evalue 1e-10) to test whether another UoQ protein was a closer match.

### Fst Rank of Candidate Genes

Type A versus type B Fst was available from two analyses of the same resequencing data (Solares et al. 2023; top 1% gene sets at https://github.com/GautLab/avo_ref_paper), and the window counts, ranks, and thresholds used here were computed in this study from those windows: 20 kb windows on the HapSolo primary contigs of Gwen (51,701 windows) and 20 kb windows on the Gwen ver1 scaffolds (50,775 windows). In each analysis, windows were ranked by mean Fst (the MEAN_FST column of the windowed output), and the top 1% comprised the 517 contig and 508 scaffold windows of the Gwen study, which are the highest-ranked windows by this measure in each analysis. For every candidate gene, the best rank among the windows overlapping its Gwen locus was recorded separately for each analysis, using contig coordinates for the contig analysis and ver1 scaffold coordinates for the scaffold analysis, and a gene was counted as a top 1% Fst gene when either rank fell within the top 1%. Both ranks are reported for every gene (Tables 4 to 6). FLOR-ID membership was assigned by matching the gene symbol of every Arabidopsis SwissProt hit of each of the 944 genes to the gene names of the 306 FLOR-ID flowering-time genes (Bouché et al., 2016). The 12 gene models set aside as indirectly annotated, whose association with flowering rests on GO or Blast2GO terms for photoperiodism, gibberellin response, blue light signaling, or meristem identity, or on protein-name keywords, were placed on the Hass (UoQ) and Thille assemblies, and one gene model was selected in each, with the procedure described above. The eight not added to the candidate genes are listed in Supplementary Table S2.

### TE and SV Density at the Candidate Clusters

For the two clusters of candidate loci (chromosome 3, 73,340,678 to 91,615,352; chromosome 5, 19,577,582 to 30,337,161), TE density was computed as the fraction of nucleotides covered by the RepeatMasker TE annotation (each nucleotide counted once, simple repeats and low-complexity sequence excluded), and SV density as the number of SyRI events of at least 50 bp per Mb in the comparisons of Thille with Hass (UoQ), Unnamed Lowland, and Puebla, in each cluster and in flanks of 100 kb, 1 Mb, and 5 Mb on both sides. Each value was ranked against all non-overlapping windows of the same length on chromosomes 1 to 12 with a custom Python script; TE bp were checked with BEDTools merge and coverage.

### Placement of Puebla in the Gwen-Study PCA

Puebla HiFi reads were aligned to the Gwen primary contigs with minimap2 v2.30 (-ax map-hifi –secondary=no) and genotyped with BCFtools v1.24 (call -m -C alleles; depth of at least 5) at the 34,472,447 SNPs of the principal component analysis (PCA) of 30 resequenced accessions in the Gwen study (Solares et al., 2023). Puebla was projected onto the published components without recomputing them, with PLINK v2.0.0-a.7.8 (–score with the panel allele weights and allele frequencies, variance-standardized), and the projected scores were placed on the published axes by a linear fit per component on the 30 panel accessions (|r| of at least 0.996). Thille HiFi reads, whose Illumina genotypes are part of the panel, were processed the same way as a control. The projection was repeated on the two PCAs of the Gwen study computed on the Gwen scaffolds (29 and 30 accessions; 25,635,186 and 26,268,931 SNPs). Allele detail and coverage figures were produced for the ten genes recovered by the combined screen (FBD/RNI-like, RER3, RPT3, PBL34, Ole e 10-like, LIP1, ZEP, CIGR2, the CCCH zinc finger, and CTR1) with the procedures used for the other candidate genes. Their Thille and Hass (UoQ) placements were added to the candidate gene BED files, each Thille gene with its 1 kb flanks was placed on the Unnamed Lowland assembly with minimap2 v2.30 as described above, haplotype variants were called from Thille HiFi, Unnamed Lowland HiFi, and Gwen CLR reads and from the Puebla and Hass (MH2022) haplotype assemblies, and read coverage was plotted in 100 kb windows on the Thille and Unnamed Lowland assemblies (Supplementary Figures S5 and S6). For RER3 and the Ole e 10-like gene, which have no Tiberius model at their Thille locus, the Gwen protein and its SwissProt match (RER3_ARATH and ALL10_OLEEU) were searched with tblastn (BLAST+ v2.2.30, e-value at most 1e-5) against each placed locus in Thille, Unnamed Lowland, Hass (UoQ), the two Hass (MH2022) haplotypes, and the two Puebla haplotypes.

### Flowering-Type Allele Classes

To identify alleles that follow flowering type, the haplotypes of each gene were compared under the model established for SDMYB, in which type A is dominant (Groh et al., 2026): B-type individuals (Thille and Unnamed Lowland) are homozygous for one allele, and A-type individuals (Gwen, Hass, and Puebla) carry that allele on one haplotype and a different allele on the other. Two tests were applied to each of the 39 genes with allele detail figures. First, a variant site was counted as following the pattern when all four B haplotypes shared the same state and each of the three A individuals carried the variant on exactly one of its two haplotypes; carried states were the per-haplotype calls of the allele analysis; a haplotype without a call but with read coverage (at least 3 reads) or an aligned assembly block at the site was scored as not carrying the variant, and sites at which any of the ten haplotypes lacked data were not scored rather than assigned the Thille reference base. Second, whole alleles were compared with the same rule, once as identical coding sequences and once as identical proteins, so that synonymous differences were ignored. SDMYB served as the positive control. Because the Hass HiFi reads available from NCBI were a fraction of those reported and the single-sequence Hass (UoQ) assembly matches one Hass haplotype or the other from locus to locus, Hass haplotypes of the flowering genes were taken from the two phased Hass (MH2022) pseudohaplotypes, and Gwen and Puebla were used in place of Hass for read-level evidence. Although not ideal, this substitution was made only for flowering genes and only where a Gwen or Puebla haplotype carries the same allele as a Hass haplotype: an identical CDS, or the same protein with synonymous differences only, reported with an asterisk.

### Gene Naming Conventions via SwissProt homology

Thille gene models without a curated SwissProt entry, including all members of the tandem arrays described below, were named by homology. The Tiberius proteins of 110 Thille gene models (tandem-array members, single-model tandem groups, and the Thille models of the candidate flowering genes) were searched with blastp (BLAST+ v2.16.0; -evalue 1e-5, -max_target_seqs 25) against UniProtKB/Swiss-Prot release 2026_03 (575,748 entries). A hit was accepted when its e-value was at most 1e-10, its identity at least 30%, and its best HSP covered at least 50% of both the query and the subject protein. Each model was named after the gene symbol of its highest-scoring accepted hit, with the suffix “-like”. A model was flagged as having a predicted flowering function when its accepted hit carried the UniProt keyword Flowering (KW-0287), or a GO biological process term equal to or descending from flower development (GO:0009908), photoperiodism, flowering (GO:0048573), vernalization response (GO:0010048), vegetative to reproductive phase transition of meristem (GO:0010228), or regulation of flower development (GO:0009909), with descendants taken through is_a and part_of relations in go-basic.obo (release 2026-07-26). The function of each model was taken from the UniProt function annotation of its accepted hit.

### Tandem-copy Evaluation

Tandem copies of genes within the top 1% of Fst windows were evaluated in the Thille assembly: 386 copies in 59 groups (296 copies in 48 groups in Hass). Copies shorter than 1 kb were flagged as unlikely to represent complete gene copies. Each copy was intersected with the Tiberius gene annotation (36,349 genes; longest transcript per gene) and with the RepeatMasker annotation of transposable elements (TEs; 1,448,929 intervals), and the fraction of each copy covered by TEs was computed from the union of overlapping TE intervals. Groups whose copies overlapped no gene model and were each at least 95% covered by TEs were classified as TEs of interest and evaluated separately from the gene arrays. The Tiberius gene models overlapping the copies of a group were treated as candidate members of a tandem array. For every pair of members, three sequences were aligned globally in gene orientation: the full gene region extracted from the Thille genome with SAMtools v1.24 faidx, the concatenated exons of the longest transcript, and the Tiberius protein. DNA sequences were aligned with EMBOSS v6.6.0 stretcher (EDNAFULL, gap open 16, gap extension 4) and proteins with EMBOSS needle (EBLOSUM62, gap open 10, gap extension 0.5) (Rice et al., 2000), run from a Singularity (Kurtzer et al., 2017) container built for this study. Before alignment, the exon sequence of every member was confirmed to translate exactly to its Tiberius protein. Identity was calculated over alignment columns in which both sequences carried a residue, excluding end gaps, and coverage was calculated as the fraction of the shorter sequence aligned. A group was considered a supported tandem array when at least two of its gene models shared at least 70% exon identity, and groups sharing gene models were merged into a single array. Supported arrays were then evaluated manually, and four were retained for further analysis: CCR1 on chromosome 2, KCS11 on chromosome 6, the leucine-rich repeat receptor kinase array containing MIK2 on chromosome 12, and the interleaved SCPDL and HST arrays on chromosome 3. For each member of a retained array, the fraction of the gene span and of the coding sequence covered by the union of overlapping RepeatMasker TE intervals was computed, and members with at least half of their coding sequence covered were considered TE-like. Coverage across each retained array was examined in the smallest of six fixed window sizes (100 kb, 250 kb, 500 kb, 1 Mb, 2.5 Mb, and 10 Mb) that contained the entire array, so that arrays within a size class were examined at the same scale (Supplementary Figure S14). Tandem repeats were then manually inspected and several candidates were removed after visual inspection by E.S.

### Visualizations

Locus-level coverage plots were drawn with pandas v2.1.4, NumPy v1.26.4, matplotlib v3.6.3, and seaborn v0.13.2 (McKinney, 2010; Harris et al., 2020; Hunter, 2007; Waskom, 2021), and the Omni-C maps with cooler and matplotlib v3.8.4. Omni-C contact maps used a single-hue sequential color scale, with unbalanced or empty bins shown in white. The genome-wide map (500 kb bins) shows raw and balanced contacts on a linear scale from 0 to the 99.5th percentile of non-zero bins; per-chromosome maps (100 kb bins, raw contacts) use a logarithmic scale set per chromosome, with the lower limit at the median count of bins at least 10 Mb from the diagonal and the upper limit at the 95th percentile of bins within 1 Mb of the diagonal. Whole-genome alignments were drawn as MUMmer-style dot plots with each genome laid end to end in chromosome order and chromosome boundaries marked. ModDotPlot window identities were redrawn using the ModDot-Plot Spectral palette in 11 bins from 86 to 100% identity, with chromosome boundaries marked and reverse-complemented Hass chromosomes labeled “(rc)”. The karyotype (Figure 5C) was drawn with ChromoPlot, an in-house Python package built on matplotlib v3.8.4, with each chromosome drawn to scale, gene models and repeats painted along its length, the centromere drawn as a circle at the centromere position joining the two arms, the centromeric region shaded, and telomeres drawn as semicircular ends. Candidate genes were marked on the karyotype by small dots along the gene-side edge of each chromosome, at the midpoint of each Thille gene model or aligned locus, with one row of dots for each set a gene belongs to: top 1% Fst genes in vermillion, coverage-selected flowering genes in blue, and FLOR-ID genes in orange. The three colors were taken from a colorblind-safe palette and checked for separation under simulated protanopia, deuteranopia, and tritanopia (minimum pairwise OKLab difference of 13.1), and each dot was outlined in black so that the orange dots remained visible against the white background. Structural variants were visualized in two ways. Ribbon plots of SyRI annotations were drawn with plotsr v1.1.5 (Goel and Schneeberger, 2022) for each pairwise comparison and as chained comparisons of Unnamed Lowland, Thille, and Hass, of Unnamed Lowland, Thille, and Puebla, and of all four genomes, with chromosomes in numerical order and rearrangements of at least 100 kb shown (Figure 5B); zoomed four-genome ribbons were drawn for the chromosome 3 and chromosome 10 loci with plotsr –reg (Supplementary Figure S10). Whole-genome dot plots of the SyRI alignments were redrawn with an in-house script, marking each inversion, translocation, duplication, deletion, insertion, and copy-number change of at least 100 kb with an open circle at its position in both genomes, sized by event length, and marking unaligned sequence of at least 100 kb as strips along the axis of the genome that carries it (Figure 5A). The chromosome 3 and chromosome 10 loci were drawn for each genome as a chromosome ideogram with the locus marked, followed by a track of SyRI annotations of at least 50 kb labeled by type, gene, TE, and tandem repeat tracks, and one read-depth panel per alignment, with the candidate flowering genes highlighted and named in the gene track (Supplementary Figure S9). In the genome-wide coverage plots, positions with coverage above 1.5 times the chromosome median were drawn at that value and highlighted, so that high-depth repeats did not compress the scale. The remaining positions were smoothed for display only by taking the median coverage within non-overlapping 30 kb windows, and the chromosome median was drawn as a reference line. Gene positions from the Gwen gene BED file were overlaid, as points for the full candidate set and as vertical lines for the selected loci. In the locus-level plots (plot_gene_stacks.py), each position was drawn as a single point colored by its coverage class, on a depth axis from -5 to 125 shared by all panels; positions with depth above 120 were drawn at 122. Dashed reference lines mark 0, the four class boundaries, and 120, and a solid line marks the chromosome median. Alignments to Thille were drawn above alignments to Hass (UoQ), and the gene model was shaded in every panel. Beneath each group of coverage panels, gene models (Thille, Tiberius exon and CDS structure of the longest transcript; Hass, UoQ gene spans from the GenBank annotation), TEs (Thille only), and tandem copies were drawn on the same horizontal scale. In the tandem array figures (Supplementary Figure S14), the gene models of the array members were drawn with their exon and CDS structure in a separate copy track, in pink, or in pink for SCPDL and fuchsia for HST where the two arrays interleave, above a track of the other gene models in the window and the TE track, and the member spans were shaded in the coverage panels, which show alignments to Thille only. For windows larger than 100 kb, the points were rasterized at 600 dpi within otherwise vector PDF files. All locus plots were written as PDF and 600 dpi PNG files. Allele matrices show each gene window drawn 5*^′^* to 3*^′^* with its CDS exons, and below it one column per variant carried by at least one haplotype, grouped as 5*^′^* flank, numbered exons and introns, and 3*^′^* flank. Coding columns were drawn wider than noncoding columns so that their amino-acid labels remain legible, and columns were colored by variant type (amino-acid changing, synonymous, or noncoding substitution; deletion or insertion of at least 2 bp). Rows were ordered Gwen, Thille, Hass (MH2022), Puebla, and Unnamed Lowland; positions without data for a haplotype were hatched rather than drawn as the reference allele, and the allele letter and AF of each haplotype were printed at the end of its row. Coverage figures in the main text show alignments to the Thille assembly only, with the gene labels and headers of the allele matrices. Coverage on the Hass (UoQ) assembly is shown for SDMYB as a supplementary example (Supplementary Figure S7), because the Hass HiFi reads available from NCBI were a fraction of those reported (Nath et al., 2022), so Hass reads could not be shown on their own assembly. Genome-wide coverage was summarized as the median depth of non-overlapping 100 kb bins divided by the chromosome median for each read set on Thille (Supplementary Figure S8); the bins were used for display only.

## Results

### Genome Assembly

Sequencing yielded 72.2 Gb of PacBio HiFi reads (81x coverage of the expected 1C genome size of 896 Mb; Arumuganathan and Earle 1991) from four SMRT cells, with a mean read quality of Q27.8 and a read N50 of 14.2 kb. Assembly with hifiasm produced 989.0 Mb in 2,021 contigs, with a contig N50 of 78.9 Mb, an NG50 of 79.4 Mb, a largest contig of 109.5 Mb, and an Nxsome of 89% (795.6 Mb in the 12 largest contigs). Omni-C scaffolding with YaHS reduced the number of sequences by 26% to 1,491 scaffolds totaling 989.4 Mb, with an N50 of 67.8 Mb, an NG50 of 84.5 Mb, a largest scaffold of 103.7 Mb, and an Nxsome of 96% (Supplementary Figure S1). After manual curation of chromosome 8, the assembly comprised 1,494 scaffolds totaling 989.4 Mb, with an N50 of 68.4 Mb and an Nxsome of 98.8% (Table 1). Figure 1 plots the cumulative assembly length against the number of sequences for the contig and scaffold assemblies, and it illustrates two points. First, Omni-C scaffolding and curation placed an additional 89.8 Mb into the 12 largest sequences (885.4 Mb versus 795.6 Mb). Second, both curves flatten beyond 12 sequences, so the remaining sequences add little length relative to the expected genome size. The 12 largest scaffolds form the final assembly (v4), totaling 885.4 Mb (885,358,400 bp), with chromosome lengths from 51.1 Mb (chromosome 10) to 103.7 Mb (chromosome 2) (Figure 5C). YaHS did not join chromosome 8: one arm was carried by the 12th largest scaffold (scaffold_12, 42.9 Mb) and the other by the 13th (scaffold_13, 39.7 Mb), which aligns to Hass (UoQ) chromosome 8 from 0 to 30.4 Mb, whereas scaffold_12 aligns from 25.0 to 65.5 Mb. The two scaffolds shared a 14 Mb region (scaffold_13 25.7 to 39.7 Mb and scaffold_12 0 to 14 Mb, 95.5 to 98.2% identity) in which HiFi read depth on the complete YaHS assembly was 0.49 to 0.86 of the scaffold median on both copies, compared with 1.03 to 1.15 outside it, the signature of a false heterotype duplication in which the two haplotypes of a region are placed in different contigs (Rhie et al., 2021). In the curation contact map, the duplication appeared as a line parallel to the diagonal between the two scaffolds, interrupted where it crosses the chromosome 8 centromere. Scaffold_13 was placed before scaffold_12 in its original orientation, the duplicated 14,193,090 bp of scaffold_13 (25,531,495 to 39,724,584) were moved to the unplaced sequences, and the two scaffolds were joined with a 100 bp gap, giving a chromosome 8 of 68.4 Mb (68,384,694 bp); the other 11 chromosomes are identical to the uncurated assembly. GC content was 40.24% in both the contig and scaffold assemblies and 39.74% in the final 12-chromosome assembly. Of 77,450,874 Omni-C read pairs, 73,744,029 (95.2%) had both ends on chromosomes 1 to 12, and 77.99% of these were intrachromosomal (Table 1).

**Table 1:**
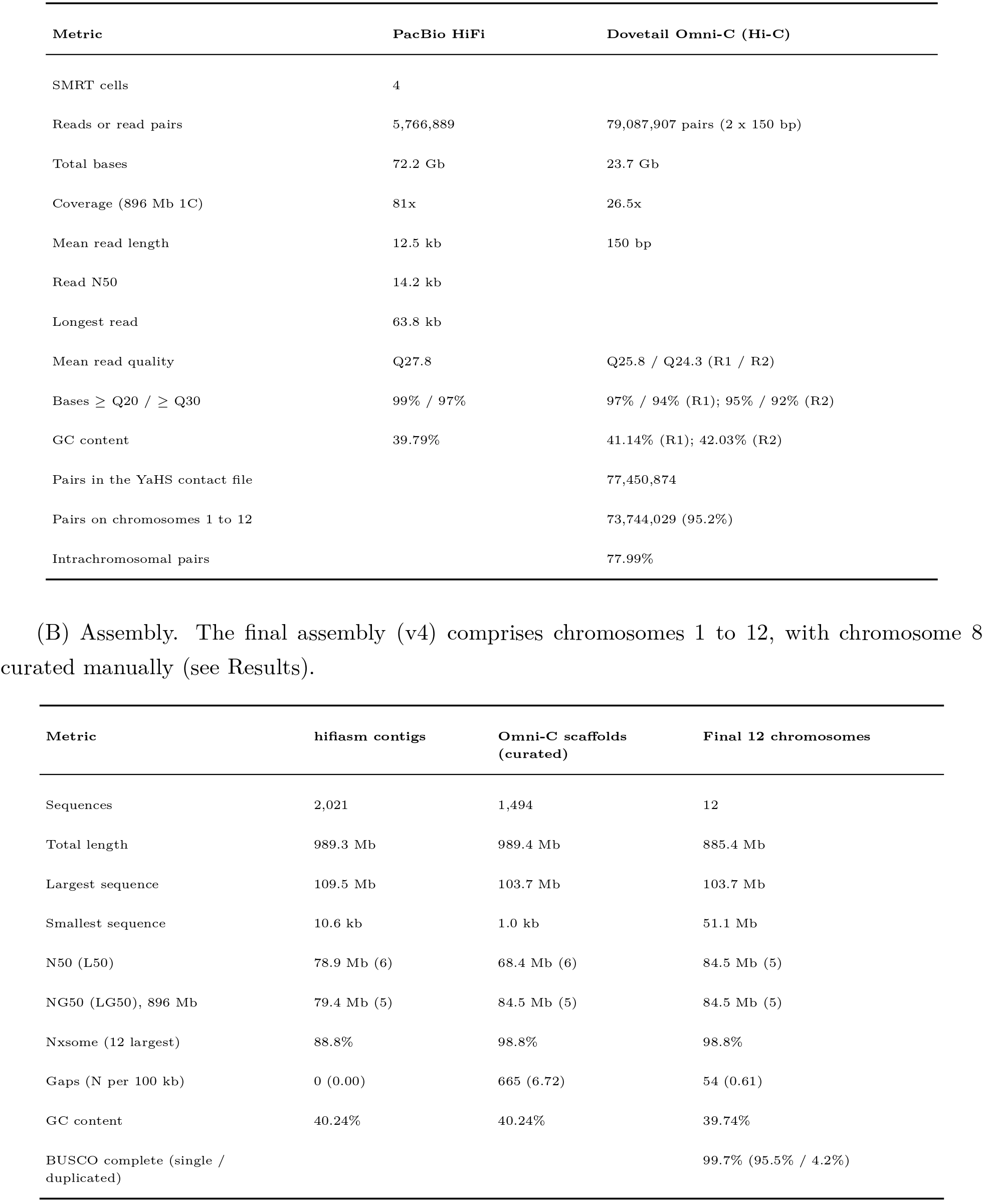
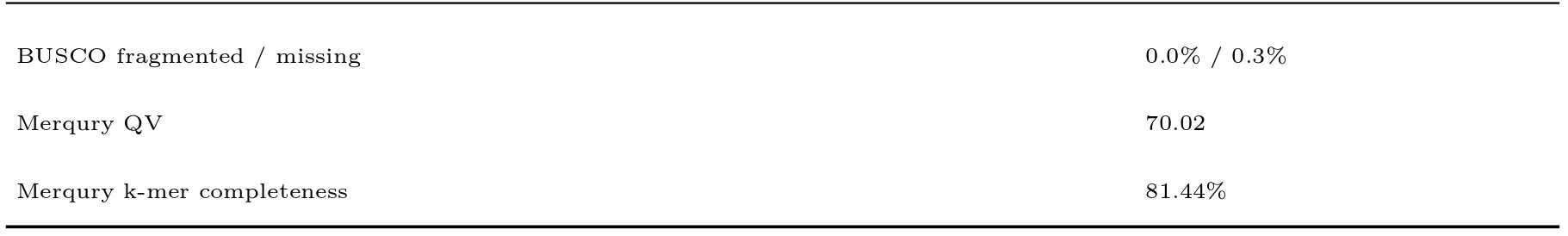
Genome Metrics. Sequencing and assembly metrics for the Thille genome. (A) Sequencing data. (B) Assembly. The final assembly (v4) comprises chromosomes 1 to 12, with chromosome 8 curated manually (see Results).

### Assembly Quality and Completeness

Of the 1,658 embryophyta_odb12.2 BUSCO genes, 1,653 (99.7%) were complete in the 12-chromosome assembly, including 1,583 (95.5%) single-copy and 70 (4.2%) duplicated genes; none was fragmented and 5 (0.3%) were missing (Figure 3B). The consensus quality value was QV 70.02, corresponding to one error per 10.0 Mb, estimated from 1,851 k-mers found in the assembly but not in the reads. Per-chromosome QV ranged from 61.0 (chromosome 12) to 80.0 (chromosome 5), and chromosome 12 contributed 969 (52%) of the assembly-only k-mers. The k-mer completeness was 81.44% (523,976,574 of 643,381,822 reliable read k-mers), as expected for a primary assembly of a heterozygous genome, which omits k-mers unique to the alternative haplotype. The Merqury copy-number spectrum placed the heterozygous and homozygous peaks at 39x and 80x (Figure 3A). K-mers found once in the assembly accounted for most of both peaks, k-mers found twice formed a minor shoulder between approximately 80x and 160x, and k-mers present only in the reads formed a peak at 39x in addition to the low-multiplicity error k-mers.

**Figure 3:**
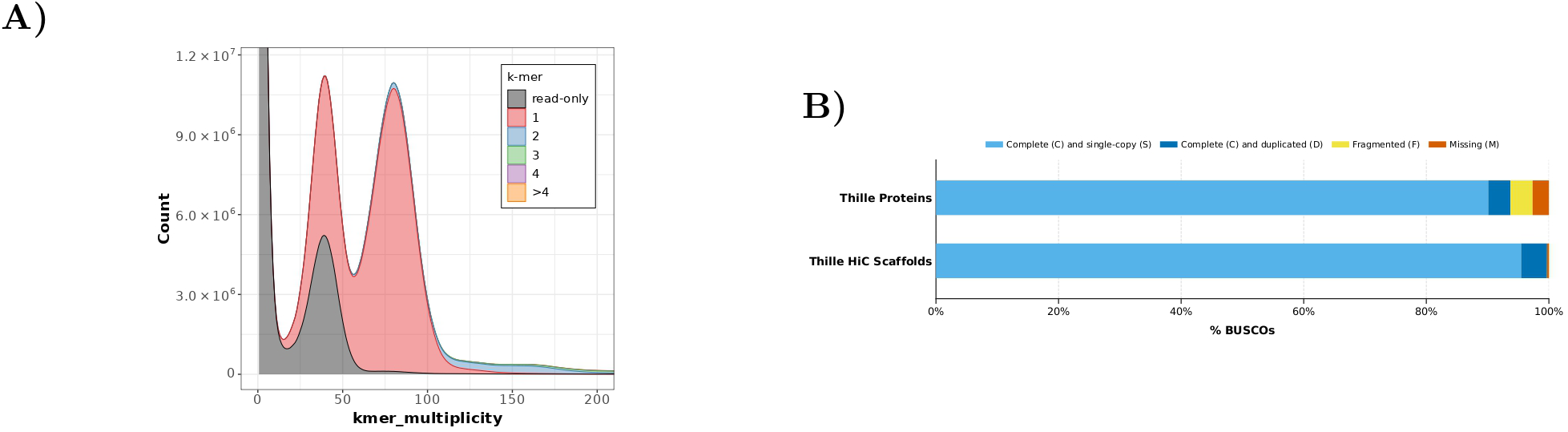
Quality and completeness of the Thille assembly and annotation. (A) Merqury k-mer copy-number spectrum (k = 21) of the 12 Thille chromosomes from Thille HiFi reads; peaks at about 40x and 80x hold k-mers from one and both haplotypes (QV 70.02, k-mer completeness 81.44%). (B) BUSCO completeness (embryophyta_odb12.2, n = 1,658) of the Tiberius proteins (C 93.7% [S 90.1%, D 3.6%], F 3.6%, M 2.7%) and of the 12 chromosomes (C 99.7% [S 95.5%, D 4.2%], F 0.0%, M 0.3%).

### Transposable Element Annotation and Masking

Repeat annotation of the 12-chromosome Thille assembly masked 566.6 Mb, or 64.0% of the 885.4 Mb assembly, of which 547.8 Mb (61.9%) were interspersed repeats and the remainder simple repeats (0.84%), low-complexity sequence (0.18%), and small RNA genes (0.99%) (Figure 2). Retroelements accounted for the largest share at 302.1 Mb (34.1% of the assembly), dominated by LTR retrotransposons at 251.9 Mb (28.5%). Gypsy elements (154.1 Mb, 17.4%) outnumbered Copia elements (78.0 Mb, 8.8%) by roughly two to one in occupied length, and together the two superfamilies made up 92% of LTR sequence. LINEs contributed 48.0 Mb (5.4%), nearly all in the L1/CIN4 (3.7%) and RTE/Bov-B (1.0%) clades, and SINEs were negligible at 0.2%. DNA transposons occupied 36.2 Mb (4.1%), led by hobo-Activator elements (2.5%) and MULE-MuDR elements (0.9%), with rolling-circle Helitrons at 0.2%. Unclassified repeats were the second largest category at 209.5 Mb (23.7% of the assembly, 38% of interspersed repeat length).

### Gene Annotation

Tiberius predicted 36,349 protein-coding genes, each with one transcript. Gene models averaged 5,706 bp (median 2,058 bp; range 201 bp to 200.4 kb) and together spanned 23.38% of the assembly, with exons covering 4.54%. Genes carried a mean of 4.3 exons (median 2), and 11,867 (32.6%) were single-exon genes. Predicted proteins averaged 368 amino acids (median 283; range 66 to 11,856). The predicted proteome recovered 1,554 (93.7%) complete embryophyta_odb12.2 BUSCO genes, including 1,494 (90.1%) single-copy and 60 (3.6%) duplicated genes, with 60 (3.6%) fragmented and 44 (2.7%) missing (Figure 3B). Gene density was highest toward chromosome ends and lowest around the centromeres (Figure 5C). The terminal 10% at each end of the chromosomes carried 62.0 genes per Mb, compared with 35.8 genes per Mb in the chromosome interiors, and ends were denser than interiors on all 12 chromosomes. Within 5 Mb of the centromeric regions, gene density fell to 17.8 genes per Mb (41.1 genome-wide), and TEs covered 86.6% of the sequence, compared with 59.4% elsewhere.

### Omni-C Contacts, Centromeres, and Telomeres

Omni-C contacts were concentrated along the diagonal within each of the 12 chromosomes in the balanced genome-wide map (Figure 4). A centromeric region was identified on every chromosome (Figure 5C). The centromeric satellite family chosen from the data had periods of 965 to 1,065 bp (seed period 1,015 bp) and totaled 29.8 Mb, of which 18.0 Mb lay in the gene-poorest 10% of windows; the next family (1,309 to 1,445 bp) contributed only 2.4 Mb to those windows. Eight chromosomes (1, 2, 3, 5, 6, 7, 8, and 12) carried 1.0 to 3.9 Mb of this family within their centromeric regions, which spanned 2.2 to 8.8 Mb. The other four (4, 9, 10, and 11) carried less than 0.1 Mb, and their centromeres were placed from Omni-C and gene density alone, in narrower regions of 0.3 to 0.8 Mb. A Omni-C centromere peak fell within 1 Mb of the called region on six chromosomes (1, 3, 6, 8, 10, and 11). Telomeric repeat arrays of 1.1 to 39.9 kb were found at 21 of the 24 chromosome ends and were absent from the right end of chromosome 3 and the left ends of chromosomes 9 and 12. Nine chromosomes were metacentric (chromosomes 1, 2, 3, 6, 7, 8, 9, 10, and 11; arm ratios 1.12 to 1.68), two were submetacentric (chromosomes 4 and 5; 2.21 and 1.95), and one was telocentric (chromosome 12; 27.61). Ranking the candidate regions by Omni-C peak height instead of satellite content selected the same region on 10 of the 12 chromosomes. The two exceptions lacked the centromeric satellite family: chromosome 4 (39.09 to 39.47 Mb instead of 44.10 to 44.38 Mb, which would make it metacentric with an arm ratio of 1.47) and chromosome 9 (25.25 Mb instead of 23.17 to 23.80 Mb, metacentric under both). After curation, chromosome 8 carries telomeric repeats at both ends, and its centromere call (30.52 to 35.51 Mb; arm ratio 1.28) moved by exactly the 25,531,594 bp added in front of the former chromosome; the centromere calls of the other 11 chromosomes are unchanged. The chromosome 12 centromere call (0 to 8.76 Mb) coincides with a segment that forms a separate block in the Omni-C map, has no telomeric repeat at its end, and has approximately 0.4 Mb of matching sequence on Hass chromosome 12; its first 4.7 Mb align, at 82 to 90% identity and in mixed orientation, to the first 1.2 Mb of Unnamed Lowland chromosome 12, and no Puebla sequence aligns to it. The segment consists of 26 contigs of 0.03 to 0.70 Mb (0 to 7.67 Mb) joined ahead of the 18.8 Mb contig that continues the chromosome. Its join to that contig is well supported, with 7.9 times as many Omni-C pairs spanning it as the median for positions inside contigs, but pairs from within the segment remain largely inside it or reach unplaced scaffolds (14 to 47% per 1 Mb bin), with 4 to 14% reaching chromosome 12 between 8 and 20 Mb.

**Figure 4:**
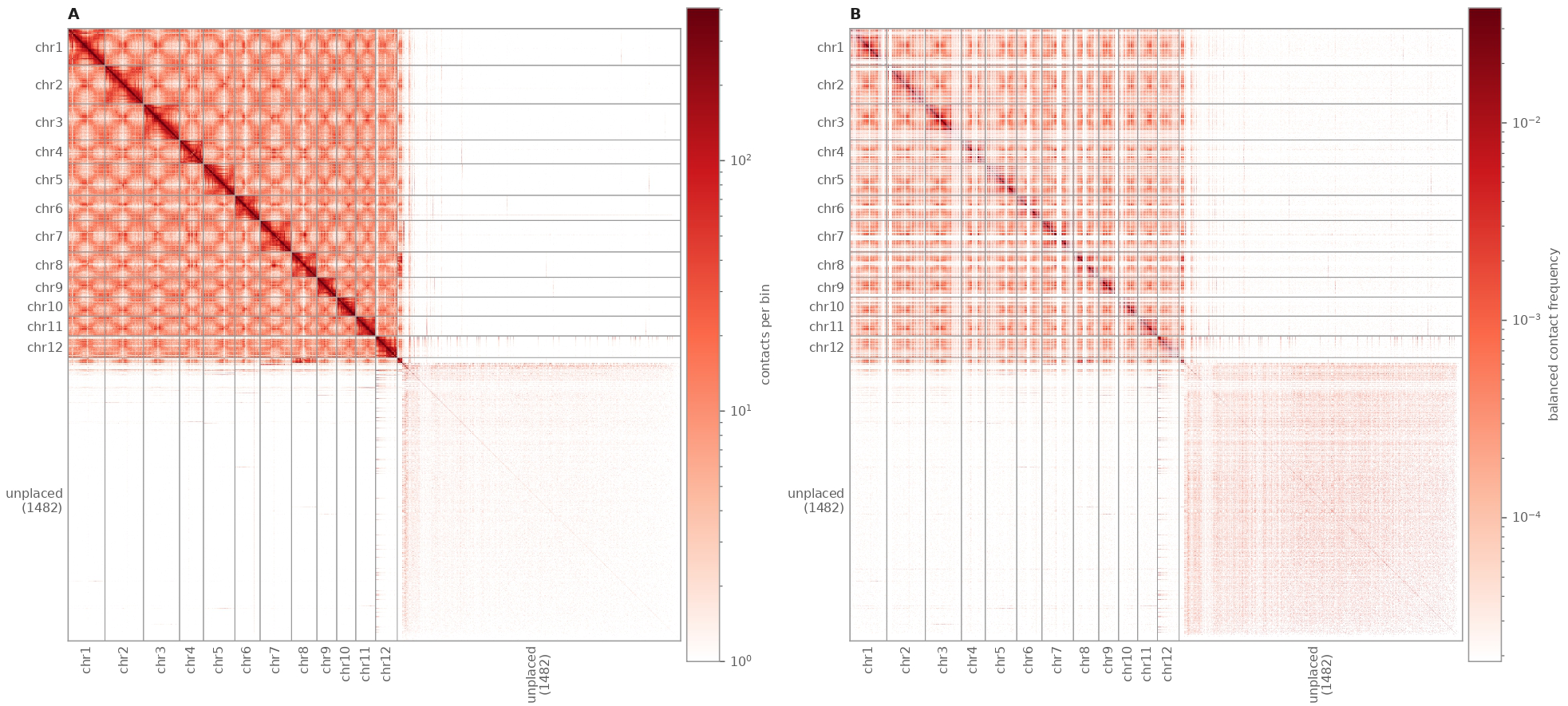
Omni-C contact map of the Thille assembly. (A-B) All 77,450,874 Omni-C read pairs (mapping quality of at least 10) in 500 kb bins, chromosomes 1 to 12 followed by the 1,482 unplaced scaffolds by size; log color scale per panel from the 1st to the 99.5th percentile of nonzero bins, white bins empty or filtered. (A) Raw contact counts. (B) ICE-balanced contact frequencies (cooler; –min-nnz 50 –mad-max 3).

### Comparative Analysis

Thille chromosome numbering was matched to Hass (UoQ) chromosome numbering for all 12 chromosomes, and the two assemblies are collinear along their full lengths (Figure 5A; Supplementary Figure S2). Six Hass chromosomes (1, 4, 5, 6, 10, and 11) are oriented on the opposite strand relative to Thille, with 99.5 to 99.9% of matched bases on each aligning to the minus strand and same-chromosome position correlations of r = 0.954 to 1.000 after reorientation (0.841 to 1.000 across all 12 chromosomes). This reverse complementation reflects the strand chosen for each assembly rather than inversions. The only internal inversion of at least 1 Mb spanned Thille chromosome 12 from 8.27 to 25.93 Mb (17.7 Mb) and Hass chromosome 12 from 0 to 18.13 Mb (14,965,427 matched bases). The first 8.3 Mb of Thille chromosome 12 had approximately 0.4 Mb of matching sequence on Hass chromosome 12, and its best hit was an unplaced Hass contig (JAQQXO010000016.1). In the ModDotPlot comparison (Supplementary Figure S15A), 2,182 of the 2,267 windows with at least 96% identity (96.3%) paired the same chromosome in both assemblies, whereas 96.2% of the windows between different chromosomes fell between 86 and 93% identity (median 89.1%). In the Thille to Gwen alignment, 849 Gwen contigs (1,022.5 Mb, 99.1% of the Gwen primary assembly) were placed on Thille chromosomes and 140 contigs (9.5 Mb) had no alignment with a mapping quality of at least 20. The Gwen contigs were collinear with the 12 Thille chromosomes. Of the 936 Gwen contigs with any alignment to Thille, 894 had at least one alignment block of 10 kb or more, and 472 of these (521.4 Mb) were oriented on the minus strand relative to Thille. The T2T assembly of Unnamed Lowland and the Puebla assembly were likewise collinear with Thille along all 12 chromosomes (Figure 5A and 5B). Projected onto the Gwen-study PCA, Puebla fell at PC1 0.129 and PC2 −0.101, about midway between the Guatemalan and Mexican accessions and among the Mexican x Guatemalan hybrids Fuerte, Zutano, Bacon, and 069-02 (Supplementary Figure S16); the two PCAs computed on the Gwen scaffolds placed Puebla in the same position. The Thille control, projected from HiFi reads, fell with Gwen, Hass, and the other Guatemalan accessions at PC1 0.077 and PC2 0.126, against 0.082 and 0.136 for the Illumina genotypes of Thille, so projected HiFi genotypes reproduce the expected placement. Each chromosome of Unnamed Lowland and of Puebla placed 96.9 to 99.9% and 98.8 to 100% of its matched bases in blocks of at least 10 kb, respectively, on the Thille chromosome of the same number, and in both assemblies the same six chromosomes as in Hass (1, 4, 5, 6, 10, and 11) were stored on the opposite strand (98.8 to 99.95% and 98.2 to 100% of matched bases on the minus strand). In the ModDotPlot comparison of Thille and Unnamed Lowland (Supplementary Figure S15B), 1,918 of the 2,003 windows with at least 96% identity (95.8%) paired the same chromosome, and 97.0% of the windows between different chromosomes fell below 93% identity (median 89.0%), the same pattern observed against Hass. Against Puebla (Supplementary Figure S15C), 2,002 of the 2,102 windows with at least 96% identity (95.2%) paired the same chromosome, and 96.5% of the windows between different chromosomes fell below 93% identity (median 89.0%).

**Figure 5:**
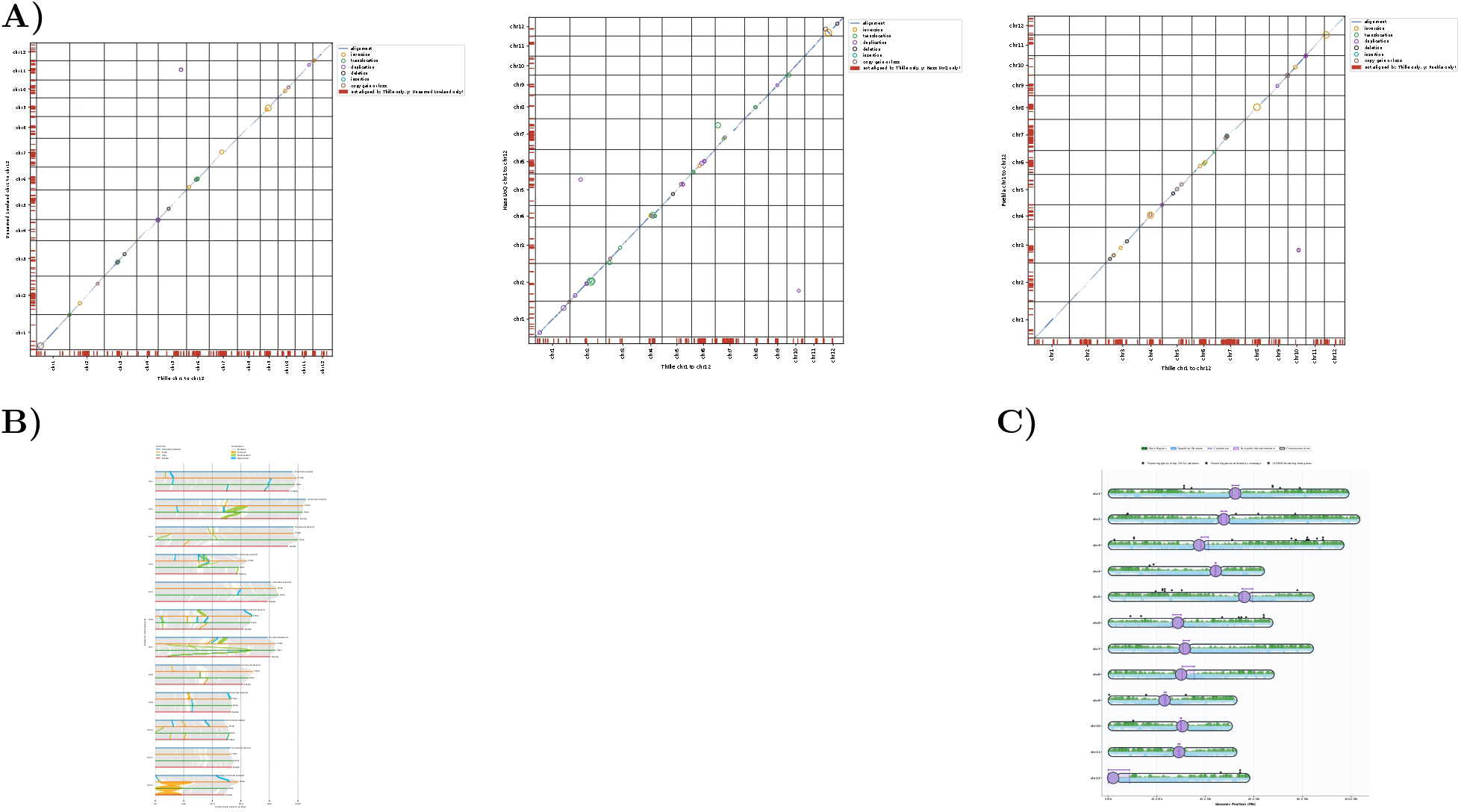
Comparison of Thille with Unnamed Lowland, Hass (UoQ), and Puebla, and the Thille karyotype. (A) SyRI dot plots of Thille against Unnamed Lowland, Hass (UoQ), and Puebla, with each query placed on the Thille strand; circles mark rearrangements of at least 100 kb (36, 43, and 39, respectively), sized by length, and red strips mark unaligned sequence of at least 100 kb. (B) Four-genome synteny and rearrangements of at least 100 kb (plotsr) for chromosomes 1 to 12 of Unnamed Lowland, Thille, Hass, and Puebla. (C) Karyotype of the 12 Thille chromosomes with gene and repeat density, the satellite-first centromere calls with their pericentromeric spans, and the candidate flowering genes; chromosomes 4 and 5 are submetacentric, 12 telocentric, and the rest metacentric.

### Large Structural Variants

SyRI placed 777.5 Mb of Thille (87.8%) in syntenic blocks with Hass, 767.8 Mb (86.7%) with Unnamed Lowland, and 758.2 Mb (85.6%) with Puebla. Rearrangements of at least 100 kb were few relative to the unaligned and highly diverged sequence (Supplementary Table S1). Against Hass, SyRI called 3 inversions, 14 translocations (2 inverted), 19 duplications (3 inverted), 2 deletions, and 5 copy losses; against Unnamed Lowland, 7 inversions, 11 translocations (2 inverted), 11 duplications (2 inverted), 2 deletions, 2 copy gains, and 3 copy losses; and against Puebla, 8 inversions, 4 translocations, 17 duplications (2 inverted), 3 deletions, 1 copy gain, and 5 copy losses. Sequence without an alignment in the other genome accounted for most of the differing sequence: 142 Thille-only (65.9 Mb) and 119 Hass-only (45.7 Mb) regions, 197 Thille-only (93.1 Mb) and 171 Unnamed Lowland-only (53.8 Mb) regions, and 235 Thille-only (100.0 Mb) and 179 Puebla-only (58.2 Mb) regions of at least 100 kb. Joining the second arm of chromosome 8 added 26 to 28 Mb of syntenic sequence on that chromosome in each comparison, and the 11 unchanged chromosomes gained a further 12 to 26 Mb, presumably because the alignments of their homologs changed once the arm was present in the reference. Thille sequence of at least 100 kb with no counterpart in Unnamed Lowland (197 unaligned regions, 2 deletions, and 3 copy losses; 95.6 Mb) was most abundant on chromosomes 7 (17.4 Mb), 5 (9.6 Mb), 12 (9.6 Mb), 6 (9.4 Mb), 4 (9.2 Mb), 2 (8.9 Mb), and 8 (8.8 Mb), and 46.3 Mb (48%) of it lay within 5 Mb of a Thille centromere call (Figure 5A). The largest such region on chromosomes 2, 5, 6, 8, and 12 (1.76 to 3.72 Mb) overlapped the centromere call, whereas the largest on chromosome 4 (2.12 Mb, 31.06 to 33.18 Mb) and on chromosome 7 (3.33 Mb, 42.13 to 45.45 Mb) lay outside it. On chromosome 7, 13 Thille-only regions totaling 14.1 Mb lie between 31 and 51 Mb, within the pericentromeric sequence. The two largest rearrangements between Thille and Hass are specific to Hass. The 22.6 Mb inversion on chromosome 12 (Thille 3.37 to 25.93 Mb) and the translocated segment on chromosome 2 (Thille 57.14 to 64.12 Mb, Hass 48.98 to 55.88 Mb) are absent from the Thille comparisons with Unnamed Lowland and Puebla, and the Hass to Puebla comparison recovers the same chromosome 12 inversion (Hass 0.07 to 17.47 Mb), so Thille, Unnamed Lowland, and Puebla share the same arrangement at both loci (Figure 5B). Because the Hass (UoQ) contigs were ordered into chromosomes with an earlier Hass assembly supported by a genetic map rather than with Hi-C data (Nath et al., 2022), these Hass-specific arrangements may reflect scaffolding errors in the Hass assembly rather than rearrangements in Hass. The largest inversion against Unnamed Lowland (1.76 Mb, Thille chromosome 9 23.01 to 24.77 Mb) overlaps the chromosome 9 centromere call, and the largest against Puebla (2.88 Mb, Thille chromosome 8 31.62 to 34.50 Mb) lies within the chromosome 8 centromere call. Puebla and Unnamed Lowland share part of their differences from Thille. Of the events of at least 50 bp called against both, 3,498 matched on Thille coordinates: 1,650 deletions, 1,677 insertions, 79 Thille-only regions (32 of them at least 100 kb), 55 duplications, 12 inversions, 11 translocations, and 14 copy gains or losses, and Hass carries the same event for 1,105 of them. The largest shared events are an inversion on chromosome 10 (Thille 20.98 to 21.29 Mb; 313 kb against Unnamed Lowland and 345 kb against Puebla), absent from Hass, and deletions of 173 kb on chromosome 3 (60.63 to 60.80 Mb) and 153 kb on chromosome 5 (31.10 to 31.25 Mb), the second also present in Hass. Several of the largest shared Thille-only regions lie in repeat-rich sequence, including the chromosome 3 satellite array (9.10 to 10.20 Mb), part of the chromosome 8 centromere region (34.81 to 36.54 Mb), and the first 3.0 Mb of chromosome 12, where the absence of an alignment may reflect repeat divergence rather than presence or absence of sequence. Of 330 Thille intervals of at least 100 kb with no Gwen contig (195 intervals, 87.3 Mb) or with two or more Gwen contigs (135 intervals, 19.0 Mb), Gwen CLR read depth placed 20 in the absent class, 116 in the half class, 168 at two copies, and 26 above two copies. The two-copy intervals without a Gwen contig mark sequence missing from the Gwen assembly rather than from Gwen, and the two-copy intervals with several contigs mark retained alternative haplotypes; the absent and half intervals are candidate deletions on both or one Gwen haplotype, and intervals above two copies are candidate duplications.

### Chromosome 3 and Chromosome 10 Loci

The Thille-only sequence on chromosome 3 (9.10 to 10.20 Mb) is a tandem satellite array rather than a deletion in the other genomes. In Thille, 995 kb of the 1.10 Mb interval consists of arrays of a satellite with a period of about 1,310 bp, and the interval is 98.9% TE and carries 22 Tiberius gene models. Between the same flanking syntenic blocks, Hass carries 0.76 Mb (CM056811.1 10.65 to 11.41 Mb) and Unnamed Lowland 0.57 Mb (CP152280.1 9.12 to 9.68 Mb) of sequence that does not align to Thille, including 592 kb and 437 kb of arrays of a related satellite with a period of about 1,390 bp; both periods fall within the 1,309 to 1,445 bp satellite family identified in the centromere analysis. HiFi read depth over the Thille array is 0.3 to 0.5 of the median. Thille reads are nearly absent from the Unnamed Lowland array and reduced over the Hass array, and Unnamed Lowland and Puebla HiFi reads are nearly absent from the Thille array (Supplementary Figure S9). The PRP8A candidate lies 0.44 Mb past the Thille array (chromosome 3, 10.63 Mb).

### Read Support for the Inversions

Raw reads support most of the inversions. Of the 52 inversions between Thille and Unnamed Lowland, the Unnamed Lowland HiFi reads aligned to Thille carried the other orientation at 35 and both orientations at 8 (43 of 52), whereas on their own assembly they followed it at 47 and showed both orientations at 5. Of the 63 inversions between Thille and Puebla, Puebla HiFi reads aligned to Thille carried the other orientation at 35 and both orientations at 12 (47 of 63), and on the Puebla assembly they followed it at 45, showed both orientations at 14, and never carried the other orientation alone (Supplementary Figure S12). Thille HiFi reads followed the Thille assembly at 96 of the 152 inversion calls and showed both orientations at 53, consistent with Thille being heterozygous at these loci. Thille reads contradicted the Thille assembly only at a 0.75 kb locus on chromosome 5 (79,039,610 to 79,040,370), called in both the Hass and the Puebla comparisons, where 28 Thille reads were split and 2 spanned the breakpoints while 128 Unnamed Lowland reads followed the Thille assembly. Across the 128 distinct inversion loci, both Unnamed Lowland and the Puebla reads aligned to Thille carried the other orientation, alone or with the Thille orientation, at 33.

### Thille-guided Scaffolds of Gwen and Hass (UoQ)

Scaffolded on Thille, the Gwen primary contigs formed 12 chromosomes of 43.9 to 115.6 Mb (864.6 Mb from 441 contigs and 429 joins), with 548 contigs (167.5 Mb) unplaced, and BUSCO found 99.3% of genes complete (91.9% single-copy and 7.4% duplicated) on these chromosomes. The Gwen alt contigs formed fragmentary chromosomes of 370.4 Mb from 3,310 contigs (BUSCO 53.4% complete), with 823 contigs (47.7 Mb) unplaced, as expected for a set that covers less than half of the genome. The broken Hass (UoQ) contigs formed 12 chromosomes of 873.3 Mb (BUSCO 99.7% complete) with two contigs (0.58 Mb, chromosome 10) unplaced. Every Hass (UoQ) contig was placed on its own chromosome, and the Thille-guided order matched the UoQ order on 10 chromosomes; on chromosome 2 the UoQ segments at 48.98 to 56.20 Mb and 56.20 to 64.85 Mb exchanged places, and on chromosome 7 the segment at 64.08 to 67.06 Mb moved, reversed, to the start of the chromosome. These two differences coincide with the chromosome 2 translocation (Hass 48.98 to 55.88 Mb) and an inverted translocation on chromosome 7 (Hass 65.99 to 67.06 Mb) called by SyRI between Thille and Hass. The chromosome 10 locus is a presence and absence difference. Thille chromosome 10 from 11.56 to 13.60 Mb (2.04 Mb) aligns continuously to Hass (CM056818.1, about 38.1 to 40.2 Mb), whereas Unnamed Lowland carries 0.66 Mb of sequence that does not align to Thille in its place (CP152287.1 37.19 to 37.84 Mb). The Thille interval is 54 to 100% TE and carries 182 short gene models (median 554 bp) and little tandem repeat (42 kb of arrays). Over its distal half (about 12.2 to 13.6 Mb), Unnamed Lowland HiFi and Puebla HiFi reads fall to 0.40 and 0.44 of the median on Thille, whereas Thille HiFi and Gwen CLR reads remain at 0.69 and 0.71, and the Hass homolog shows the same pattern (0.43 for Unnamed Lowland, 0.72 for Thille, and 0.74 for Gwen) (Supplementary Figure S9). The SDMYB candidate lies about 1.25 Mb from the locus in Thille (chromosome 10, 10.31 Mb) and about 1.3 Mb from the Unnamed Lowland interval (CP152287.1 39.11 Mb).

### Raw Read Mapping Back Alignment

Gwen CLR and Thille HiFi reads were aligned to both references (Table 3); Hass HiFi alignments were not completely available as only a fraction of what was reported in the original Hass genome paper (Nath et al., 2022) was present when downloading from NCBI. Thille HiFi reads mapped at 98.44% to Thille and 98.28% to Hass (UoQ), with a mean mapping quality of 55.00 and 53.90 and 89.09% and 87.02% of reads at a mapping quality of 60. Gwen CLR reads mapped at 84.78% to Thille and 84.82% to Hass, with a mean mapping quality of 51.42 and 51.40 and 81.29% and 81.04% of reads at a mapping quality of 60. The alignment error rate (edit distance per aligned base) was 0.0384 for Thille HiFi reads on Thille (QV 14.16) and 0.0580 on Hass (QV 12.36), and 0.1677 (QV 7.76) and 0.1731 (QV 7.62) for Gwen CLR reads on Thille and Hass, respectively. On the full assemblies, Thille HiFi reads mapped at 99.56% to all Thille scaffolds and 98.77% to all Hass (UoQ) sequences, with mean identities of 97.43% and 96.11%. Unnamed Lowland HiFi reads mapped at 99.34% to Thille and 99.33% to Hass (mean identity 92.08% and 91.85%), Puebla HiFi reads at 99.77% to Thille (93.60%), and Gwen CLR reads had mean identities of 83.98% to Thille and 83.69% to Hass (Table 3).

### Candidate flowering genes

All 52 candidate genes were placed on both references (Tables 4 to 6). Blastn located each gene in the Hass (UoQ) genome at 94.1 to 100% identity (median 99.2%), with 49 of the 52 genes covered over at least 90% of their length, and the tblastn protein search recovered the same locus for 47 genes. On transfer to Thille, the 1 kb flanks tiled contiguously with the gene for 45 genes and with gaps of 1 to 117 bp for four more (ALP1, CCR1, FPP7, and PRP8A); the remaining three (EMF1, PAR1, and KCS11) were placed from the spliced alignments and curated manually. The Thille placements had a mapping quality of 60 for 50 of the 52 genes (PAR1, 33; KCS11, 11), and each gene was placed on the same chromosome number in both assemblies except PAR1, whose placement was corrected manually. Allelic distribution graphs and read coverage in 100 kb windows on Thille are shown for the six genes with an allele carried by all three A-type cultivars and by neither B-type cultivar, SDMYB, AGAMOUS-like, CSU2, NPR5-BOP2, SGRL, and LCYB (Figures 6 and 7); allelic distribution graphs of SPA1, VOZ1 (ChrU.ver1.g387390; Thille g22280), NRX1, and the two CCR1 genes (Thille g512 and g514) are in Supplementary Figure S11, and their read coverage in Supplementary Figure S13. The remaining candidate genes are shown in Supplementary Figures S3 and S4. SDMYB, SPA1, SPA1-like, and the two loci listed as EOBI in the Gwen study were each located on both references as single full-length hits at 98.9 to 100% identity, and in each case the 1 kb flanks tiled contiguously with the gene on Thille (Tables 4 and 6); the coordinates below are those of the selected Hass (UoQ) and Thille (Tiberius) gene models. SDMYB maps to Hass chromosome 10 (CM056818.1, 41,397,672 to 41,405,576) and Thille chromosome 10 (10,311,765 to 10,319,665; Tiberius g34985), on opposite strands, at 7,905 and 7,901 bp. SPA1, the gene under the A/B Fst peak, maps to Hass chromosome 1 (CM056809.1, 67,285,700 to 67,310,269; 24,570 bp; MRB53_002480) and Thille chromosome 1 (31,235,863 to 31,267,407; 31,545 bp; g6249). The SPA1-like paralog reported as SPA1 in Table 2 of Solares et al. (2023) maps to Hass chromosome 10 (3,265,869 to 3,315,640; 49,772 bp; MRB53_030155) and Thille chromosome 10 (47,790,066 to 47,818,305; 28,240 bp; g36229); the Hass locus carries two insertions absent from Thille, of approximately 4.4 kb and 770 bp, and the Thille locus contains a repeated element of approximately 900 bp with more than 40 copies on chromosome 10. The chromosome 7 EOBI locus maps to Hass CM056815.1 (61,934,518 to 61,936,584; 2,067 bp; MRB53_024002) and Thille chromosome 7 (63,591,299 to 63,593,468; 2,170 bp; g18626), and the chromosome 5 EOBI locus maps to Hass CM056813.1 (74,932,829 to 74,935,306) and Thille chromosome 5 (11,704,616 to 11,707,093; g13504), 2,478 bp in both. Searching the Petunia EOBI protein against the Thille assembly recovered the R2R3 domain of approximately 200 MYB family members across all 12 chromosomes. Ranked by summed bitscore, the six closest homologs were on chromosomes 10 (2.70 Mb), 5 (11.70 Mb), 9 (2.27 Mb), 10 (11.66 Mb), 9 (35.31 Mb), and 9 (2.29 Mb), and the chromosome 7 locus ranked seventh at 52.8% identity across the domain. The Arabidopsis SPA1 protein recovered four SPA-family loci on Thille chromosomes 1, 10, 6, and 8, of which the chromosome 1 and chromosome 10 loci were near-tied on summed bitscore (759 and 768). Of the 110 Thille gene models searched against Swiss-Prot, 85 had an accepted hit and 11 were flagged with a predicted flowering function (Tables 4 to 7). The UniProt keyword Flowering (KW-0287) was carried by the hits of four candidates (AP2, SDMYB, the AGAMOUS-like gene, and ELF3), and flowering GO terms by seven others (HOS1, NPR5, NAC25, AGD5, VOZ1, and two annex genes that were not retained, ATX2 and UBP18).

**Table 2:**
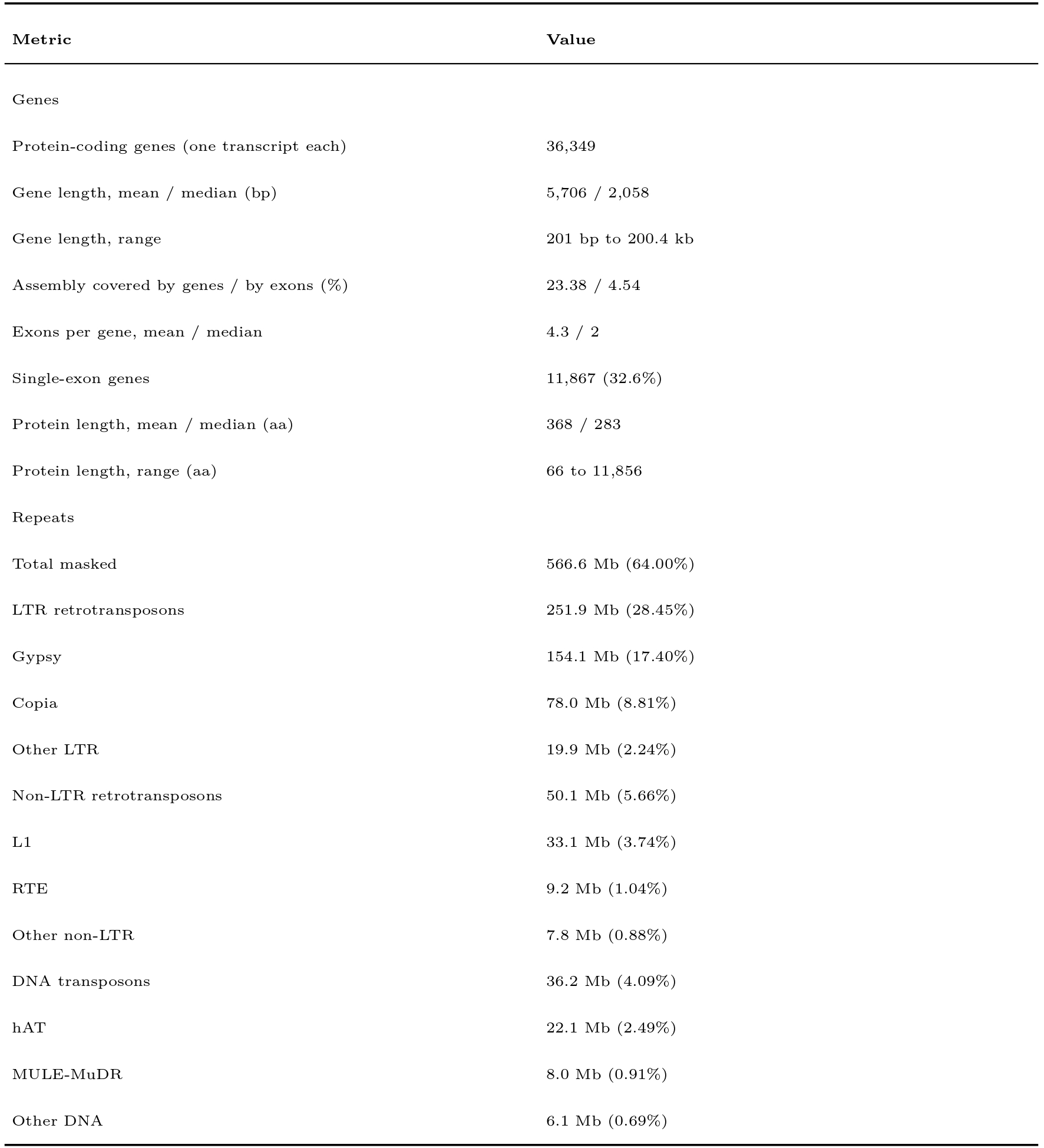
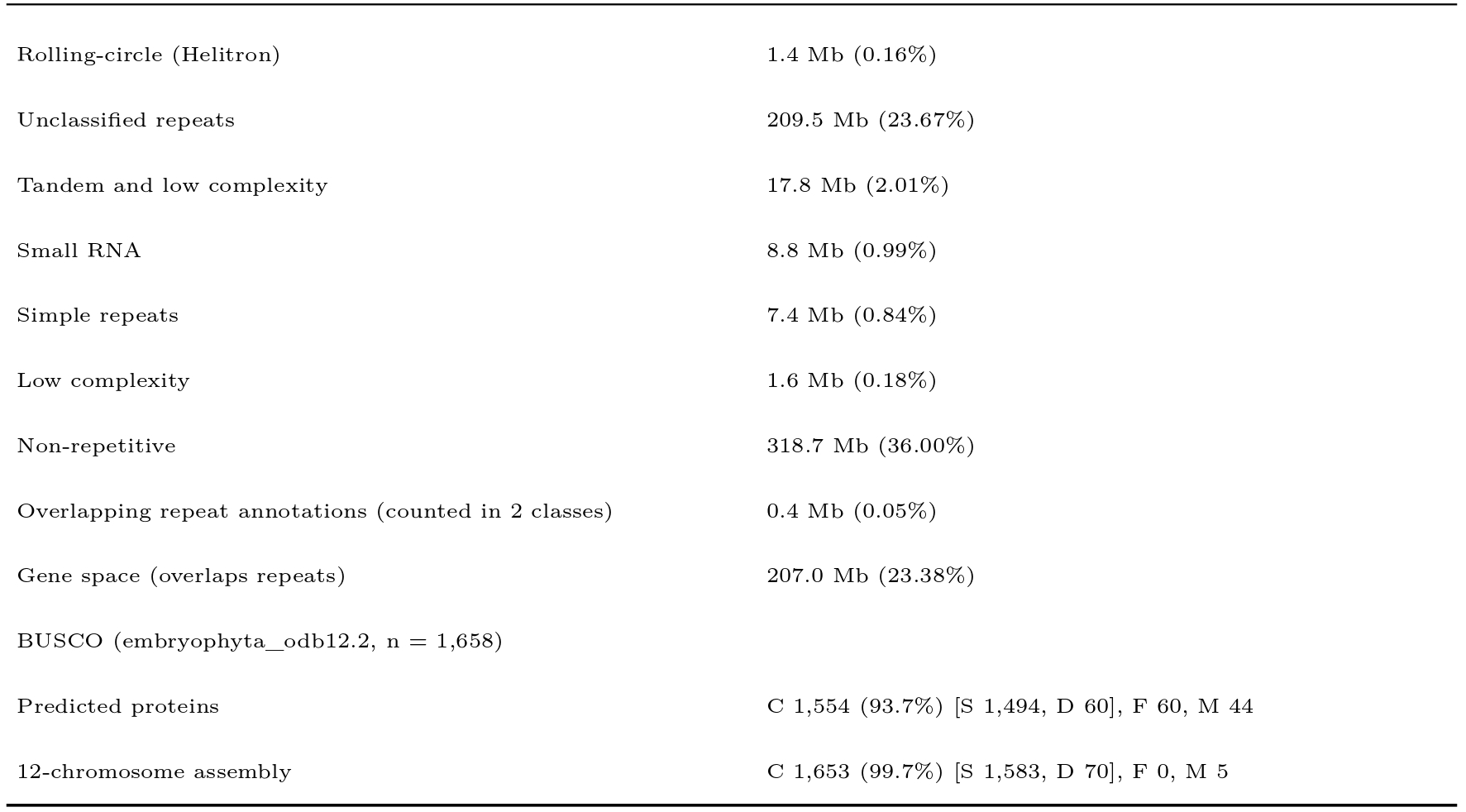
Annotation. Gene models: Tiberius; repeats: RepeatMasker v4.2.4 on the 12 chromosomes (885.4 Mb), classes as in Figure 2; BUSCO: embryophyta_odb12.2, genome mode (miniprot) and protein mode.

**Table 3:**
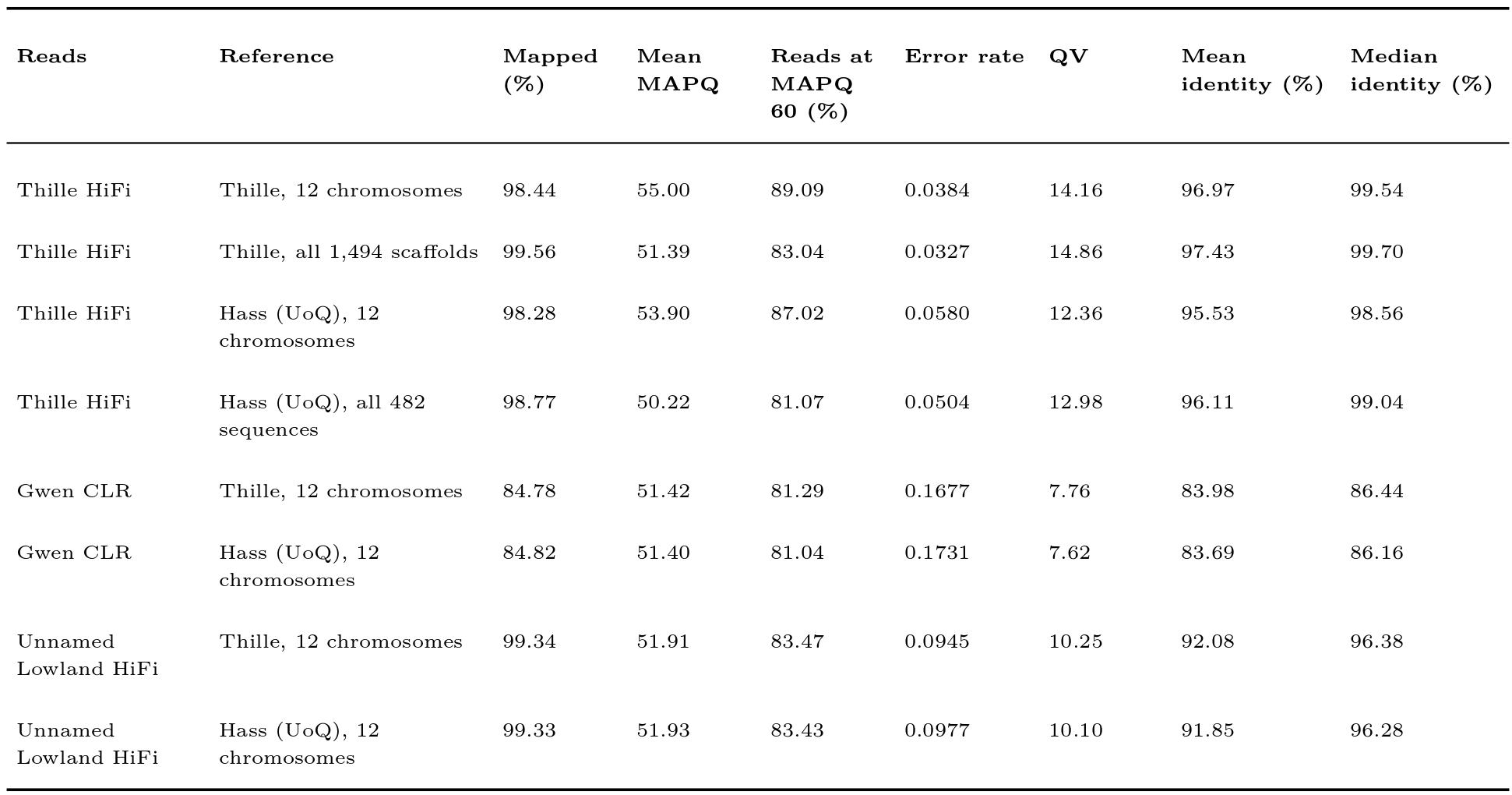
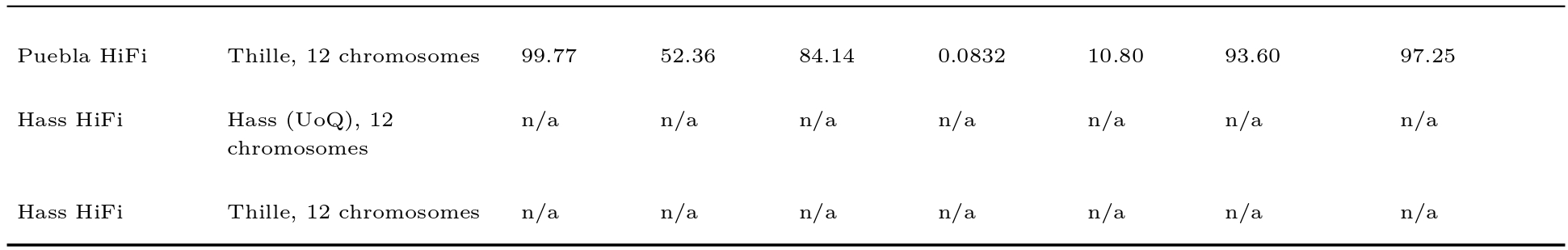
Raw read mapping to the Thille (v4) and Hass (UoQ) assemblies, on the 12 chromosomes and, for Thille HiFi reads, on the full assemblies (all Thille Omni-C scaffolds and all Hass (UoQ) sequences). Error rate is the edit distance (NM) per aligned base over primary and supplementary alignments, and QV = -10 log10(error rate); identity is the BLAST identity (matching bases over alignment columns) of the primary alignment of each read, given as the mean and median over reads. Hass HiFi rows are n/a because the Hass HiFi reads available from NCBI were a fraction of those reported (Nath et al., 2022).

### Flowering Gene Screen of the Combined Top 1% Genes

The flowering screen of the 944 genes in the union of the contig and scaffold top 1% sets yielded 28 candidate gene models representing 24 genes after manual checking (Table 4). Seventeen of the 24 genes were not among the nine genes reported in Table 2 of the Gwen study (Solares et al., 2023): EMF1, HY5, TIC, AGAMOUS-like, ALP1, ARF2A-like, ICR1, MIK2 (chromosome 12), NAC25, NRX1, PAR1, PLA2/LHD2, ZFP2, FBD/RNI-like, RER3, RPT3, and PBL34. The other seven were recovered by the screen: VOZ1, CCR1, SPA1, AP2, NPR5, CSU2, and SDMYB, which Table 2 of the Gwen study lists as EOBI. Nine of the 17 added genes lie in top 1% windows of the contig analysis only (TIC, AGAMOUS-like, ARF2A-like, ICR1, PLA2/LHD2, ZFP2, FBD/RNI-like, RER3, and PBL34).

**Table 4:**
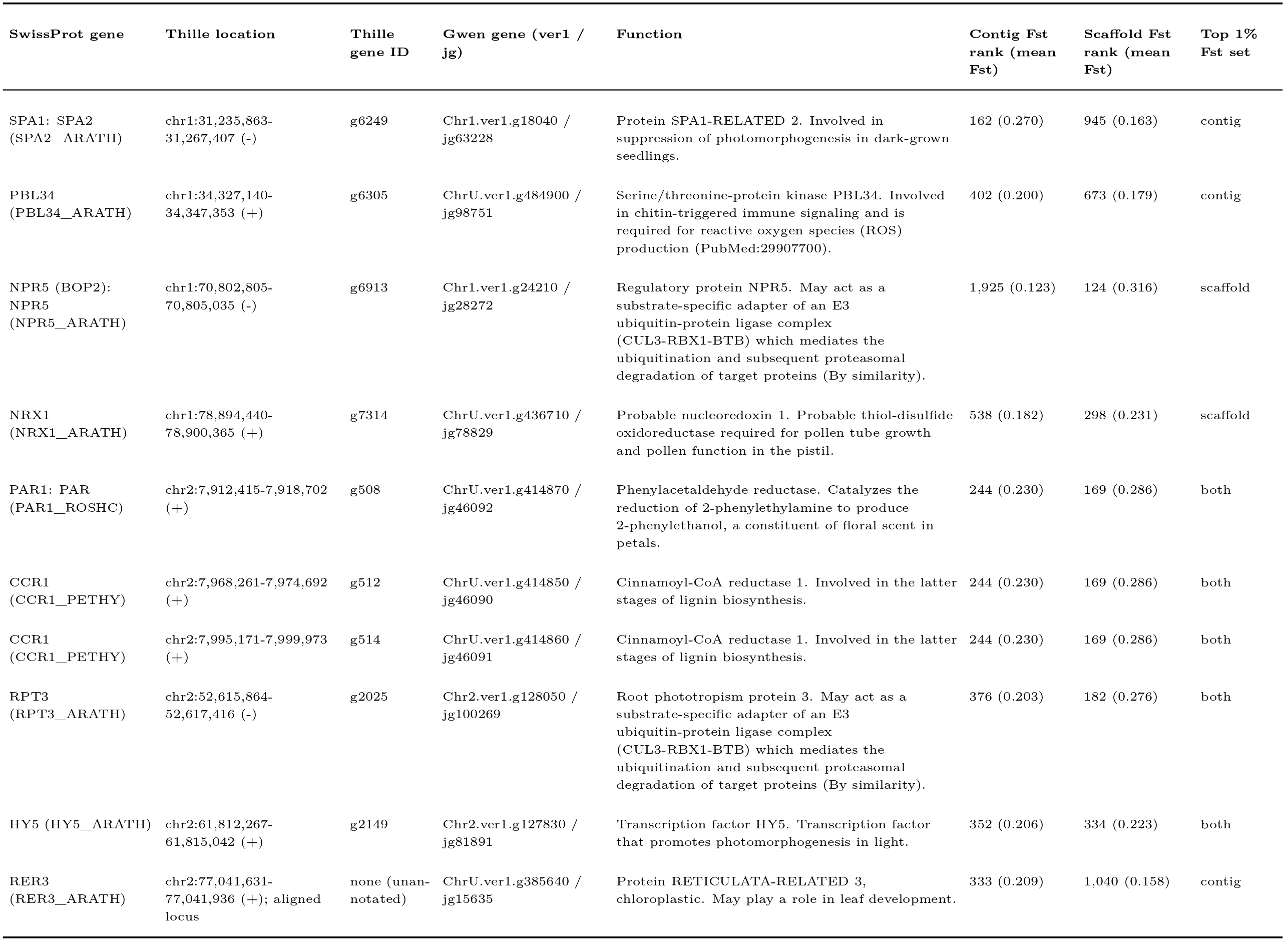
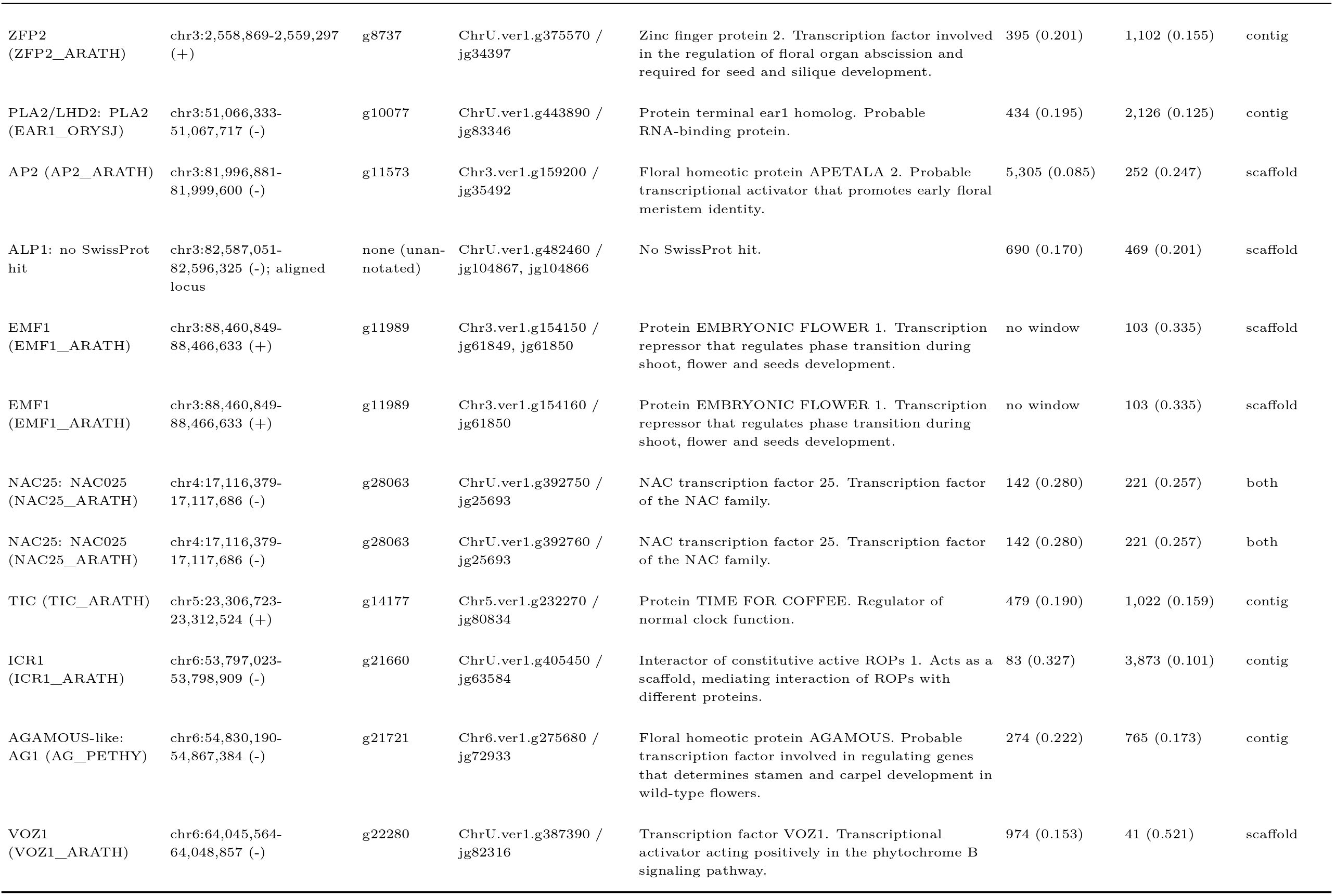
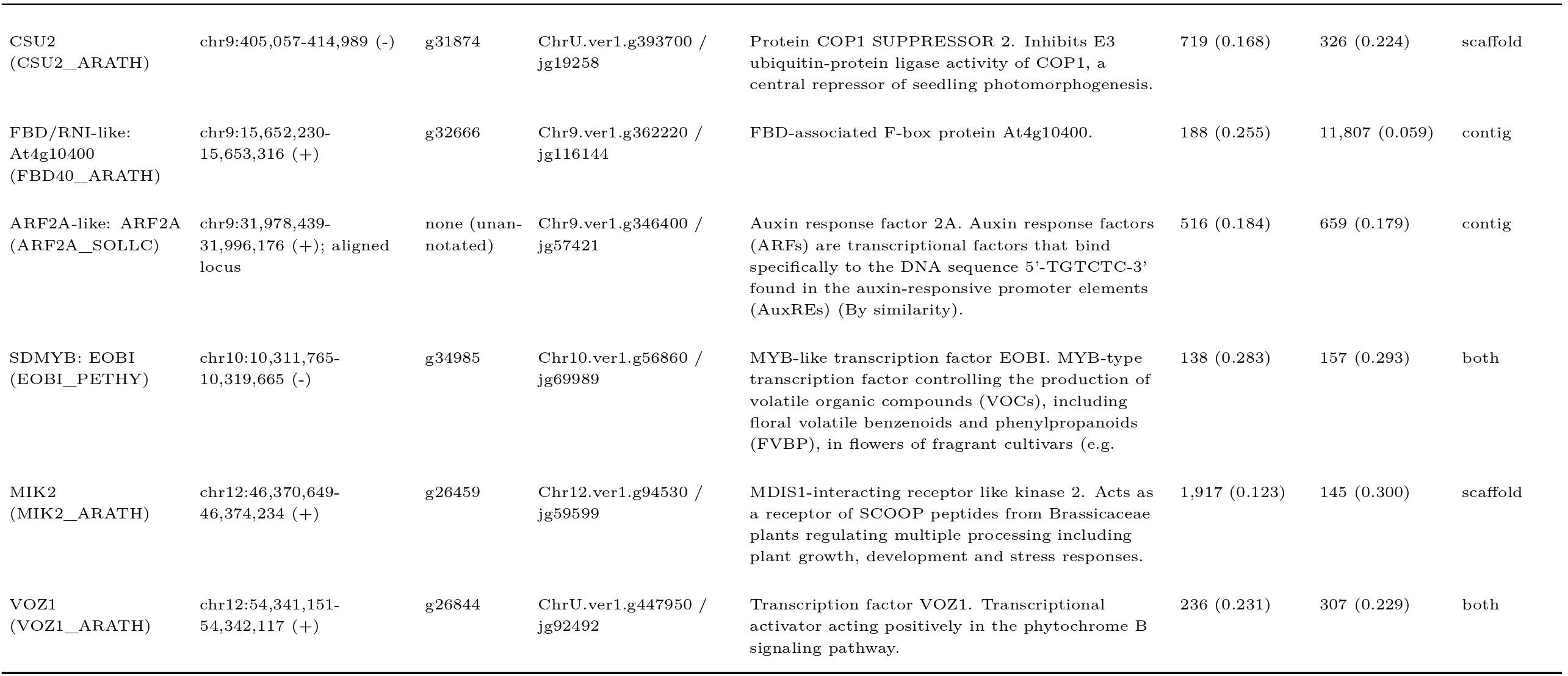
Candidate flowering genes in top 1% Fst windows (n = 28)

### Contig and Scaffold Fst of Candidate Genes

The 26 loci of the Fst candidate table (Table 4) fall in top 1% windows of both analyses for 8 loci (CCR1 x2, HY5, NAC25, PAR1, RPT3, SDMYB, and VOZ1 on chromosome 12), of the contig analysis only for 10 (AGAMOUS-like, ARF2A-like, FBD/RNI-like, ICR1, PBL34, PLA2/LHD2, RER3, SPA1, TIC, and ZFP2), and of the scaffold analysis only for 8 (ALP1, AP2, CSU2, EMF1, MIK2, NPR5, NRX1, and VOZ1 on chromosome 6). EMF1 has no contig rank, since its Gwen contig carries no window in the contig analysis. Nine of the 14 genes with coverage differences in the Gwen study also lie in top 1% windows (Table 6): APT3, FPP7, MTERF2, OTU7, RS16, SUD1, UPL2, and Y5713 in the scaffold analysis (ranks 195 to 441), and KCS11 in both (contig rank 154, scaffold rank 280). The two loci listed as EOBI under their published ver1 IDs (Chr5.ver1.g224860 and ChrU.ver1.g377230), which encode other MYB proteins, and SPA1-like rank at 10,870 or lower in both analyses. The only gene among the 944 top 1% genes with an EOBI hit is SDMYB (best hit EOBI_PETHY; contig rank 138, scaffold rank 157; Table 1). None of the 11 coverage-selected flowering loci reaches the top 1% of either analysis; the closest are AGD5 (scaffold rank 640) and FTIP7 (contig rank 1,368) (Table 5). The FLOR-ID screen of the 944 top 1% genes matched seven Gwen gene models, which correspond to five genes already among the candidates (SPA1, EMF1, AP2, TIC, and VOZ1), and identified no additional flowering-time gene. Twelve further top 1% genes carried indirect flowering annotation, and all 12 were placed on both references, with the tblastn search recovering the same Hass locus and the 1 kb flanks tiling contiguously on Thille in every case. Four were added to the candidate table (Table 4): FBD/RNI-like (Chr9.ver1.g362220; contig rank 188), annotated by Blast2GO with negative regulation of long-day photoperiodism, although its best SwissProt hit (FBD40_ARATH) carries no flowering annotation; RER3 (ChrU.ver1.g385640; contig rank 333), with the GO term photoperiodism; RPT3 (Chr2.ver1.g128050; contig rank 376, scaffold rank 182), with the GO term blue light signaling pathway; and PBL34 (ChrU.ver1.g484900; contig rank 402), with GO terms for the maintenance of shoot and root meristem identity. The remaining eight Gwen gene models, representing six genes (Ole e 10-like, LIP1, ZEP, CIGR2, a CCCH zinc finger, and CTR1), are listed in Supplementary Table S2. Figure 5C marks the 35 Thille loci of top 1% Fst genes, the 11 loci of coverage-selected flowering genes, and the 9 loci of FLOR-ID genes. The Fst loci lie on nine of the 12 chromosomes, with none on chromosomes 7, 8, and 11, and 11 of the 35 lie on chromosome 3, where six (RS16, Y5713, SUD1, OTU7, FPP7, and AP2) fall within 115 kb (81.88 to 82.00 Mb) and appear as a single cluster at the scale of the figure. Two intervals carry most of this clustering (Figure 5C). On chromosome 3, the 18.3 Mb of the long arm from 73.3 to 91.6 Mb holds 13 of the 55 loci in the gene tables, which form 8 of the 47 regions that remain after loci within 200 kb of one another are merged, whereas the interval holds 2.9% of the 36,349 Tiberius genes, or 1.4 regions expected from gene density. The interval contains three genes with a flowering function: AP2, which promotes floral meristem identity (Jofuku et al. 1994; scaffold rank 252); EMF1, a repressor of the transition to flowering (Moon et al. 2003; scaffold rank 103); and FTIP7, an FT-interacting protein (UniProt Consortium, 2025) selected by coverage. It also contains the candidate ALP1 (scaffold rank 469), the 115 kb group formed by AP2 and five genes from the Gwen study (RS16, Y5713, SUD1, OTU7, and FPP7), UPL2, GAUT13, and two genes with indirect flowering annotation (CIGR2 and LIP1). On chromosome 5, the 10.8 Mb from 19.6 to 30.3 Mb holds 5 loci in 5 regions (MTERF2, ELF3, TIC, SEC24A, and PDIL2-1), against 1.7% of Tiberius genes, or 0.8 regions expected. These include the two circadian clock genes of the candidate set, ELF3 (selected by coverage) and TIC (contig rank 479), which are both FLOR-ID genes and lie 0.95 Mb apart. Both intervals are depleted of TEs relative to the genome: TEs cover 42.6% of the chromosome 3 interval and 45.1% of the chromosome 5 interval, against 63.0% genome-wide (percentiles 0 and 13 among windows of the same length), and their 100 kb, 1 Mb, and 5 Mb flanks are similarly depleted (35.2 to 46.9%). SV density in the same regions ranged from 150 to 170 SyRI events of at least 50 bp per Mb, above that of most windows of the same length (percentiles 75 to 92). Both intervals are conserved in Hass (UoQ). Hass chromosome 3 carries UPL2, the AP2 group, FTIP7, and EMF1 in the same order from 80.4 to 91.6 Mb, and Hass chromosome 5 carries the five chromosome 5 loci in reverse order, consistent with its opposite orientation relative to Thille, from 55.8 to 67.0 Mb, with ELF3 and TIC again 0.95 Mb apart.

**Table 5:**
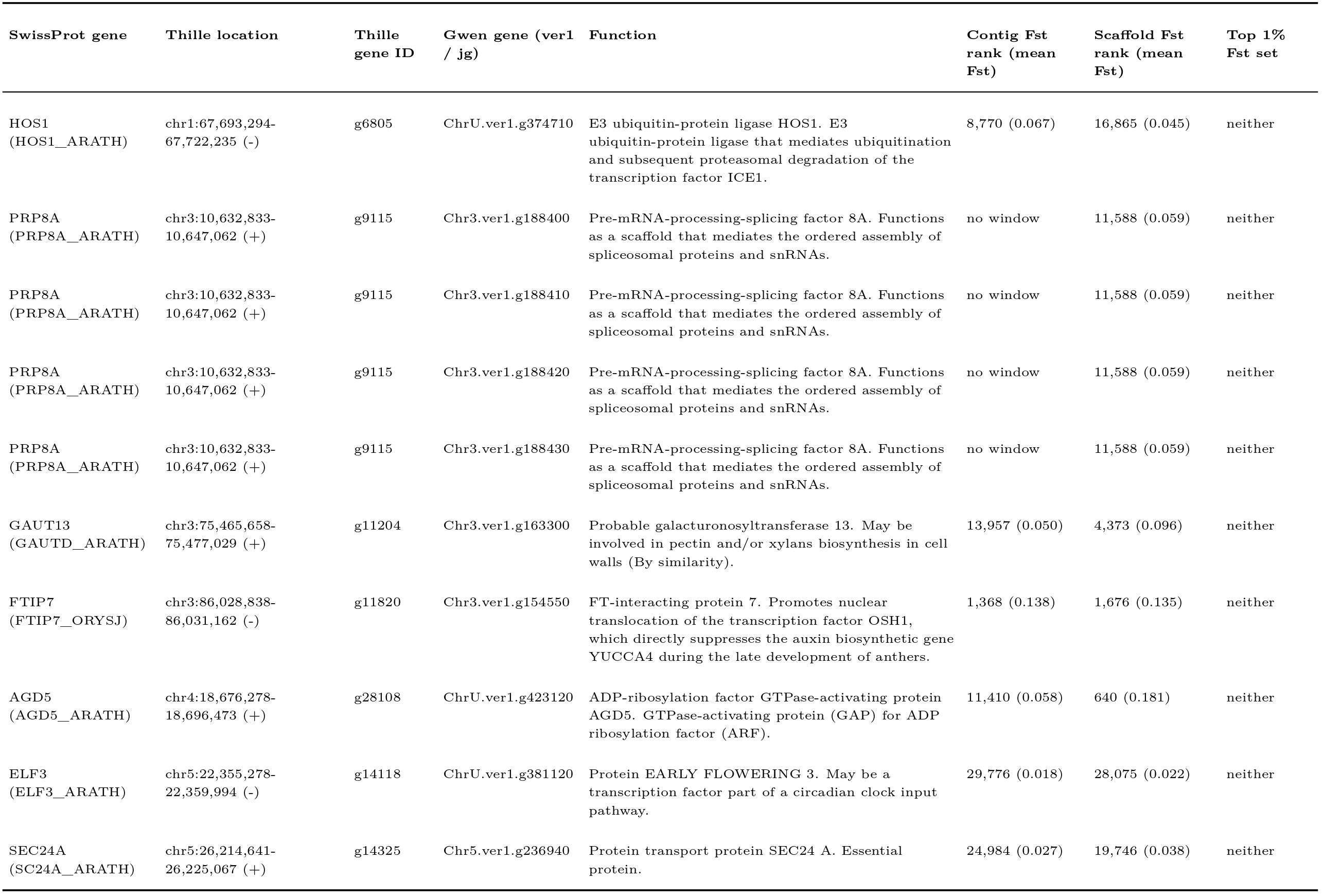
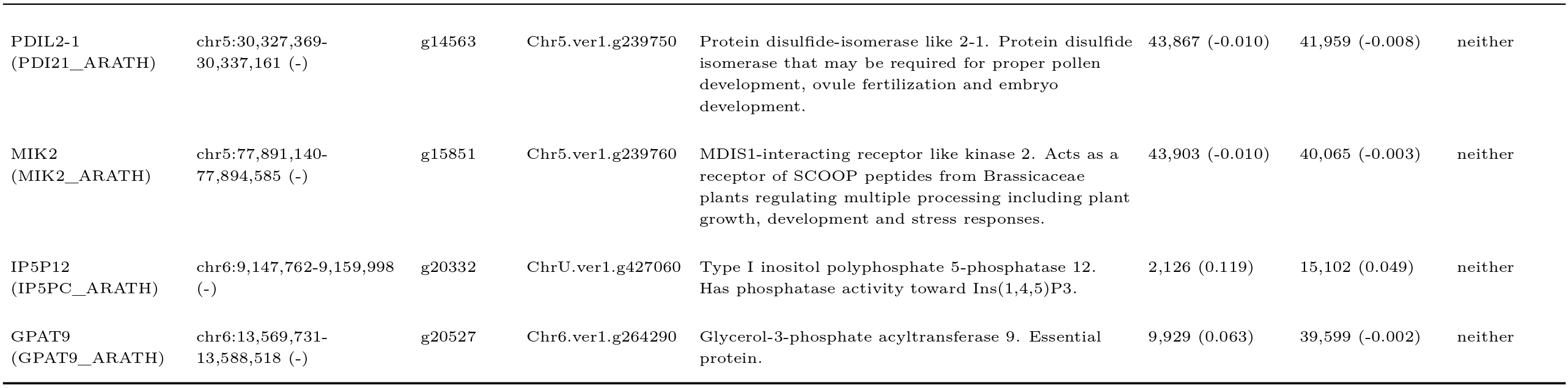
Flowering genes without Fst peaks (selected by coverage) (n = 14)

**Table 6:**
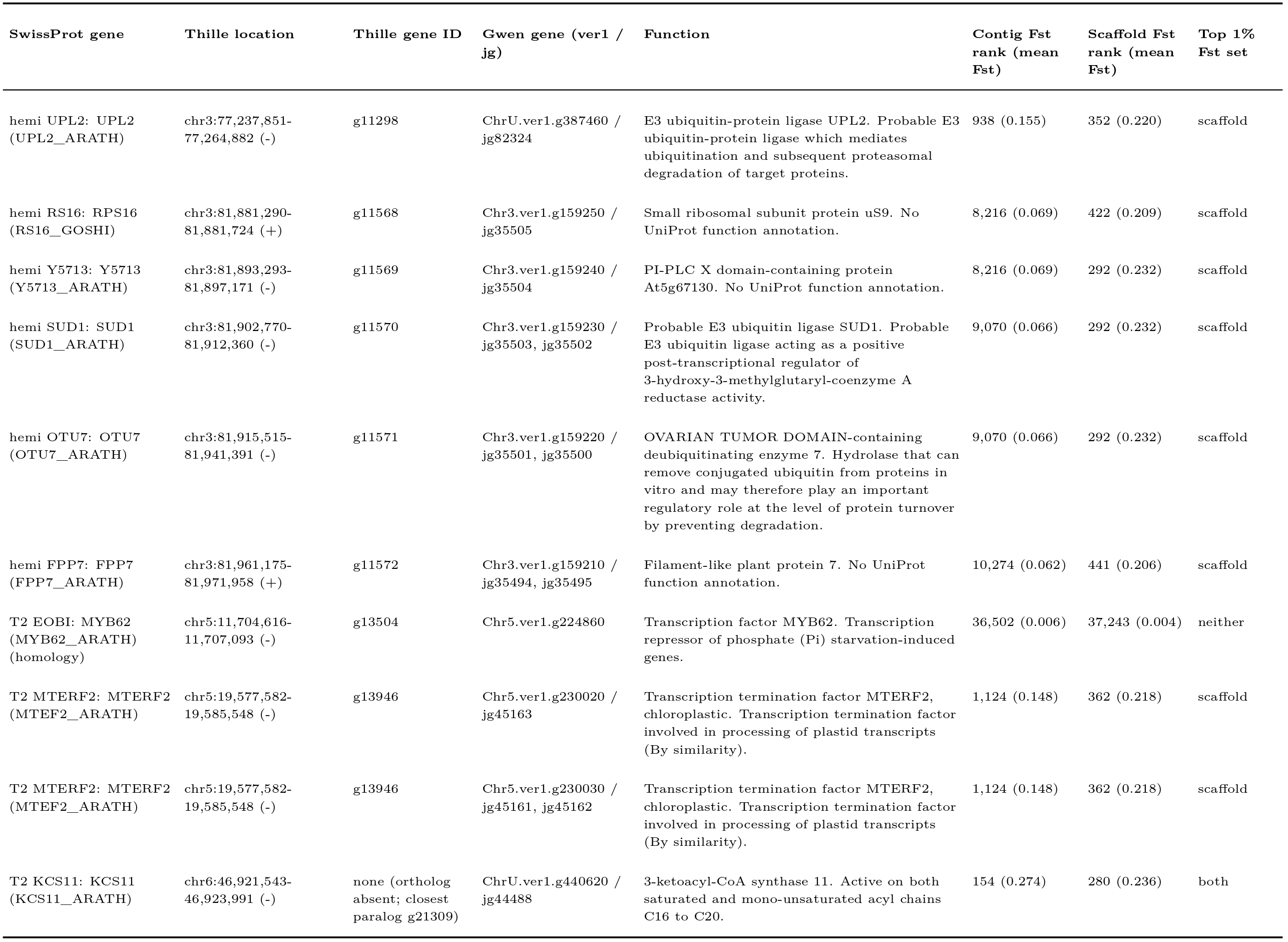
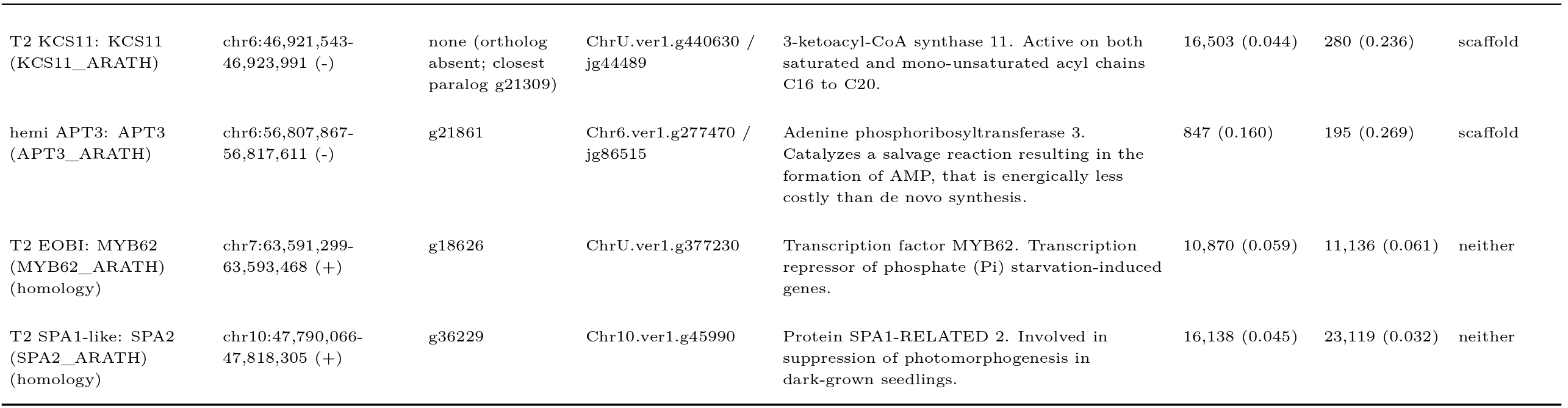
Genes with coverage differences from the Gwen study (n = 14)

### Gene Models of RER3 and the Ole e 10-like Gene

RER3 and the Ole e 10-like gene lack Tiberius models in Thille, although the corresponding sequence is present. The 101-residue Gwen RER3 protein aligns over its full length at 99 to 100% identity to Thille, Unnamed Lowland, Hass (UoQ), both Hass (MH2022) haplotypes, and both Puebla haplotypes, but it corresponds to residues 213 to 308 of the 337-residue Arabidopsis RER3, and the UoQ model at this locus spans the same 306 bp, so the existing models represent a fragment of the gene. The 124-residue Gwen Ole e 10-like protein aligns as a single 121-residue match at 99% identity in Hass (UoQ) and in one Hass (MH2022) haplotype, but as two overlapping matches in different reading frames (94 and 64% identity) in Thille, Unnamed Lowland, the other Hass haplotype, and both Puebla haplotypes, and base-level alignment confirmed a frameshift in these copies: against the 426 bp open reading frame of the intact UoQ copy, they carry a 13 bp insertion near position 123 (split into 8, 3, and 2 bp in Unnamed Lowland, which also carries a 2 bp insertion near position 369), so that a stop codon falls at codon 56, whereas the UoQ copy and the alternate Hass (MH2022) pseudohaplotype encode an intact 141-residue protein. Neither the Thille nor the Unnamed Lowland annotation contains a gene at this locus.

### SDMYB Alleles in A-Type and B-Type Cultivars

Phasing of the reads aligned to Thille resolved two SDMYB haplotypes in each read set (Figure 6B). Both Thille haplotypes and both Unnamed Lowland haplotypes carried a CDS identical to the Thille reference over all 573 bp (SDMYB^a^, the recessive allele). Gwen carried SDMYB^a^ on one haplotype (91 reads; AF 70%) and SDMYB^b^ on the other (37 reads; AF 30%). SDMYB^b^ differed from SDMYB^a^ at 17 coding substitutions, 8 of them amino-acid changing (N7S, D65G, S139G, D144N, T158I, T163M, A167T, and S170N), and carried deletions of 3 bp and 1 bp in the final exon that shift the reading frame from codon 183; translated to its own stop codon in the assembled sequence, SDMYB^b^ encodes a 190-residue protein ending in EDLWFHCSALTE, compared with EDLWSSHLFSAD in SDMYB^a^. The SDMYB^b^ CDS was identical in Gwen haplotype 2, Puebla haplotype 2, and the alternate Hass (MH2022) pseudohaplotype, whereas Puebla haplotype 1 and the principal Hass pseudohaplotype carried SDMYB^a^. The single-sequence Hass (UoQ) assembly matched the principal pseudohaplotype at SDMYB, and over all 29 gene windows it matched one pseudohaplotype variant for variant (12 the principal, 16 the alternate, and 1 both), so it represents one Hass haplotype per locus. An alignment-free count of 21-mers spanning ten of the coding substitutions in the raw reads agreed: Unnamed Lowland HiFi reads contained 701 SDMYB^a^ and no SDMYB^b^ 21-mers, Thille HiFi reads 974 and 1, Gwen CLR reads 149 and 78, and Puebla HiFi reads 238 and 130. At SDMYB, Unnamed Lowland therefore carries two copies of the recessive allele, as does the B-type Thille, and each A-type cultivar carries one copy of the same dominant allele.

**Figure 6:**
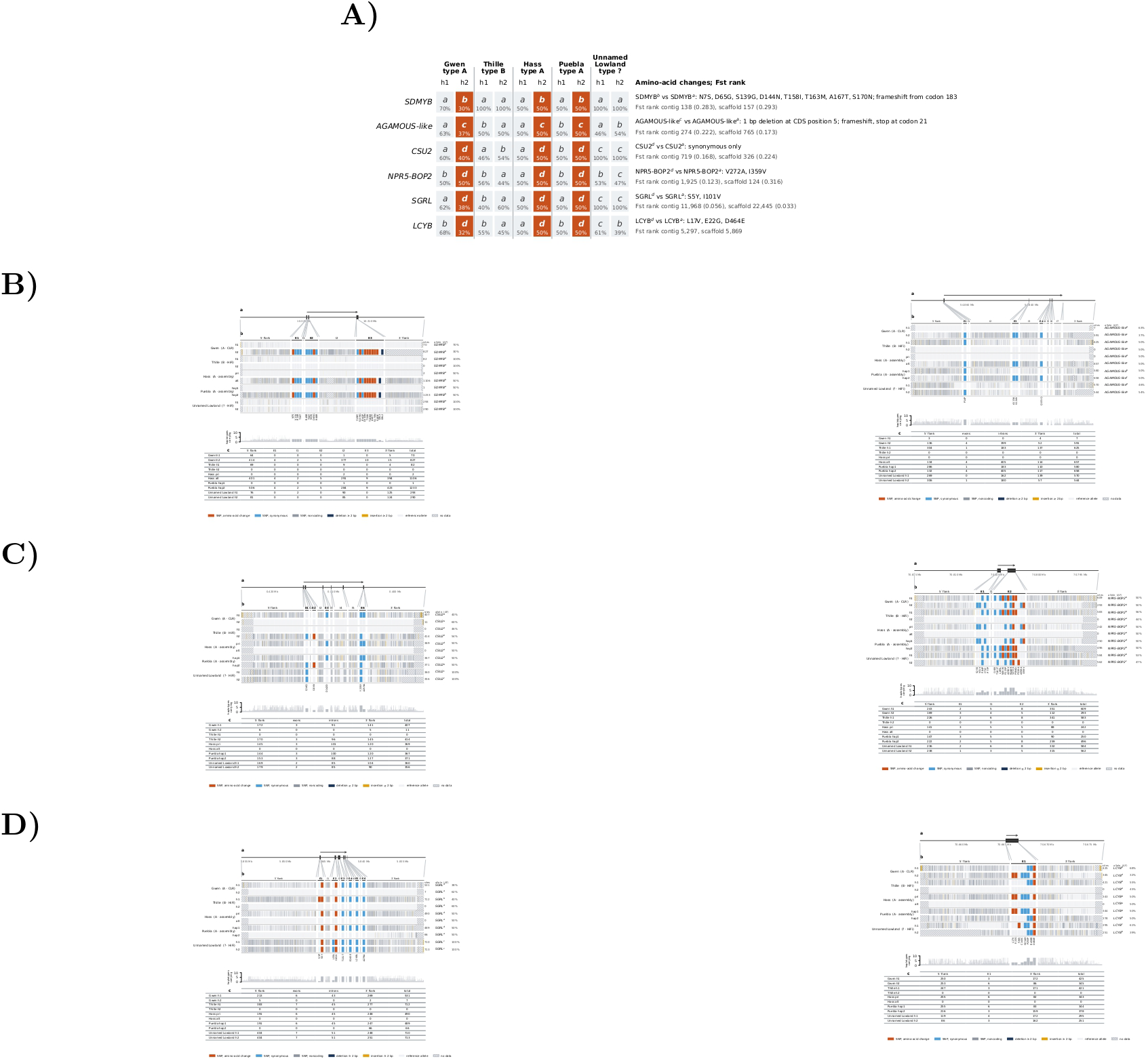
Alleles of the genes that separate the B-type and A-type genomes. (A) CDS allele and allele frequency (AF) of each haplotype at the six genes where Gwen, Hass, and Puebla each carry an allele absent from Thille and Unnamed Lowland (filled cells). Letters are CDS alleles, with a the Thille reference CDS, and numbers are AF. Amino-acid changes of the A-shared allele are given as reference residue, position, and A-shared residue (N7S: Asn7 to Ser), and Fst ranks are type A versus type B ranks of the Gwen 20 kb windows, with the mean Fst in parentheses (top 1% = ranks 1 to 517 of the contig analysis and 1 to 508 of the scaffold analysis). Haplotypes are not phased between genes, so the A-shared allele is drawn on the second haplotype of each A-type cultivar. Unnamed Lowland carries none of the six CDS alleles, although at the protein level it encodes the SGRL^d^ protein on both haplotypes. (B-D) Allele matrices on Thille: (a) the Thille locus, gene drawn 5*^′^* to 3*^′^*; (b) variant sites (columns) carried by each haplotype (rows), with the allele letter and allele frequency (AF) of each haplotype; (c) carried sites per region and haplotype. Allele a is the Thille reference CDS. A haplotype carries a variant when at least 50% of its reads carry it, or otherwise its most frequent allele; for unphased read sets, the most frequent allele is drawn on h1 and the second (at least 25% of reads) on h2, and for assemblies the variant is present in the aligned sequence. Variants are relative to the Thille assembly. Haplotypes share an allele letter only when their CDS are identical, andmarks a CDS not fully covered by the reads of that haplotype. AF is the share of the cultivar’s reads carrying that exon allele, pooled over CDS sites where its two haplotypes differ and counting the two most frequent alleles per site; 100% means both haplotypes share the exon allele, and assembled haplotypes are 50% or 100%. (B) SDMYB (g34985, chromosome 10) and AGAMOUS-like (g21721, chromosome 6). (C) CSU2 (g31874, chromosome 9) and NPR5-BOP2 (g6913, chromosome 1). (D) SGRL (g30157, chromosome 11) and LCYB (g6907, chromosome 1).

### Flowering-Type Allele Classes

SDMYB, the positive control, had 474 variant sites that followed the flowering-type pattern among the 1,220 sites with data in all ten haplotypes, including 17 coding sites carrying the eight amino-acid changes of SDMYB^b^, and its coding and protein alleles both matched the model. Among the other genes, AGAMOUS-like had the most pattern sites (309), followed by CTR1 (234), KCS11 (153), SGRL (131), NPR5-BOP2 (78), SPA1 (77), CIGR2 (71), CSU2 (69), LCYB (54), and RPT3 (34), and 17 genes had none. Sites at which any haplotype lacked data were not scored; at PAR1, for example, 1,989 of 2,553 variant sites were not scored, and no site could be scored at ZFP2 or FBD/RNI-like, which could not be placed on Puebla haplotype 2. At the level of whole alleles, AGAMOUS-like was the only gene besides SDMYB to match the model at the protein level: the four B haplotypes encode the Thille reference protein, and each A individual carries one copy of the same variant protein (AGAMOUS-like^c^), in which a 1 bp deletion at CDS position 5 shifts the reading frame to a stop codon at codon 21; its CDS alleles do not match the model, because the two Thille haplotypes differ at synonymous positions. Ten genes could not be scored at the allele level, because they lack a Thille CDS model (ALP1, ARF2A-like, RER3, and the Ole e 10-like gene), because part of the coding sequence of a haplotype was not covered (CTR1, MIK2, and PAR1 in Unnamed Lowland, and LIP1 in the principal Hass pseudohaplotype), or because no Puebla haplotype 2 sequence was placed (ZFP2 and FBD/RNI-like).

### Alleles Shared by the A-Type Cultivars

Six genes carried an allele present in all three A-type cultivars (Gwen, Hass, and Puebla) and absent from both Thille haplotypes: SDMYB (SDMYB^b^), AGAMOUS-like (AGAMOUS-like^c^), CSU2 (CSU2^d^), NPR5-BOP2 (NPR5-BOP2^d^), SGRL (SGRL^d^), and LCYB (LCYB^d^) (Figure 6A; Supplementary Table S4). Neither Unnamed Lowland haplotype carried any of the six alleles, and each A-type cultivar carried the allele on one haplotype only, the configuration of the dominant SDMYB allele. NPR5-BOP2^d^ differed from NPR5-BOP2^a^ by V272A and I359V, LCYB^d^ by L17V, E22G, and D464E, and SGRL^d^ by S5Y and I101V, and CSU2^d^ differed only at synonymous positions. At the protein level, Unnamed Lowland is homozygous for the protein encoded by SGRLd (S5Y and I101V), from which its CDS differs only at synonymous positions; for the other five genes, neither haplotype of Unnamed Lowland encodes the A-shared protein. In place of Hass read evidence, a Gwen or Puebla haplotype carried the same CDS as 53 of the 69 Hass haplotypes with a complete CDS at the 39 genes, and the same protein with synonymous differences only at 5 more (CCR1 (Thille g514)*, TIC*, VOZ1 (Thille g22280)*, VOZ1 (Thille g26844)*, and SGR1*); no Gwen or Puebla haplotype matched the Hass allele at the other 11 (both Hass haplotypes of AP2 and NRX1, and one haplotype of CCR1 (Thille g512), HY5, MIK2, NPR5-BOP2, PAR1, RPT3, and LCYB). Four of the six genes were flagged in the top 1% of type A versus type B Fst: SDMYB in both analyses (contig rank 138, scaffold rank 157), AGAMOUS-like in the contig analysis (rank 274), and CSU2 (rank 326) and NPR5-BOP2 (rank 124) in the scaffold analysis (Table 4). LCYB and SGRL were not flagged in either analysis. SPA1, the gene under the type A versus type B Fst peak of the Gwen study (Thille chromosome 1; the identifier published for SPA1 belongs to the chromosome 10 paralog), does not separate the flowering types at the haplotype level: its reference allele SPA1a is carried by both B-type genomes (Thille and Unnamed Lowland) and by the A-type Puebla and Hass, and the allele shared by Gwen and Hass (SPA1c) is absent from Puebla (Supplementary Figure S11).

### Allele Frequencies of Phased Haplotypes

Phased haplotypes of a read set usually carried similar numbers of reads, but 28 read sets at 19 genes had one haplotype at 60 to 80% of the reads over the CDS, including SDMYB in Gwen (70% SDMYB^a^ and 30% SDMYB^b^), TIC in Gwen (71%), VOZ1 (Thille g26844) in Gwen (79%), and PAR1 in Thille (74%) (Supplementary Table S5). None of these loci showed the depth expected for an additional copy on one haplotype: the depth of all reads over the gene ranged from 0.72 to 1.21 times the depth over flanks 2 to 6 kb away, against about 1.5 expected for one extra copy. PAR1 in Gwen and in Thille (0.74 and 0.72) was the only locus with reduced depth, consistent with the deletion runs reported there. Clusters of clipped reads were present in 26 of the 28 windows and did not distinguish these loci. The skewed frequencies are therefore more consistent with unequal recovery of the two haplotypes, for example fewer reads of the divergent SDMYB^b^ haplotype aligning to Thille, than with duplication of one haplotype. Of the 51 candidate genes with a UoQ gene model at the placed Hass locus, 28 had UoQ and Thille proteins at least 95% identical over at least 80% of both proteins. At the CCR1 and PAR1 cluster on chromosome 2, the UoQ models covered 19% (PAR1, MRB53_004714) and 30% (CCR1, MRB53_004716) of the Thille proteins, no UoQ model lay at the second CCR1 gene, and the closest UoQ protein elsewhere in the proteome was a paralog of 45 to 47% identity (MRB53_009075, chromosome 3). The UoQ model at MIK2 on chromosome 12 did not align to the Thille protein, and the UoQ models at MIK2 on chromosome 5, the chromosome 7 EOBI locus, SEC24A, and VOZ1 (ChrU.ver1.g447950) covered 9 to 54% of the Thille proteins at 96.7 to 100% identity. The Hass coordinates of these genes therefore rest on the DNA alignments rather than on the UoQ protein models.

### Coverage analysis

BEDTools coverage and genomecov were compared on the same alignments and produced near-identical values, differing by at most 1 at a small number of positions. Genome-wide, 63.6% of the Thille assembly fell in the medium coverage class in the Gwen alignment and 61.7% in the Thille self-alignment (Supplementary Table S3). The half class covered 161.9 Mb (18.3%) in the Gwen alignment and 215.5 Mb (24.3%) in the Thille self-alignment, and the low class covered 33.3 Mb (3.8%) and 4.6 Mb (0.5%), respectively. High and outlier coverage together covered 126.7 Mb (14.3%) in the Gwen alignment and 118.6 Mb (13.4%) in the Thille self-alignment. Per-chromosome median coverage ranged from 66x to 76x for Gwen CLR reads and from 70x to 80x for Thille HiFi reads. On the Hass (UoQ) reference (873.9 Mb across 12 chromosomes), the medium class covered 566.8 Mb (64.9%) in the Gwen alignment and 534.3 Mb (61.1%) in the Thille alignment, the half class 151.7 Mb (17.4%) and 162.1 Mb (18.6%), and the low class 40.8 Mb (4.7%) and 56.6 Mb (6.5%), respectively (Supplementary Table S3). High and outlier coverage together covered 114.6 Mb (13.1%) and 120.8 Mb (13.8%). Per-chromosome median coverage on Hass ranged from 70x to 76x for Gwen CLR reads and from 73x to 79x for Thille HiFi reads. Unnamed Lowland HiFi reads aligned to Thille had per-chromosome medians of 70x to 84x and placed 52.6% of the assembly in the medium class, 12.4% in the half class, and 18.0% in the low class, as expected for a more divergent read set (Supplementary Figure S8; Supplementary Table S3). In the Thille self-alignment, a median of 99.4% of each candidate gene span fell in the medium coverage class. Three candidate loci showed half coverage across most or all of the gene span in the Gwen alignment while remaining in the medium class in the Thille self-alignment (Supplementary Figure S4): AP2 (100% of the span in the half class), MIK2 on chromosome 12 (100%), and NAC25 (83%; the two Gwen NAC25 genes share one Thille model, g28063). SUD1 was in the half class for 55% of its span in Gwen and 99% in the Thille self-alignment. No candidate gene had 10% or more of its span in the low class in the Gwen alignment. Five candidates showed half coverage in the Thille self-alignment: SUD1 (100% of the span), VOZ1 (ChrU.ver1.g447950; 100%), Y5713 (100%), APT3 (92%), and SPA1 (58%). Eight candidates had at least a quarter of their span in the high or outlier classes in the Thille self-alignment: the CCR1 (ChrU.ver1.g414850) locus and the KCS11 locus, represented in Thille by its closest paralog g21309 because no Thille ortholog was found (100% each), MIK2 on chromosome 5 (75%), AGD5 (58%), GPAT9 (54%), SPA1-like (45%), UPL2 (40%), and EMF1 (27%). On the Hass reference, a median of 96.0% of each candidate gene span was in the medium class in the Gwen alignment and 99.2% in the Thille alignment. AP2 and NAC25 were again in the half class across their full span in Gwen (100% each) and in the medium class in Thille (99.5% and 100%). SUD1 (99.6%), APT3 (89.4%), Y5713 (73.7%), and SPA1 (50.5%) remained in the half class in the Thille reads on Hass, whereas VOZ1 (ChrU.ver1.g447950) was in the medium class (99.8%). Both Hass KCS11 gene models were in the low class across their full span in the Thille alignment and in the half class in the Gwen alignment (100% each). With respect to the Hass assembly, a coverage hit was called where a candidate gene overlapped a contiguous run of at least 1 kb in the low or half classes (deletion hit) or in the high or outlier classes (duplication hit). On the Hass reference, 10 of the 46 distinct candidate loci carried a deletion hit in the Gwen alignment and 2 a duplication hit, compared with 14 and 10 in the Thille alignment (summaries/coverage_sv_hits_hass.tsv). Deletion hits spanning the full gene in Gwen but absent in Thille were found at AP2 (run of 6,759 bp) and NAC25 (4,893 bp). The two KCS11 gene models lay within a deletion run of 91,590 bp in the Thille alignment and 41,655 bp in the Gwen alignment. PAR1 lay within a 64,156 bp deletion run in the Gwen alignment (23,587 of 40,862 bp of the gene) and a 15,198 bp run in the Thille alignment, and the adjacent CCR1 locus carried deletion runs of 11,585 and 14,683 bp in the two alignments. SUD1, Y5713, and OTU7 shared a single 20,762 bp deletion run in the Thille alignment. MIK2 on chromosome 12 carried both deletion (4,827 bp) and duplication (6,827 to 6,839 bp) runs in both alignments, and VOZ1 (ChrU.ver1.g387390) carried the only other Gwen duplication hit (10,725 of 21,439 bp). On the Thille reference, 7 of 45 distinct loci carried a deletion hit and 1 a duplication hit in the Gwen alignment, compared with 9 and 7 in the Thille self-alignment (summaries/coverage_sv_hits_thille.tsv); MIK2 on chromosome 12 lay within a 22,261 bp deletion run in Gwen, and AP2 within a 6,654 bp run.

### Tandem copies and tandem arrays

Our Fst tandem-copy search found 386 copies in 59 groups on the Thille assembly (296 copies in 48 groups on Hass). Of the 386 Thille copies, 105 (27%) were shorter than 1 kb. We split these candidates into four distinct groups by screening the 59 groups against the gene and TE annotations: TE tandem arrays, gene tandem array candidates, no tandem array, or unknown. Forty groups overlapped with no gene model and had every copy at least 95% covered by annotated TEs, and were labeled TE tandem arrays. Three further groups overlapped fewer than two gene models (no tandem array): one overlapped no gene model and no TE (ChrU.ver1.g459990) and was labeled unknown, and two overlapped a single gene model each (ChrU.ver1.g464950 and m1-g59046), and we classified this as no tandem array as well. The remaining 16 groups overlapped 64 gene models between them and we labeled as gene tandem array candidates. Pairwise alignment of these 64 models (990 alignments) supported 15 of the 16 groups, which merged into nine distinct arrays of 2 to 12 gene models, with mean lengths from 302 bp to 6.5 kb (Table 7). PAR1 was the one group not supported: its two overlapping models, g508 and g528, shared 56.7% exon identity, and g508 is a member of the CCR1 array. The KCS11 array on chromosome 6 (46.92 to 47.41 Mb) comprised 11 gene models on the minus strand, six of them full length (2.0 to 2.8 kb, with coding sequences of 1,527 to 1,566 bp and two exons) and five fragments of 354 to 450 bp; best-partner exon identities ranged from 91.5 to 99.7%, and six members had KCS11 (KCS11_ARATH) as their best Swiss-Prot hit. None of the tandem-array members was flagged with a predicted flowering function. Two arrays at 75.7 Mb on chromosome 3 interleave on opposite strands: three gene models on the plus strand whose best hit is a saccharopine dehydrogenase-like oxidoreductase (SCPDL) and two on the minus strand whose best hit is shikimate O-hydroxycinnamoyltransferase (HST), with within-array exon identities of 98.3 to 98.7%. Four arrays were retained after manual evaluation (Supplementary Figure S14): CCR1 on chromosome 2 (six gene models over 87.6 kb, from 7.91 to 8.00 Mb, five of them full length with coding sequences of 972 to 981 bp; best-partner exon identities of 90.2 to 96.1%), KCS11 on chromosome 6 (11 gene models over 483.9 kb), the leucine-rich repeat receptor kinase array containing MIK2 on chromosome 12 (nine gene models over 475.4 kb, from 46.19 to 46.66 Mb, whose best Swiss-Prot hits are Y1571, FEI1, and FEI2; best-partner exon identities of 94.9 to 100%), and the interleaved SCPDL and HST arrays on chromosome 3 (five gene models over 75.7 kb). TEs covered little or none of the coding sequence of the CCR1, KCS11, SCPDL, and HST members (0 to 18%), except g510 in the CCR1 array, a 248 bp fragment covered entirely by TEs. In the MIK2 array, by contrast, TEs covered 48 to 50% of the coding sequence of six members and all of it in the other three, including the curated MIK2 model g26459, so that this array interleaves receptor kinase genes with TE sequence.

**Table 7:**
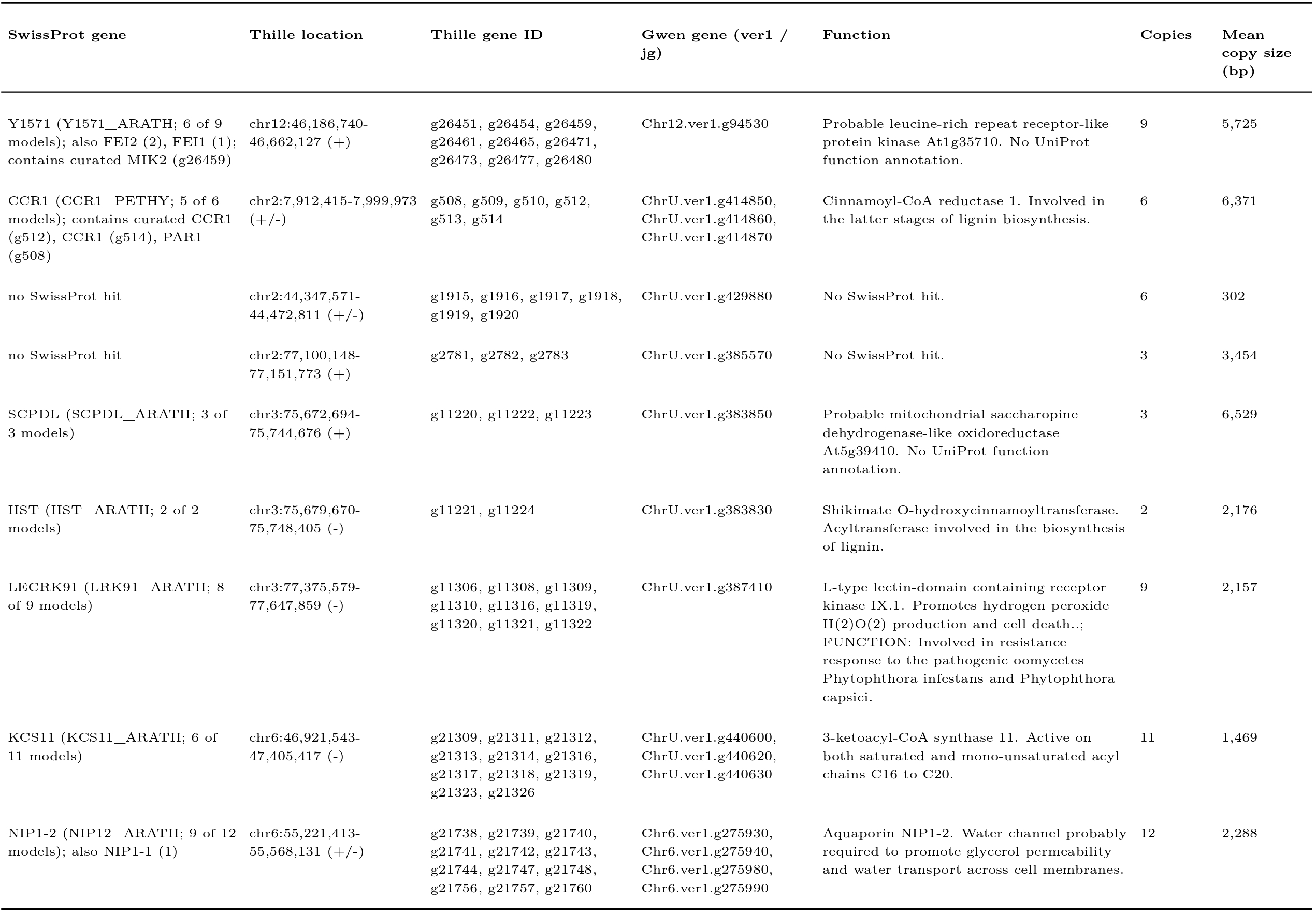
Supported tandem arrays in Thille (gene models with >= 70% pairwise exon identity) (n = 9)

## Discussion

### Genome Assembly

The Thille assembly is a primary, pseudo-haploid representation of a heterozygous genome. hifiasm resolves heterozygous sites within each contig using the HiFi reads alone, so without parental data each contig is most likely a mosaic of the two parental haplotypes rather than a single phased haplotype (Cheng et al., 2021; Rhie et al., 2020). Thille is a Hass seedling selected by J.N. Thille (Delphey, 1960), so its seed parent is Hass and its pollen parent is unknown. Gwen, a child of Thille, shows no large inversions relative to Thille, as expected since one Gwen haplotype descends from Thille, although the fragmentation of the Gwen contig assembly would also hide inversions that span contigs. For both Thille and Gwen, the pollen donor is unknown, and until they are identified a trio-based phasing approach such as trio binning (Koren et al., 2018) cannot be applied. With respect to the Merqury copy-number spectrum, we find it is consistent with the assumptions of Thille being an outcrossed individual (Rhie et al., 2020): the read-only k-mers peak at the heterozygous multiplicity (39x), as expected when k-mers unique to the alternative haplotype are absent from a primary assembly, and the k-mer completeness of 81.44% reflects the same missing haplotype rather than missing sequence of the represented one. Chromosome 12 contributed more than half of the assembly-only k-mers, and its first 7.7 Mb, 26 short contigs, form a separate block in the Omni-C map that contacts unplaced scaffolds more than the rest of the chromosome and has little matching sequence in Hass; this segment is the most likely region of residual assembly error and warrants targeted validation. The Unnamed Lowland assembly, which is telomere to telomere, carries related but divergent sequence over the first 1.2 Mb of its chromosome 12, so the segment may also be an expanded repeat region at the chromosome end rather than a misjoin. Chromosome 8 required manual curation. Its second arm had been assembled as a separate scaffold, and the 14 Mb region shared by the two scaffolds carried half the read depth on each copy and appeared as a parallel line beside the diagonal of the Omni-C map, the signatures of a false heterotype duplication (Rhie et al., 2021; Howe et al., 2021). Removing one copy and joining the arms produced a 68.4 Mb chromosome with telomeric repeats at both ends and a metacentric centromere, and raised the sequence in chromosomes from 859.8 to 885.4 Mb, BUSCO completeness from 97.3 to 99.7%, and the fraction of Omni-C pairs on chromosomes from 91.7 to 95.2%. Because the region was a heterozygous duplication, either copy could have been kept; we kept the copy in the former chromosome 8, which carries the centromere call, and the removed copy remains among the unplaced sequences. The raw reads give the inversions an independent test that the assembly comparison alone cannot. Where each genome’s own reads were available, they followed their own assembly and carried the other orientation on Thille at most inversions, so these inversions reflect sequence rather than assembly error, and the both-orientation calls in the Thille reads point to heterozygous inversions. The shared events show that Puebla and Unnamed Lowland, which belong to different flowering types, share many differences from Thille; that is not to say, however, that these differences are specific to Thille, since Hass carries about a third of them. Reference-guided scaffolds take the order of the reference, so the Thille-guided Gwen and Hass (UoQ) chromosomes cannot reveal rearrangements that fall between contigs, but the two Hass (UoQ) segments that moved coincide with rearrangements called independently by SyRI and are candidates for scaffolding errors in that assembly or for real differences, which read or Omni-C data from Hass would distinguish.

### Transposable Element Annotation and Masking

The overall repeat fraction (64.0%) and the predominance of Gypsy over Copia elements are consistent with the Gwen assembly, in which about 61% of the genome was repetitive, LTR retroelements formed roughly half of the repeat content, and more than half of these were Gypsy elements (Solares et al., 2023). Unclassified repeats nevertheless made up 23.7% of the assembly, so the class-level proportions are lower bounds for each named class.

### Coverage at Candidate flowering gene loci in Hass and Thille

In this study, we observe half coverage in Gwen at AP2, MIK2 (chromosome 12), and NAC25, with medium coverage in Thille, which is consistent with these loci being present on only one Gwen haplotype relative to Thille, through hemizygosity or haplotype divergence. Half coverage in the Thille self-alignment (SUD1, VOZ1, Y5713, APT3, and SPA1) suggests either that both Thille haplotypes of these regions are represented separately in the assembly, or that the regions are hemizygous in Thille; the Hass alignments separate these explanations. SUD1, APT3, Y5713, and SPA1 remained in the half class when the Thille reads were aligned to Hass, which is consistent with hemizygosity in Thille, whereas VOZ1 (ChrU.ver1.g447950) returned to the medium class on Hass, which is consistent with two Thille haplotypes represented separately in the Thille assembly. AP2 and NAC25 were in the half class in Gwen on both references, so their apparent hemizygosity in Gwen does not depend on the reference. The low coverage of Thille reads across the Hass KCS11 locus, within a 91.6 kb run, is consistent with the absence of a Thille ortholog of this gene. The genome-wide half class is larger in the Thille self-alignment (24.3%) than in the Gwen alignment (18.3%), which is expected where both haplotypes of a heterozygous region are present in the primary assembly and the Thille reads split between them. High and outlier coverage at the CCR1 locus and at g21309 is consistent with collapsed copies of these tandem arrays in the assembly..

### Fst Candidates

The contig and scaffold Fst analyses agree on 8 of the 26 candidate loci, and each places in its top 1% loci that the other ranks well outside it (SPA1, contig rank 162 and scaffold rank 945; AP2, scaffold rank 252 and contig rank 5,305). The two analyses differ in their reference sequence: scaffold windows may span contig joins and gaps, including those of the unplaced ChrU scaffold, whereas the contig analysis has no windows on some contigs (EMF1 and PRP8A). We therefore report both ranks rather than a single merged value, and we treat a gene as an Fst candidate when either analysis places it in the top 1%. Nine of the genes reported in the Gwen study for coverage differences also lie in top 1% scaffold windows, and six of them (RS16, Y5713, SUD1, OTU7, FPP7, and AP2) lie within 115 kb of one another, in a 160 kb stretch of ver1 chromosome 3 (15.94 to 16.10 Mb) that contains five top 1% scaffold windows; their ranks therefore reflect one differentiated region rather than six independent signals. The four genes added from the indirectly annotated set rest on annotation alone (photoperiodism, blue light signaling, and meristem identity terms), and their relationship to flowering type requires functional evidence. The candidate loci are not spread evenly across the genome. The long arm of chromosome 3 combines two regulators of the floral transition and floral organ identity (AP2 and EMF1) and FTIP7, whose rice ortholog is required for anther dehiscence (Song et al., 2018; UniProt Consortium, 2025), within 6.5 Mb, and chromosome 5 places the clock components ELF3 and TIC (Hall et al., 2003) within 1 Mb of each other, and both arrangements are shared with Hass. That is not to say, however, that these intervals are enriched for flowering genes in a statistical sense. The two intervals were delimited after the loci were plotted, and the candidate set combines genes chosen by Fst, by coverage, and by annotation, so the observed and expected counts are descriptive; a test would require intervals fixed in advance and a null that preserves the clustering of genes, for example by permuting Fst windows along the genome. Flowering type itself is determined by SDMYB on chromosome 10 (Groh et al., 2026), the chromosome on which linkage mapping had placed a QTL for flowering type (Ashworth et al., 2019), so the chromosome 3 and chromosome 5 clusters are more likely to mark linked differentiation, or genes that modify the timing of flower opening, than the locus that sets type. Since heterodichogamy is a daily rhythm (Groh et al., 2026), the proximity of ELF3 and TIC makes chromosome 5 a clear target for comparison of phased A-type and B-type haplotypes.

### Gene Identifiers of the Gwen Study Candidates

Two candidates reported in Table 2 of the Gwen study (Solares et al., 2023), EOBI and SPA1, were listed under ver1 gene identifiers that do not correspond to the genes under the type A versus type B Fst peaks. The identifiers given for EOBI (Chr5.ver1.g224860 and ChrU.ver1.g377230) resolve to other MYB genes, an EOBI-like gene and two identical copies of a MYB62-like gene, which rank at 10,870 or lower in both Fst analyses, and the identifier given for SPA1 (Chr10.ver1.g45990) resolves to the SPA1-like paralog on chromosome 10, which ranks at 16,138 or lower. Among the 944 top 1% genes, the only gene with an EOBI hit is Chr10.ver1.g56860 (contig gene jg69989), whose best contig-level match is EOBI_PETHY (best scaffold-level match MYB40_ANTMA) and which ranks 138 in the contig analysis and 157 in the scaffold analysis. This gene is SDMYB, the MYB transcription factor shown by Groh et al. (2026) to determine flowering type. Likewise, the top 1% gene with a SPA1 hit is SPA1 on chromosome 1 (Chr1.ver1.g18040; contig rank 162). The Gwen study therefore recovered the flowering-type gene among its type A versus type B Fst candidates, under the name of its closest characterized homolog, EOBI, but attached the identifiers of paralogs to it. The working gene tables of that study show how such a substitution can arise: in the scaffold type A versus type B SwissProt table, 30 of 213 rows carry a ver1 identifier that belongs to a different gene, in 29 of them the identifier of the adjacent row. We did not identify the step at which the EOBI and SPA1 identifiers were exchanged, but the corrected assignments rest on sequence, since each ver1 gene was mapped to its Gwen contig and to the Hass and Thille assemblies by alignment (Tables 4 and 6). We conclude that the EOBI candidate of the Gwen study is SDMYB, and that the Fst scan of that study had already identified the locus that controls flowering type in avocado.

### Flowering-Type Allele Classes

Beyond SDMYB, we observe five genes, AGAMOUS-like, CSU2, NPR5-BOP2, SGRL, and LCYB, in which each of the three A-type cultivars carries one copy of an allele that neither Thille haplotype carries, and Unnamed Lowland carries none of these alleles either. We report this as an observation rather than as evidence of linkage to flowering type. The six genes differ in their type A versus type B differentiation: SDMYB was flagged in the top 1% of both Fst analyses and AGAMOUS-like, CSU2, and NPR5-BOP2 in one of them, whereas LCYB and SGRL were not flagged. The pattern rests on five cultivars, three of type A and two of type B, and the same configuration arises when the A-type cultivars share ancestry that Thille and Unnamed Lowland lack; Unnamed Lowland is assigned to type B here from its SDMYB genotype rather than from a recorded flowering phenotype. AGAMOUS-like is the only gene other than SDMYB whose A-type protein allele follows the dominant model exactly, and its frameshift at the start of the coding sequence may instead reflect the use of a downstream start codon, so the protein should be confirmed from transcripts. Phased haplotypes from additional cultivars of both flowering types, Fst at these loci in larger samples, and expression of the alternative alleles will be necessary to determine whether any of these alleles follows flowering type. Allele frequencies of 60 to 80% at 19 genes, without the depth of an additional copy, more likely reflect unequal recovery of divergent haplotypes by alignment to Thille than duplications, although somatic variants in the sampled tissue cannot be excluded from leaf DNA alone. RER3 and the Ole e 10-like gene also show that a missing gene model in Thille does not mean a missing gene: RER3 is present and conserved in every assembly, and the Ole e 10-like gene appears disrupted in some haplotypes of both flowering types, so neither absence reflects flowering type. These alleles come from related cultivars. Hass is the seed parent of Thille and Thille is a parent of Gwen, so the alleles that Gwen and Hass share are better treated as one lineage than as two type A observations; Puebla is the only type A cultivar outside this pedigree. Chen et al. (2009) assigned Puebla 82% Guatemalan ancestry from 5,960 bp of sequence at four nuclear loci. In the Gwen-study PCA, built from 34,472,447 SNPs of whole-genome resequencing, Puebla instead falls about midway between the Guatemalan and Mexican accessions, with the Mexican x Guatemalan hybrids Fuerte, Zutano, Bacon, and 069-02 (Supplementary Figure S16). This difference most likely reflects the small subset of loci sequenced in 2009 compared with our whole-genome resequencing approach. The flowering type of Unnamed Lowland is predicted from its SDMYB genotype rather than observed. Within these limits, our results reaffirm SDMYB as a two-allele locus with a dominant type A allele (Groh et al., 2026): Thille and Unnamed Lowland are both homozygous for the recessive allele. We also observe five other genes with the same allelic configuration (Figure 6A) and a clustering of flowering-function genes on chromosomes 3 and 5, where TE density is lower and SV density higher than in most of the genome. We report these patterns as observations and draw no conclusions about their role in flowering type. In Juglandaceae, heterodichogamy is controlled by loci that range from a single gene to a haplotype of about 20 linked genes (Groh et al., 2025a,b), and whether any of the patterns we observe is relevant in avocado will require phased genomes from additional cultivars of both flowering types, genotyped pedigrees, and the expression of the candidate alleles in developing flowers. RER3 and the Ole e 10-like gene show that a missing gene model in Thille does not mean a missing gene. RER3 is present and conserved in every assembly, and the Ole e 10-like gene appears disrupted in haplotypes of both flowering types (Thille, Unnamed Lowland, one Hass haplotype, and both Puebla haplotypes), so neither absence reflects flowering type.

### Pigment Genes

We also examined the structural variant landscape of two pigment genes, SGRL and LCYB, and observed differences between the accessions (Figure 7C).

**Figure 7:**
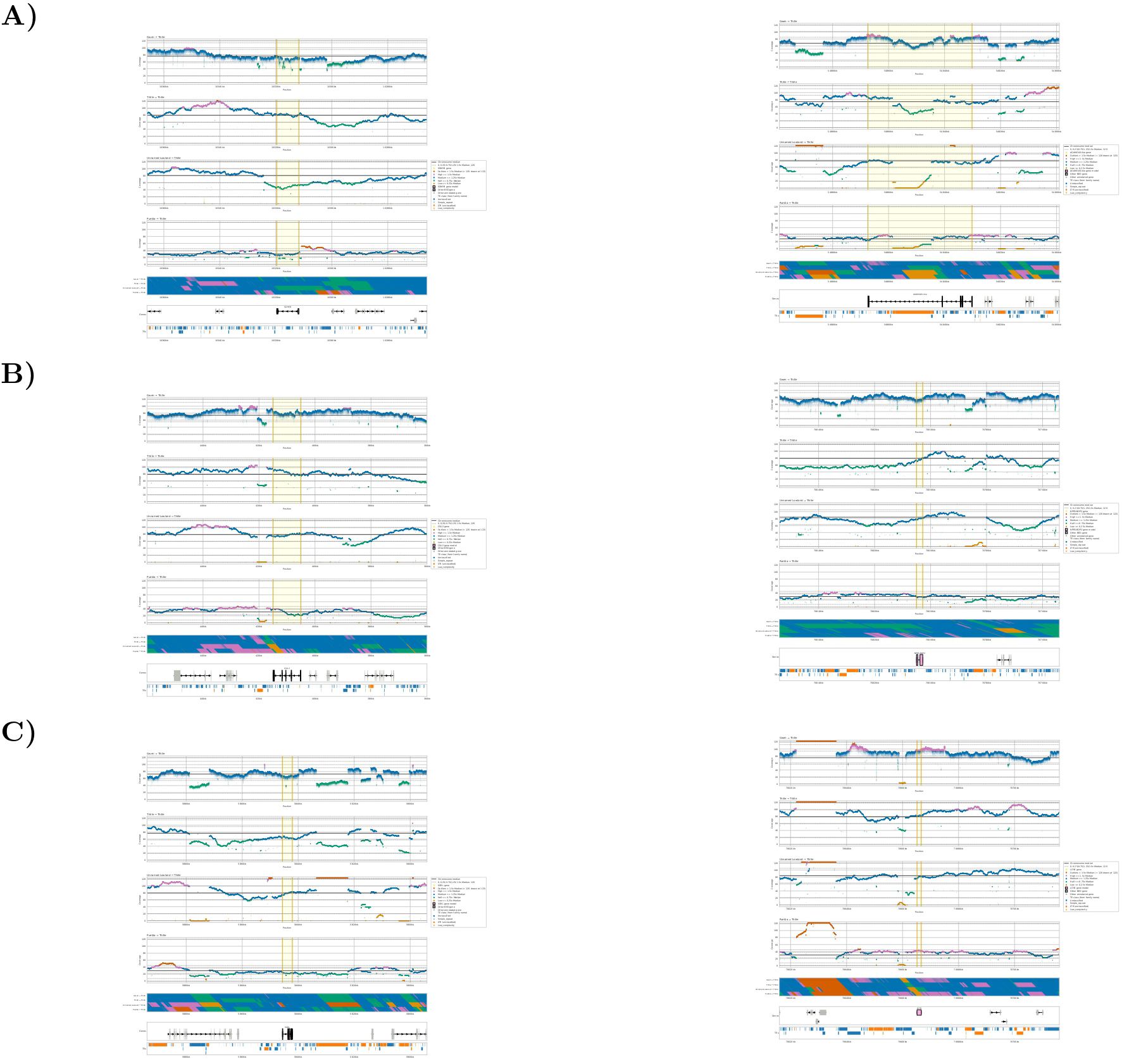
Coverage at the genes with an A-shared allele. (A-C) Gwen CLR, Thille HiFi, Unnamed Lowland HiFi, and Puebla HiFi reads on Thille (top to bottom). Read depth per position in a 100 kb window centered on the gene (shaded), colored by coverage class relative to the chromosome median (solid line); below, the class strip, gene models, and TEs. Genes on the minus strand are drawn right to left. Chromosome medians are given in that order. (A) SDMYB (g34985, chromosome 10; medians 76x, 80x, 81x, 30x) and AGAMOUS-like (g21721, chromosome 6; medians 68x, 74x, 78x, 28x). (B) CSU2 (g31874, chromosome 9; medians 74x, 78x, 79x, 31x) and NPR5-BOP2 (g6913, chromosome 1; medians 75x, 79x, 84x, 30x). (C) SGRL (g30157, chromosome 11; medians 73x, 77x, 79x, 30x) and LCYB (g6907, chromosome 1; medians 75x, 79x, 84x, 30x).

### Tandem Arrays

KCS11 encodes a 3-ketoacyl-CoA synthase, the condensing enzyme of the fatty acid elongase complex that extends acyl chains to very-long-chain fatty acids (Haslam and Kunst, 2013). Six full-length members with 91.5 to 99.7% exon identity make the chromosome 6 array a recent expansion of this family in Thille, and the absence of a one-to-one Thille ortholog of the Gwen KCS11 gene, together with the high Thille coverage over the array, suggests that copy number differs between cultivars. Whether the expansion affects flowering is not addressed by these data; in Arabidopsis, the KCS gene CER6 is required for pollen coat lipids and pollen hydration, and cer6 mutants are conditionally male sterile (Fiebig et al., 2000), which would be a testable connection to flowering type. The 40 TEs of interest were selected by Fst in the Gwen study and may mark TE insertion polymorphisms between flowering types; they are evaluated separately from the gene arrays. Of the four arrays retained, the MIK2 array differs from the others in that TEs cover half or all of the coding sequence of its members, so its copy number may reflect TE activity as much as gene duplication, and the gene models of this array should be confirmed from transcripts. The SCPDL and HST arrays interleave on opposite strands with similar within-array identities (98.3 to 98.7%), which is consistent with duplication of a block that carried both genes.

### Structural Variants

Most of the sequence that differs between the four chromosome-level assemblies is unaligned rather than rearranged, and much of it lies in pericentromeric satellite and TE sequence. SyRI calls of sequence without an alignment in these regions are therefore not deletions in the other genome, as the chromosome 3 locus illustrates: the Thille-only interval is a longer array of a satellite family that Hass and Unnamed Lowland also carry at the same position. Array lengths in any one assembly may be underestimated or collapsed, so this size difference needs read-level support, for example from ultra-long ONT reads spanning the array, before it is interpreted. The chromosome 12 inversion and the chromosome 2 translocation that distinguish Thille from Hass are specific to Hass among the four assemblies compared, so they are not features of Thille or of B-type flowering. The chromosome 10 locus lies about 1.25 Mb from SDMYB, the gene that determines flowering type (Groh et al., 2026). The block is present in Thille (type B) and Hass (type A) and absent from Unnamed Lowland (type B), so its presence does not follow flowering type, although the reduced read depth over its distal half in every read set warrants a phased examination.

## Data Availability

The Thille genome assembly and the chromosome-level assembly of Gwen scaffolded on Thille have been submitted to NCBI GenBank and are under review (BioProject PRJNA1142556); Thille HiFi reads are available under the same BioProject. Code is available at https://github.com/ESB-AI-Lab/thille-genome-report, and the annotations and read analyses are available on AvoBase. Third-party data analyzed in this study are available from NCBI: the Hass (UoQ) assembly (GCA_029852735.1) and the MH2022 haplotype assemblies of Hass (GCA_031208685.1 and GCA_031208655.1), the Puebla haplotype assemblies (GCA_060308345.1 and GCA_060308325.1) and HiFi reads (SRR36605778), the Unnamed Lowland T2T assembly (GCA_051132755.1), and the Gwen assembly and PacBio subreads (GCA_023638045.1, BioProject PRJNA758103; Solares et al. 2023), which we used together with the new chromosome-level Gwen assembly. Hass HiFi reads (SRR13510945, SRR13510946, and SRR13510947; Nath et al. 2022) are available but were not used in this study.

## Acknowledgments

This work used Expanse at the San Diego Supercomputer Center (SDSC) and Bridges-2 at the Pittsburgh Supercomputing Center (PSC) through allocation MCB180035 (Species Rescue and Rapid Low Cost Assembly of Food for Structural Variant Detection using Artificial Intelligence) from the Advanced Cyberinfrastructure Coordination Ecosystem: Services & Support (ACCESS) program (Boerner et al., 2023), which is supported by U.S. National Science Foundation grants #2138259, #2138286, #2138307, #2137603 and #2138296. E.A.S. thanks Dr. Brandon Gaut for all his mentorship, support, help, and always willing to lend an ear, and Dr. Pavel Pevzner for his mentorship and guidance while junior teaching faculty at UC San Diego.

