## Supplementary material for "Thille, a chromosome-level type B avocado genome, and the structural and allelic variation at candidate flowering-type loci": Supplmentary Figures

[*]Contributed significantly to the analysis of this study under the mentorship of Dr. Solares.

### Supplementary Tables

**Supplementary Table S2:** Top 1% Fst genes with indirect flowering annotation, not added to the candidate flowering genes (n = 8 Gwen gene models, 6 genes).

| SwissProt gene | Thille location | Thille gene ID | Gwen gene (ver1 / jg) | Function | Flowering-related annotation | Contig Fst rank (mean Fst) | Scaffold Fst rank (mean Fst) | Top 1% Fst set |
|---|---|---|---|---|---|---|---|---|
| CIGR2 (CIGR2_ORYSJ) | chr3:73,340,679-73,342,448 (+) | g11118 | ChrU.ver1.g434090 / jg40771 | Chitin-inducible gibberellin-responsive protein 2. May play a regulatory role in the early step of oligosaccharide elicitor response, downstream of the membrane-associated high-affinity chitin-binding protein. | protein name: Chitin-inducible gibberellin-responsive protein 2 | 330 (0.210) | 4,593 (0.094) | contig |
| LIP1 (LIP1_ARATH) | chr3:91,603,237-91,615,352 (+) | g12182 | Chr3.ver1.g151770 / jg115659 | Triacylglycerol lipase 1. Triacylglycerol (TAG) lipase active on triolein, trioctanoin, tributyrin and 1,3-Diolein, but not on phospho- and galactolipids (PubMed:16226259). | GO: response to gibberellin | 245 (0.230) | 867 (0.167) | contig |
| CCCH zinc finger: ENDO4 (ENDO4_ARATH) | chr4:18,183,255-18,203,047 (-) | g28093 | ChrU.ver1.g423570 / jg100119 | Endonuclease 4. Endonuclease that can use single-stranded RNA and DNA as substrates (PubMed:23620482). | Blast2GO: flower development | 709 (0.169) | 400 (0.213) | scaffold |
| CTR1 (CTR1_ARATH) | chr6:23,174,818-23,209,327 (+) | g20795 | ChrU.ver1.g468080 / jg104369 | Serine/threonine-protein kinase CTR1. Acts as a negative regulator in the ethylene response pathway (PubMed:8431946). | GO: gibberellin biosynthetic process | 446 (0.194) | 828 (0.170) | contig |
| Ole e 10-like (ALL10_OLEEU) | chr9:32,021,211-32,022,691 (-); aligned locus | none (unannotated) | Chr9.ver1.g346450 / jg57424 | Major pollen allergen Ole e 10. Carbohydrate-binding protein binding preferentially 1,3-beta-glucans. | protein name: Major pollen allergen Ole e 10 (allergen Ole e 10) | 203 (0.247) | 564 (0.189) | contig |
| ZEP (ZEP_ORYSJ) | chr11:10,926,650-10,941,480 (+) | g30489 | Chr11.ver1.g78900 / jg31797 | Zeaxanthin epoxidase, chloroplastic. Zeaxanthin epoxidase that plays an important role in the xanthophyll cycle and abscisic acid (ABA) biosynthesis. | protein name: abscisic acid (ABA) biosynthesis. Converts zeaxanthin into antheraxanthin and subsequently violaxanthin. Required for resista | 286 (0.219) | 1,178 (0.151) | contig |
| ZEP (ZEP_ORYSJ) | chr11:10,926,650-10,941,480 (+) | g30489 | Chr11.ver1.g78910 / jg31797 | Zeaxanthin epoxidase, chloroplastic. Zeaxanthin epoxidase that plays an important role in the xanthophyll cycle and abscisic acid (ABA) biosynthesis. | protein name: abscisic acid (ABA) biosynthesis. Converts zeaxanthin into antheraxanthin and subsequently violaxanthin. Required for resista | 7,145 (0.074) | 1,178 (0.151) | neither |

| SwissProt gene | Thille location | Thille gene ID | Gwen gene (ver1 / jg) | Function | Flowering-related annotation | Contig Fst rank (mean Fst) | Scaffold Fst rank (mean Fst) | Top 1% Fst set |
|---|---|---|---|---|---|---|---|---|
| ZEP (ZEP_ORYSJ) | chr11:10,926,650-10,941,480 (+) | g30489 | Chr11.ver1.g78920 / jg31797 | Zeaxanthin epoxidase, chloroplastic. Zeaxanthin epoxidase that plays an important role in the xanthophyll cycle and abscisic acid (ABA) biosynthesis. | protein name: abscisic acid (ABA) biosynthesis. Converts zeaxanthin into antheraxanthin and subsequently violaxanthin. Required for resista | 286 (0.219) | 1,178 (0.151) | contig |

### Supplementary Figures

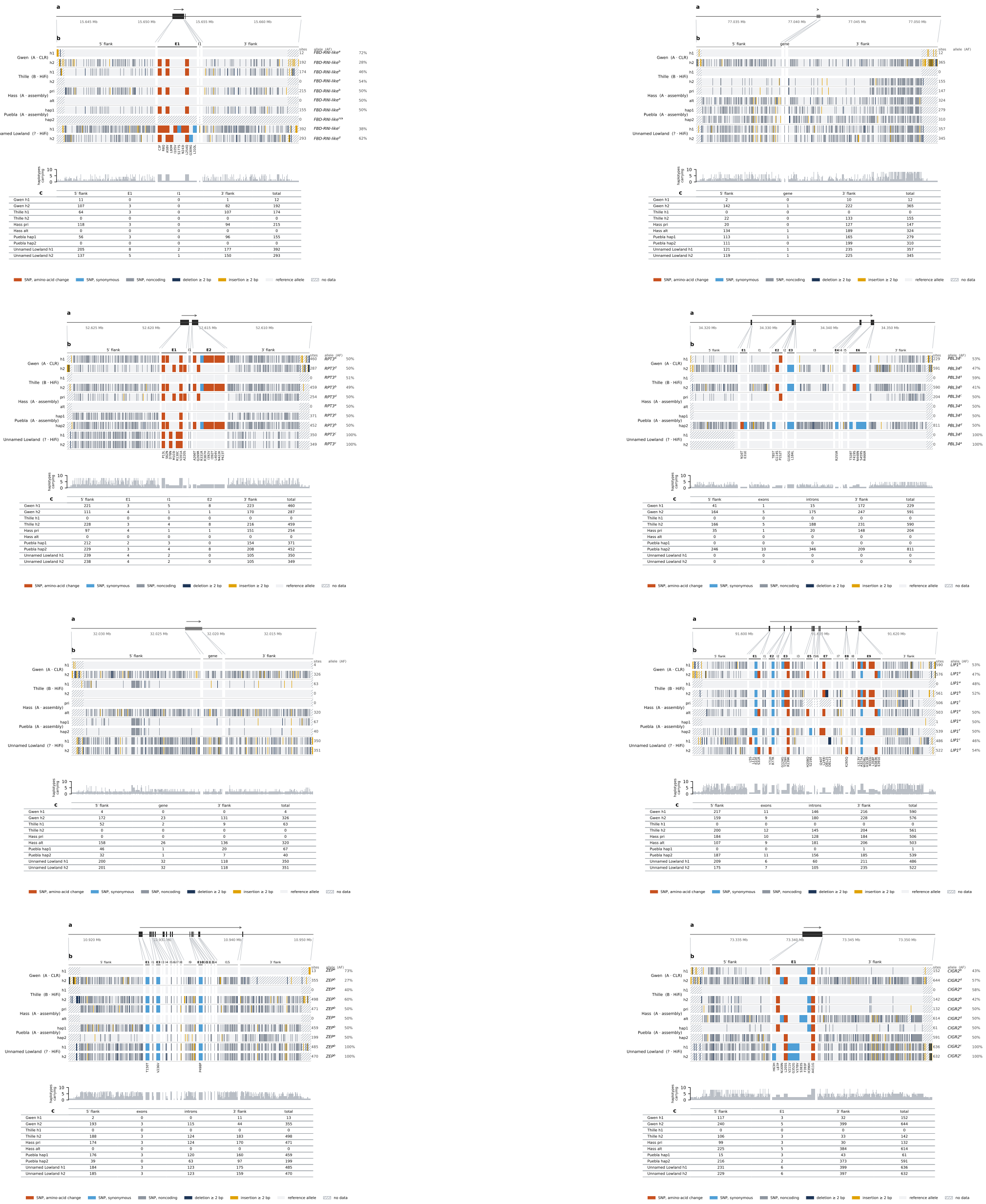

**Supplementary Figures S5 and S6:** S5: Allele matrices of the ten genes recovered by the combined Fst screen. Drawn as in Figure 6B, left to right: FBD-RNI-like (g32666, chromosome 9); RER3 (no Thille CDS model, chromosome 2); RPT3 (g2025, chromosome 2); PBL34 (g6305, chromosome 1); Ole-e-10-like (no Thille CDS model, chromosome 9); LIP1 (g12182, chromosome 3); ZEP (g30489, chromosome 11); CIGR2 (g11118, chromosome 3); CCCH-zinc-finger (g28093, chromosome 4); CTR1 (g20795, chromosome 6). S6: Coverage at the ten genes recovered by the combined Fst screen. Drawn as in Figure 7, on Thille (first ten panels; Gwen, Thille, Unnamed Lowland, Puebla) and on Unnamed Lowland (last ten; Gwen, Thille, Unnamed Lowland). On Thille: FBD-RNI-like (g32666, chromosome 9; medians 74x, 78x, 79x, 31x); RER3 (no Thille CDS model, chromosome 2; medians 73x, 77x, 80x, 30x); RPT3 (g2025, chromosome 2; medians 73x, 77x, 80x, 30x); PBL34 (g6305, chromosome 1; medians 75x, 79x, 84x, 30x); Ole-e-10-like (no Thille CDS model, chromosome 9; medians 74x, 78x, 79x, 31x); LIP1 (g12182, chromosome 3; medians 74x, 79x, 83x, 30x); ZEP (g30489, chromosome 11; medians 73x, 77x, 79x, 30x); CIGR2 (g11118, chromosome 3; medians 74x, 79x, 83x, 30x); CCCH-zinc-finger (g28093, chromosome 4; medians 67x, 70x, 70x, 27x); CTR1 (g20795, chromosome 6; medians 68x, 74x, 78x, 28x). On Unnamed Lowland: FBD-RNI-like (Thille g32666, chromosome 9; medians 73x, 77x, 83x); RER3 (no Thille CDS model, chromosome 2; medians 72x, 76x, 84x); RPT3 (Thille g2025, chromosome 2; medians 72x, 76x, 84x); PBL34 (Thille g6305, chromosome 1; medians 76x, 80x, 85x); Ole-e-10-like (no Thille CDS model, chromosome 9; medians 73x, 77x, 83x); LIP1 (Thille g12182, chromosome 3; medians 75x, 79x, 84x); ZEP (Thille g30489, chromosome 11; medians 72x, 76x, 83x); CIGR2 (Thille g11118, chromosome 3; medians 75x, 79x, 84x); CCCH-zinc-finger (Thille g28093, chromosome 4; medians 69x, 72x, 80x); CTR1 (Thille g20795, chromosome 6; medians 70x, 77x, 83x).

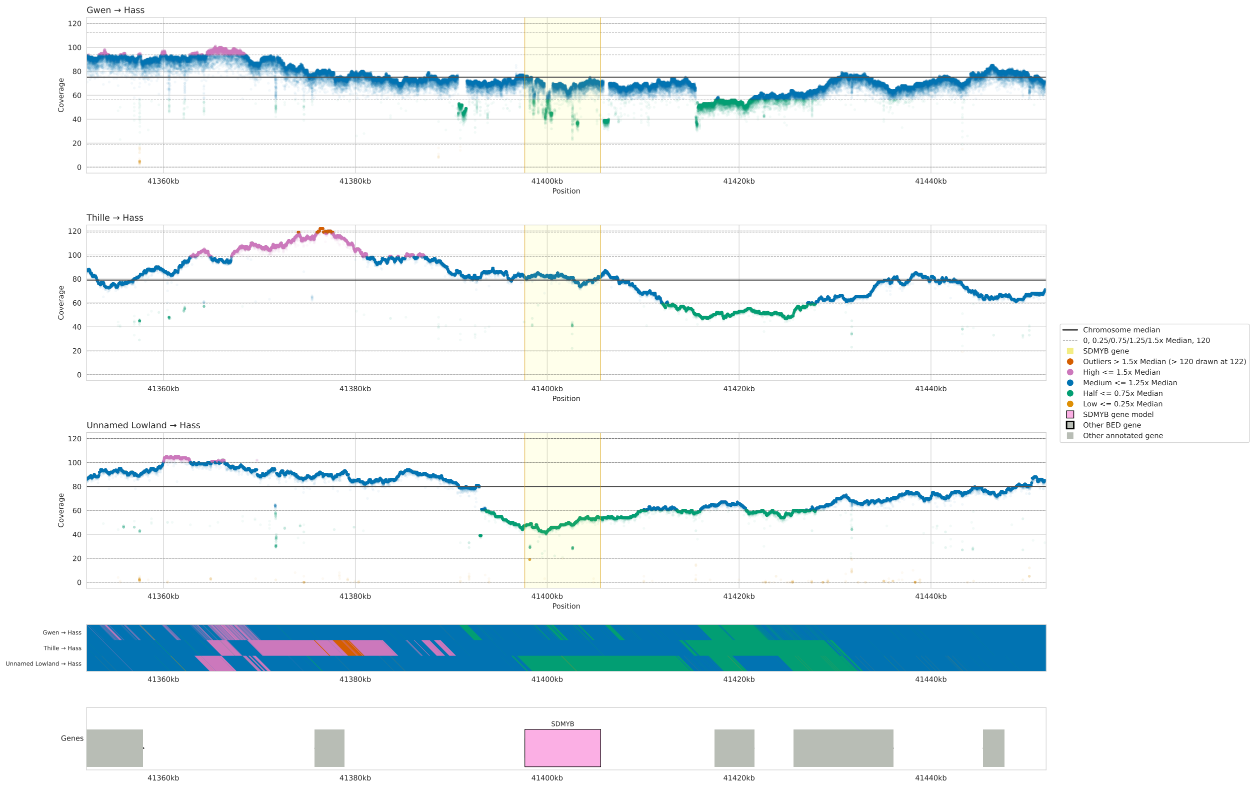


**Supplementary Figure S7:** Read coverage at SDMYB on the Hass (UoQ) assembly. Gwen CLR, Thille HiFi, and Unnamed Lowland HiFi reads, drawn as in Figure 7; SDMYB (Thille g34985, chromosome 10; medians 75x, 79x, 80x). Hass reads are not shown on their own assembly because the HiFi reads available from NCBI are a fraction of those reported (Nath et al., 2022).

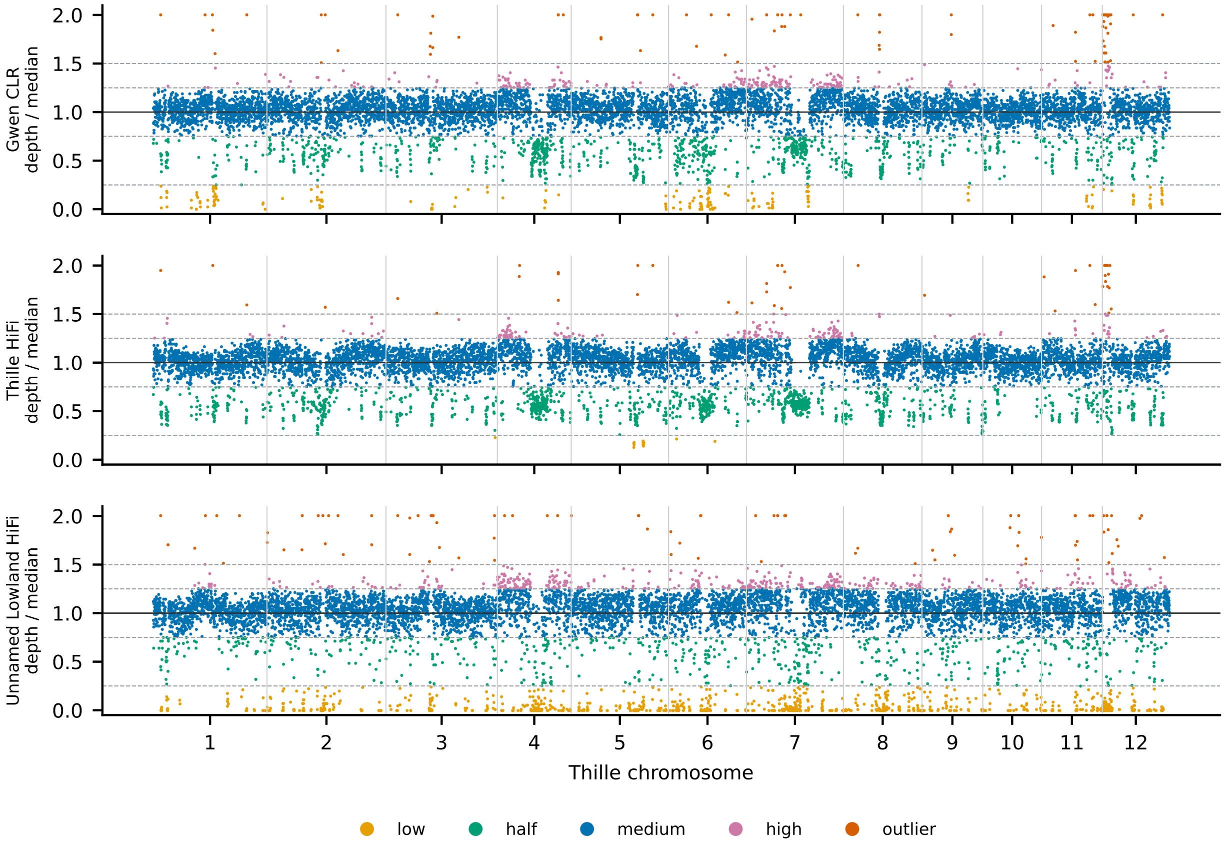


**Supplementary Figure S8:** Genome-wide read coverage on Thille. Median depth of 100 kb bins divided by the chromosome median for Gwen CLR, Thille HiFi, and Unnamed Lowland HiFi reads; dashed lines mark the class boundaries (0.25, 0.75, 1.25, and 1.5) and values above 2.0 are drawn at 2.0.

**A)**

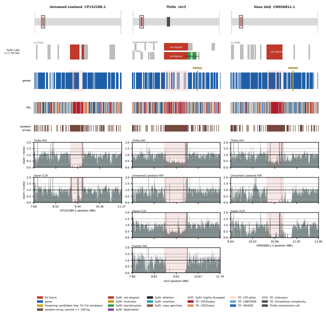

**B)**

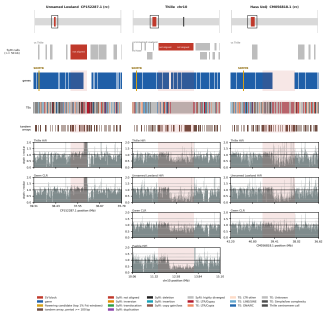

**Supplementary Figure S9:** The chromosome 3 and chromosome 10 loci in Unnamed Lowland, Thille, and Hass (UoQ). (A-B) One column per genome: the chromosome with the window boxed, SyRI events of at least 50 kb, genes with the flowering candidates named, TEs, tandem arrays, and read depth relative to the chromosome median in 10 kb bins; rc marks chromosomes drawn on the Thille strand. (A) The chromosome 3 satellite array (Thille 9.10 to 10.20 Mb, 1.10 Mb), shorter in Hass (0.76 Mb) and Unnamed Lowland (0.57 Mb). Medians on Thille: Thille HiFi 79x, Unnamed Lowland HiFi 83x, Gwen CLR 74x, Puebla HiFi 30x. (B) The chromosome 10 block shared by Thille and Hass (Thille 11.56 to 13.60 Mb) and replaced by 0.66 Mb of unaligned sequence in Unnamed Lowland; over its distal half, Unnamed Lowland and Puebla reads fall to 0.40x and 0.44x on Thille. Medians on Thille: Thille HiFi 80x, Unnamed Lowland HiFi 81x, Gwen CLR 76x, Puebla HiFi 30x.

**A)**

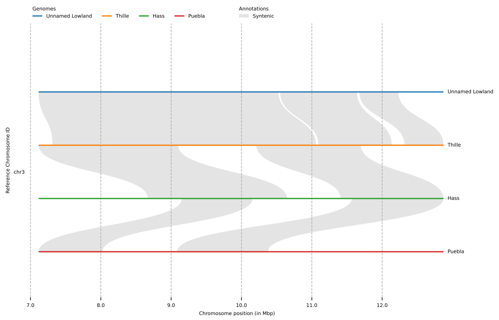


**B)**

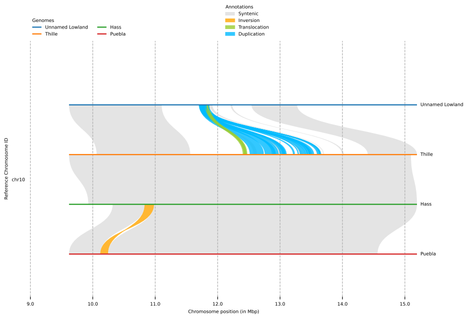


**Supplementary Figure S10:** Four-genome synteny at the chromosome 3 and chromosome 10 loci. (A-B) SyRI ribbons (plotsr, events of at least 10 kb) of Unnamed Lowland, Thille, Hass, and Puebla, zoomed on the Thille window. (A) Chromosome 3, Thille 7.60 to 11.70 Mb. (B) Chromosome 10, Thille 10.06 to 15.10 Mb.

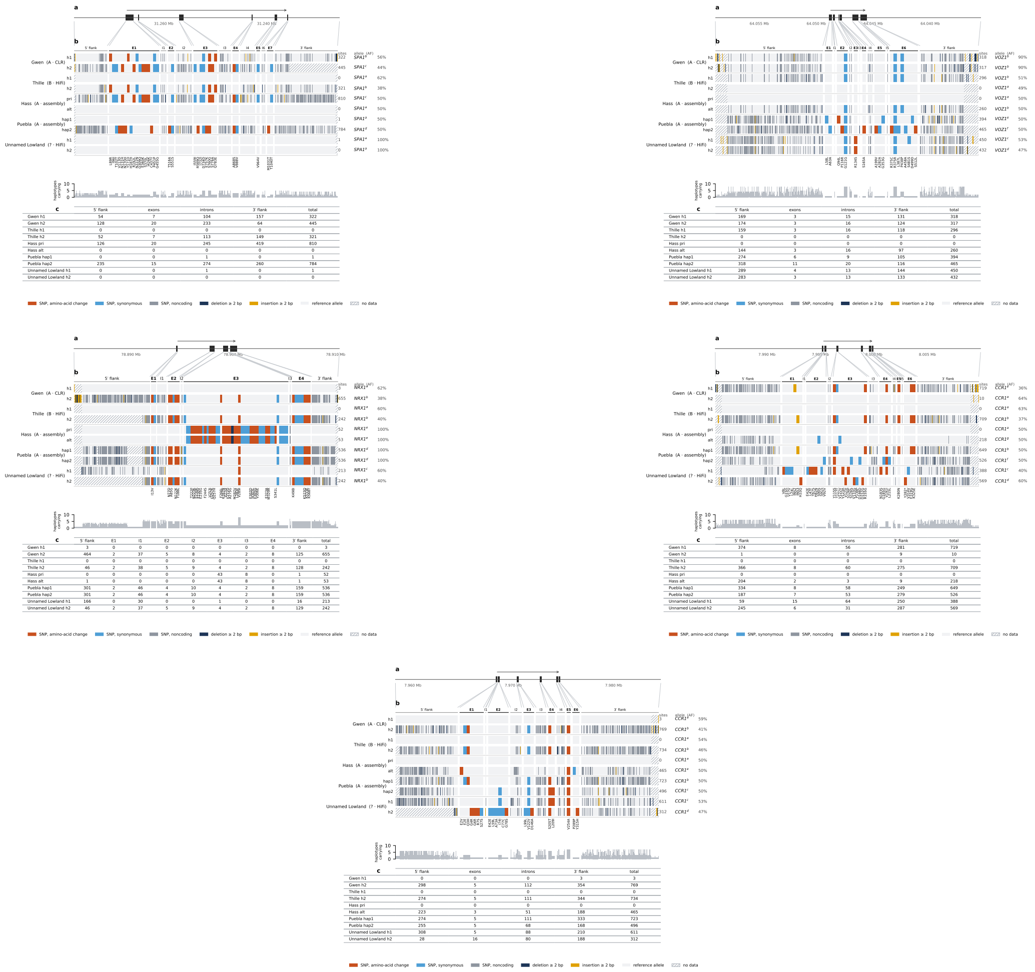

**Supplementary Figure S11:** Allele matrices of the other candidate flowering genes. Drawn as in Figure 6B, left to right: SPA1 (g6249, chromosome 1); VOZ1 (g22280, chromosome 6); NRX1 (g7314, chromosome 1); CCR1 (g514, chromosome 2); CCR1 (g512, chromosome 2).

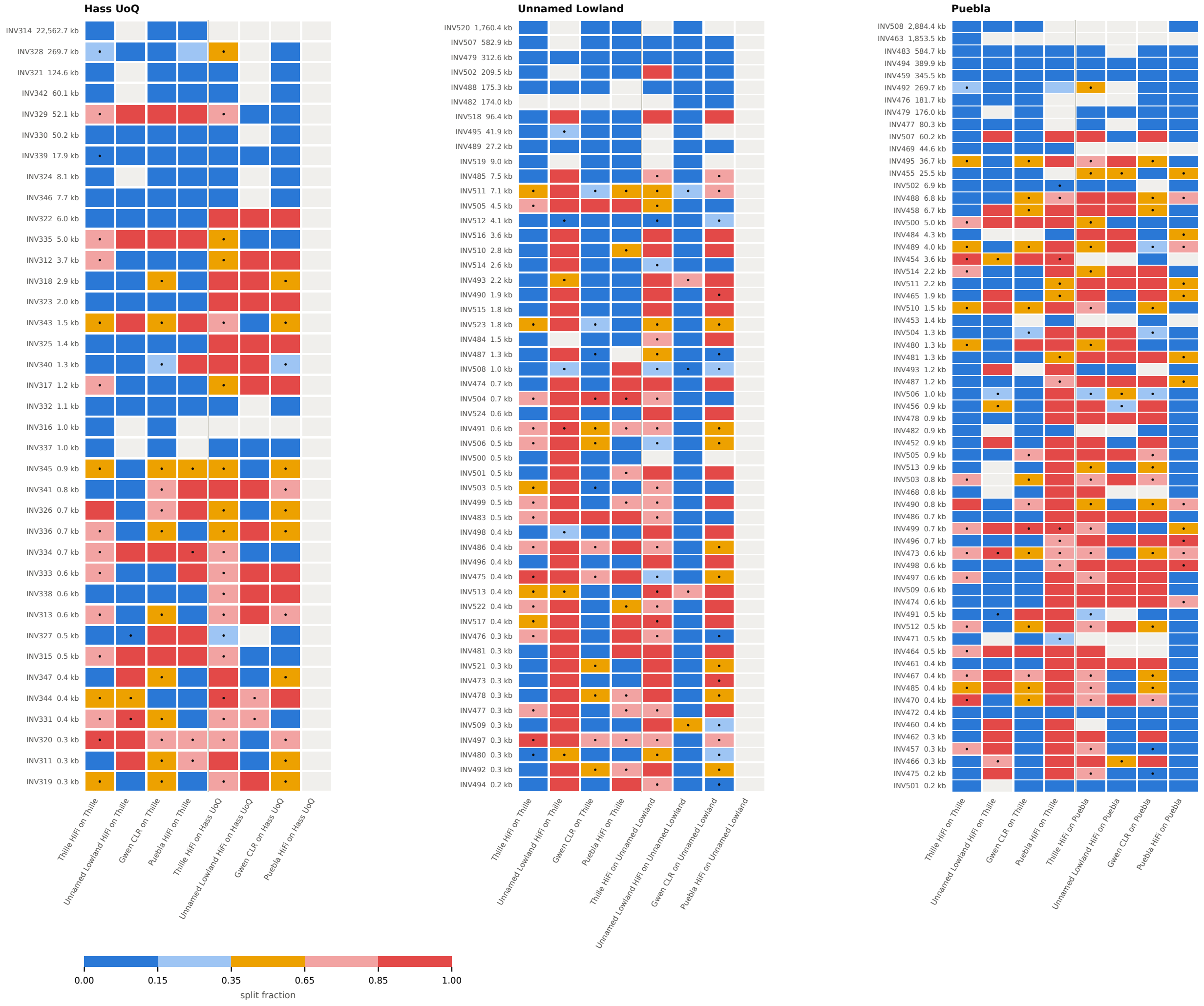


**Supplementary Figure S12:** Read support at the 152 inversions longer than 100 bp. One page per inversion (37 against Hass, 52 against Unnamed Lowland, 63 against Puebla), named by its SyRI identifier, and an overview. Each page shows 100 kb windows on Thille and on the other assembly with the read depth of Thille HiFi, Unnamed Lowland HiFi, Gwen CLR, and Puebla HiFi reads relative to the chromosome median, clipped and split reads at the breakpoints, flowering genes, gene models, and TEs; read counts per inversion are in the inversion BED files. The overview gives the fraction of split reads per inversion and read set; dots mark inversions with both orientations present.

**A)**

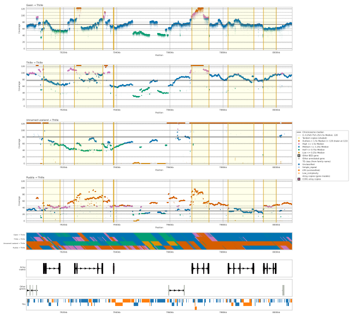

**B)**

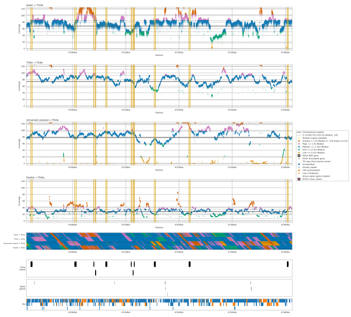

**C)**

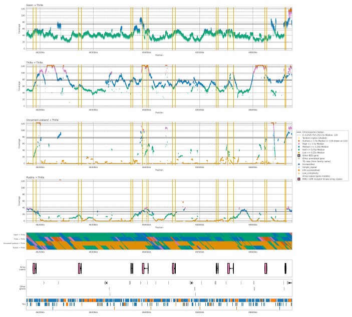

**D)**

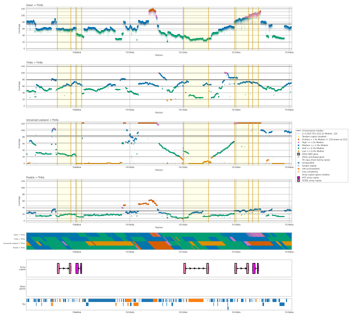

**Supplementary Figure S14:** Selected tandem arrays. (A-D) Coverage on Thille drawn as in Figure 7, in the smallest fixed window that holds the array, with the array copies shaded and drawn as a separate track. (A) CCR1, chromosome 2, 100 kb (6 copies over 87,559 bp; medians 73x, 77x, 80x, 30x). (B) KCS11, chromosome 6, 500 kb (11 copies over 483,875 bp; medians 68x, 74x, 78x, 28x). (C) The leucine-rich repeat receptor kinase array containing MIK2, chromosome 12, 500 kb (9 copies over 475,388 bp; medians 74x, 78x, 77x, 28x). (D) The interleaved SCPDL (pink) and HST (fuchsia) arrays, chromosome 3, 100 kb (5 copies over 75,712 bp; medians 74x, 79x, 83x, 30x).

**A)**

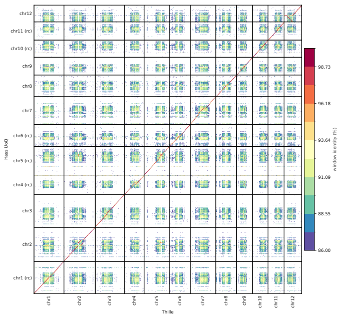

**B)**

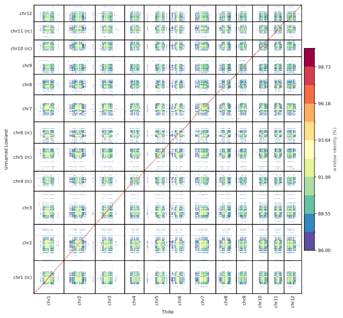

**C)**

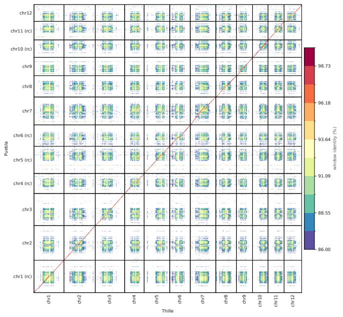

**Supplementary Figure S15:** Whole-genome ModDotPlot comparisons of Thille with Hass (UoQ), Unnamed Lowland, and Puebla. (A-C) ModDotPlot v0.9.9 (k = 21) of the Thille chromosomes (x axis) against each assembly (y axis), joined in chromosome order; window identity of at least 86%, and rc marks chromosomes reverse-complemented to the Thille strand. The SyRI dot plots of the same comparisons are in Figure 5A. (A) Hass (UoQ), 875 kb windows. (B) Unnamed Lowland, 843 kb windows. (C) Puebla, 836 kb windows.

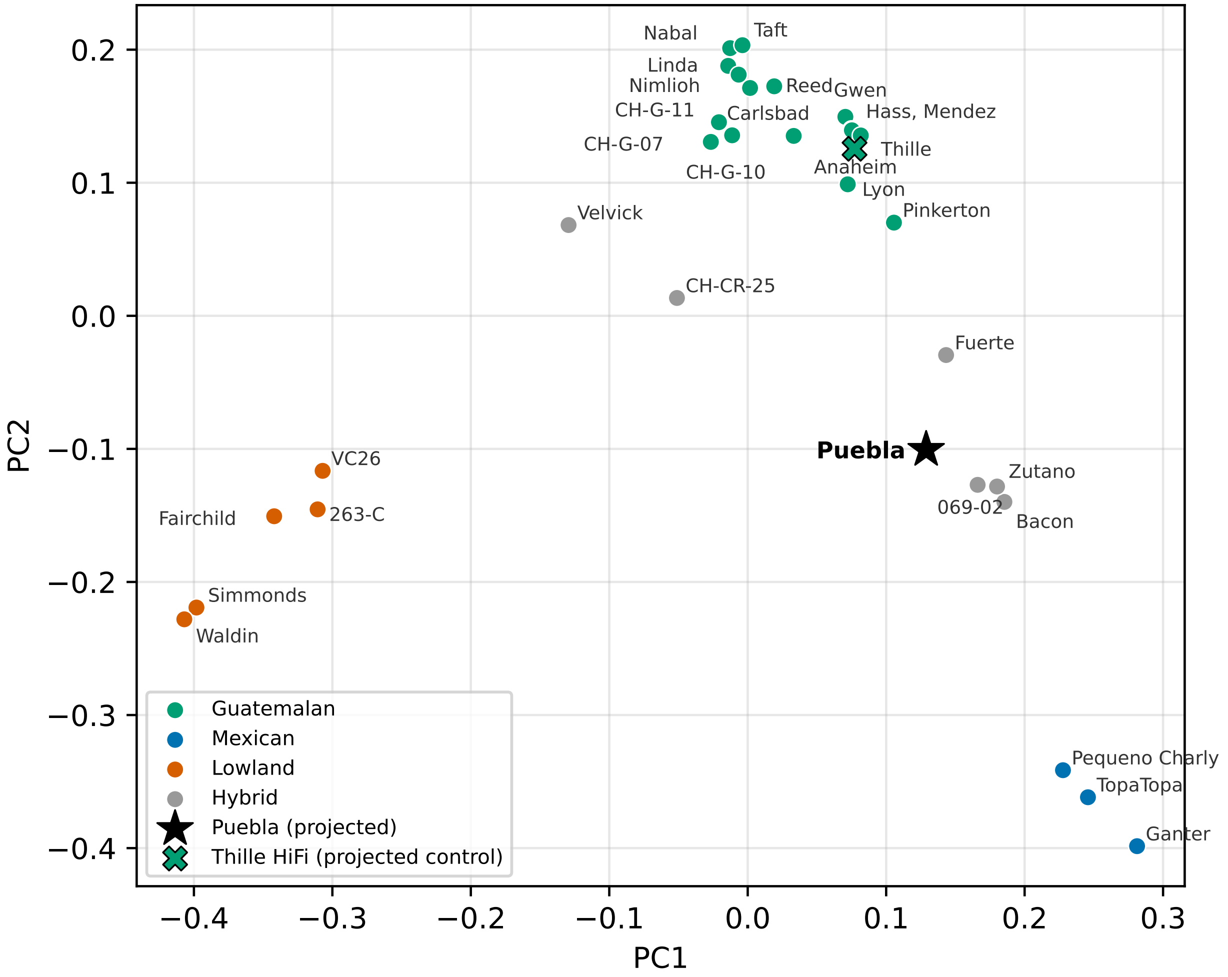


**Supplementary Figure S16:** Puebla projected onto the principal component analysis of the Gwen study. PC1 versus PC2 of the 30 resequenced accessions of Solares et al. (2023), colored by botanical group (Guatemalan, Mexican, Lowland, or hybrid, from their admixture analysis at K = 3), with Puebla HiFi reads projected onto the same axes (star) and Thille HiFi reads projected as a control (cross), which falls on the Illumina position of Thille.

**Supplementary Table S3:** Coverage classes of Gwen CLR, Thille HiFi, and Unnamed Lowland HiFi reads on Thille, per chromosome and genome-wide.

| Read set | Chromosome | Median depth | Length (bp) | Low bp (%) | Half bp (%) | Medium bp (%) | High bp (%) | Outlier bp (%) |
|---|---|---|---|---|---|---|---|---|
| Gwen CLR | chr1 | 75 | 99,146,049 | 5,995,789 (6.0) | 11,559,602 (11.7) | 70,231,363 (70.8) | 8,328,568 (8.4) | 3,030,727 (3.1) |
| Gwen CLR | chr2 | 73 | 103,667,863 | 3,676,253 (3.5) | 19,918,472 (19.2) | 67,083,058 (64.7) | 9,666,849 (9.3) | 3,323,231 (3.2) |
| Gwen CLR | chr3 | 74 | 97,080,987 | 2,315,665 (2.4) | 14,125,405 (14.6) | 68,785,967 (70.9) | 8,492,197 (8.7) | 3,361,753 (3.5) |
| Gwen CLR | chr4 | 67 | 64,330,002 | 1,591,974 (2.5) | 17,643,555 (27.4) | 32,016,277 (49.8) | 9,359,088 (14.5) | 3,719,108 (5.8) |
| Gwen CLR | chr5 | 73 | 84,859,498 | 1,142,270 (1.3) | 15,436,619 (18.2) | 58,271,145 (68.7) | 7,376,710 (8.7) | 2,632,754 (3.1) |
| Gwen CLR | chr6 | 68 | 67,830,436 | 7,542,861 (11.1) | 13,886,473 (20.5) | 34,816,869 (51.3) | 8,536,189 (12.6) | 3,048,044 (4.5) |
| Gwen CLR | chr7 | 66 | 84,521,855 | 5,109,093 (6.0) | 22,300,528 (26.4) | 38,566,060 (45.6) | 13,299,951 (15.7) | 5,246,223 (6.2) |
| Gwen CLR | chr8 | 73 | 68,384,694 | 473,466 (0.7) | 13,890,370 (20.3) | 45,224,286 (66.1) | 6,030,570 (8.8) | 2,766,002 (4.0) |
| Gwen CLR | chr9 | 74 | 53,055,096 | 793,080 (1.5) | 8,275,865 (15.6) | 37,386,275 (70.5) | 4,718,520 (8.9) | 1,881,356 (3.5) |
| Gwen CLR | chr10 | 76 | 51,141,787 | 252,373 (0.5) | 6,489,699 (12.7) | 39,301,045 (76.8) | 3,399,790 (6.6) | 1,698,880 (3.3) |
| Gwen CLR | chr11 | 73 | 53,004,496 | 1,312,780 (2.5) | 8,615,361 (16.3) | 36,486,785 (68.8) | 4,209,716 (7.9) | 2,379,854 (4.5) |
| Gwen CLR | chr12 | 74 | 58,335,637 | 3,060,480 (5.2) | 9,776,887 (16.8) | 35,349,642 (60.6) | 4,807,367 (8.2) | 5,341,261 (9.2) |
| Gwen CLR | genome | 66 to 76 | 885,358,400 | 33,266,084 (3.8) | 161,918,836 (18.3) | 563,518,772 (63.6) | 88,225,515 (10.0) | 38,429,193 (4.3) |
| Thille HiFi | chr1 | 79 | 99,146,049 | 205,642 (0.2) | 18,666,722 (18.8) | 69,913,067 (70.5) | 8,054,224 (8.1) | 2,306,394 (2.3) |
| Thille HiFi | chr2 | 77 | 103,667,863 | 426,931 (0.4) | 26,098,971 (25.2) | 64,790,754 (62.5) | 9,814,023 (9.5) | 2,537,184 (2.4) |
| Thille HiFi | chr3 | 79 | 97,080,987 | 249,755 (0.3) | 19,083,901 (19.7) | 67,769,073 (69.8) | 7,619,788 (7.8) | 2,358,470 (2.4) |
| Thille HiFi | chr4 | 70 | 64,330,002 | 235,348 (0.4) | 21,958,106 (34.1) | 28,734,406 (44.7) | 10,240,925 (15.9) | 3,161,217 (4.9) |
| Thille HiFi | chr5 | 77 | 84,859,498 | 1,428,881 (1.7) | 18,931,171 (22.3) | 54,846,465 (64.6) | 7,410,168 (8.7) | 2,242,813 (2.6) |
| Thille HiFi | chr6 | 74 | 67,830,436 | 507,563 (0.7) | 19,822,219 (29.2) | 37,094,626 (54.7) | 8,319,020 (12.3) | 2,087,008 (3.1) |
| Thille HiFi | chr7 | 70 | 84,521,855 | 213,426 (0.3) | 29,696,698 (35.1) | 36,419,157 (43.1) | 13,751,940 (16.3) | 4,440,634 (5.3) |

| Read set | Chromosome | Median depth | Length (bp) | Low bp (%) | Half bp (%) | Medium bp (%) | High bp (%) | Outlier bp (%) |
|---|---|---|---|---|---|---|---|---|
| Thille HiFi | chr8 | 76 | 68,384,694 | 185,820 (0.3) | 17,781,350 (26.0) | 42,327,331 (61.9) | 6,258,785 (9.2) | 1,831,408 (2.7) |
| Thille HiFi | chr9 | 78 | 53,055,096 | 228,839 (0.4) | 11,110,076 (20.9) | 35,403,321 (66.7) | 4,592,466 (8.7) | 1,720,394 (3.2) |
| Thille HiFi | chr10 | 80 | 51,141,787 | 117,585 (0.2) | 7,531,490 (14.7) | 39,012,211 (76.3) | 3,240,655 (6.3) | 1,239,846 (2.4) |
| Thille HiFi | chr11 | 77 | 53,004,496 | 113,524 (0.2) | 11,524,420 (21.7) | 34,723,082 (65.5) | 4,703,129 (8.9) | 1,940,341 (3.7) |
| Thille HiFi | chr12 | 78 | 58,335,637 | 731,881 (1.3) | 13,344,786 (22.9) | 35,556,965 (61.0) | 4,967,290 (8.5) | 3,734,715 (6.4) |
| Thille HiFi | genome | 70 to 80 | 885,358,400 | 4,645,195 (0.5) | 215,549,910 (24.3) | 546,590,458 (61.7) | 88,972,413 (10.0) | 29,600,424 (3.3) |
| Unnamed Lowland HiFi | chr1 | 84 | 99,146,049 | 6,490,559 (6.5) | 13,134,933 (13.2) | 68,304,063 (68.9) | 6,294,237 (6.3) | 4,922,257 (5.0) |
| Unnamed Lowland HiFi | chr2 | 80 | 103,667,863 | 16,940,726 (16.3) | 13,330,950 (12.9) | 59,239,604 (57.1) | 8,688,210 (8.4) | 5,468,373 (5.3) |
| Unnamed Lowland HiFi | chr3 | 83 | 97,080,987 | 11,291,790 (11.6) | 12,563,166 (12.9) | 60,234,141 (62.0) | 6,868,990 (7.1) | 6,122,900 (6.3) |
| Unnamed Lowland HiFi | chr4 | 70 | 64,330,002 | 17,781,706 (27.6) | 7,162,586 (11.1) | 22,218,675 (34.5) | 12,114,497 (18.8) | 5,052,538 (7.9) |
| Unnamed Lowland HiFi | chr5 | 79 | 84,859,498 | 16,873,735 (19.9) | 9,929,042 (11.7) | 45,006,177 (53.0) | 8,903,230 (10.5) | 4,147,314 (4.9) |
| Unnamed Lowland HiFi | chr6 | 78 | 67,830,436 | 13,899,702 (20.5) | 8,089,587 (11.9) | 33,784,662 (49.8) | 7,814,550 (11.5) | 4,241,935 (6.3) |
| Unnamed Lowland HiFi | chr7 | 72 | 84,521,855 | 21,957,858 (26.0) | 10,289,190 (12.2) | 32,213,195 (38.1) | 14,287,488 (16.9) | 5,774,124 (6.8) |
| Unnamed Lowland HiFi | chr8 | 75 | 68,384,694 | 15,658,884 (22.9) | 7,890,530 (11.5) | 31,501,615 (46.1) | 9,333,077 (13.6) | 4,000,588 (5.9) |
| Unnamed Lowland HiFi | chr9 | 79 | 53,055,096 | 8,975,596 (16.9) | 7,002,522 (13.2) | 28,636,614 (54.0) | 5,423,775 (10.2) | 3,016,589 (5.7) |
| Unnamed Lowland HiFi | chr10 | 81 | 51,141,787 | 6,768,558 (13.2) | 7,046,092 (13.8) | 29,363,542 (57.4) | 4,363,252 (8.5) | 3,600,343 (7.0) |
| Unnamed Lowland HiFi | chr11 | 79 | 53,004,496 | 8,745,427 (16.5) | 7,145,393 (13.5) | 28,558,960 (53.9) | 5,290,468 (10.0) | 3,264,248 (6.2) |

| Read set | Chromosome | Median depth | Length (bp) | Low bp (%) | Half bp (%) | Medium bp (%) | High bp (%) | Outlier bp (%) |
|---|---|---|---|---|---|---|---|---|
| Unnamed Lowland HiFi | chr12 | 77 | 58,335,637 | 13,940,412 (23.9) | 6,070,451 (10.4) | 26,500,281 (45.4) | 6,920,670 (11.9) | 4,903,823 (8.4) |
| Unnamed Lowland HiFi | genome | 70 to 84 | 885,358,400 | 159,324,953 (18.0) | 109,654,442 (12.4) | 465,561,529 (52.6) | 96,302,444 (10.9) | 54,515,032 (6.2) |

**Supplementary Table S4:** Alleles carried by all three A-type cultivars and absent from Thille, with their carriers, amino-acid changes, and Fst status.

| Gene | A-shared allele | Amino-acid changes | In Unnamed Lowland | Carriers | Contig Fst rank (mean Fst) | Scaffold Fst rank (mean Fst) | Top 1% | Note |
|---|---|---|---|---|---|---|---|---|
| SDMYB | SDMYB[b] | N7S, D65G, S139G, D144N, T158I, T163M, A167T, S170N; frameshift from codon 183 | no | Gwen h2;Hass h2;Puebla h2 | 138 (0.283) | 157 (0.293) | both | |
| AGAMOUS-like | AGAMOUS-like[c] | 1 bp deletion at CDS position 5; frameshift, stop at codon 21 | no | Gwen h2;Hass h2;Puebla h2 | 274 (0.222) | 765 (0.173) | contig | |
| CSU2 | CSU2[d] | synonymous only | no | Gwen h1;Hass h1;Puebla h1 | 719 (0.168) | 326 (0.224) | scaffold | |
| NPR5-BOP2 | NPR5-BOP2[d] | V272A, I359V | no | Gwen h2;Hass h1;Puebla h1 | 1,925 (0.123) | 124 (0.316) | scaffold | |
| SGRL | SGRL[d] | S5Y, I101V | no | Gwen h1;Hass h1;Puebla h1 | 11,968 (0.056) | 22,445 (0.033) | none | |
| LCYB | LCYB[d] | L17V, E22G, D464E | no | Gwen h2;Hass h1;Puebla h1 | 5,297 | 5,869 | none | |

**Supplementary Table S5:** Read sets with one phased haplotype at 60 to 80% of the reads, with read depth over the gene and flanks and clipped-read clusters.

| label | gene | read set | af h1 | af h2 | h1 gene depth | h1 flank depth | h1 gene over flank | h2 gene depth | h2 flank depth | h2 gene over flank | all gene depth | all flank depth | all gene over flank | clip breakpoints | call |
|---|---|---|---|---|---|---|---|---|---|---|---|---|---|---|---|
| AGAMOUS-like | AGAMOUS-like | Gwen | 63.0 | 37.0 | 37.8 | 31.3 | 1.21 | 25.0 | 26.4 | 0.95 | 69.8 | 71.2 | 0.98 | 54814284:3reads(h1=0,h2=2,unphased=1);<br>54818285:3reads(h1=0,h2=0,unphased=3);<br>54820728:9reads(h1=0,h2=1,unphased=8);<br>54824469:7reads(h1=6,h2=0,unphased=1);<br>54825761:50reads(h1=0,h2=19,unphased=31);<br>54826832:48reads(h1=0,h2=21,unphased=27);<br>54835319:8reads(h1=0,h2=0,unphased=8);<br>54836004:3reads(h1=0,h2=0,unphased=3);<br>54837283:3reads(h1=1,h2=0,unphased=2);<br>54838021:15reads(h1=1,h2=0,unphased=14);<br>54838872:4reads(h1=2,h2=1,unphased=1);<br>54840383:8reads(h1=1,h2=6,unphased=1);<br>54842254:3reads(h1=2,h2=1,unphased=0);<br>54843950:51reads(h1=0,h2=21,unphased=30);<br>54844849:4reads(h1=1,h2=0,unphased=3);<br>54853339:3reads(h1=1,h2=0,unphased=2);<br>54853774:3reads(h1=0,h2=0,unphased=3);<br>54854630:3reads(h1=0,h2=0,unphased=3);<br>54856085:4reads(h1=1,h2=0,unphased=3);<br>54858544:56reads(h1=1,h2=22,unphased=33);<br>54871783:3reads(h1=0,h2=0,unphased=3);<br>54877206:3reads(h1=0,h2=0,unphased=3);<br>54883396:7reads(h1=0,h2=4,unphased=3);<br>54900965:10reads(h1=0,h2=7,unphased=3) | no depth signal; clip clusters present |
| AP2 | AP2 | Thille | 61.0 | 39.0 | 47.7 | 43.1 | 1.11 | 26.5 | 22.8 | 1.16 | 74.3 | 78.9 | 0.94 | none | no depth signal |
| CCR1_g514 | CCR1 | Gwen | 36.0 | 64.0 | 25.7 | 31.1 | 0.83 | 34.3 | 29.5 | 1.16 | 60.0 | 70.5 | 0.85 | 7958324:3reads(h1=2,h2=0,unphased=1);<br>7967544:8reads(h1=5,h2=0,unphased=3);<br>7984186:10reads(h1=0,h2=0,unphased=10);<br>7986748:6reads(h1=0,h2=0,unphased=6);<br>7991700:4reads(h1=0,h2=0,unphased=4);<br>7993538:3reads(h1=0,h2=1,unphased=2);<br>7997443:13reads(h1=12,h2=1,unphased=0);<br>8019316:4reads(h1=2,h2=0,unphased=2) | no depth signal; clip clusters present |
| CCR1_g514 | CCR1 | Thille | 63.0 | 37.0 | 55.4 | 45.8 | 1.21 | 30.9 | 26.8 | 1.16 | 86.3 | 84.2 | 1.03 | 7967544:3reads(h1=0,h2=0,unphased=3);<br>7984209:4reads(h1=0,h2=0,unphased=4);<br>7990945:17reads(h1=0,h2=0,unphased=17);<br>7991584:22reads(h1=0,h2=0,unphased=22);<br>7997443:14reads(h1=0,h2=14,unphased=0) | no depth signal; clip clusters present |

| label | gene | read set | af h1 | af h2 | h1 gene depth | h1 flank depth | h1 gene over flank | h2 gene depth | h2 flank depth | h2 gene over flank | all gene depth | all flank depth | all gene over flank | clip breakpoints | call |
|---|---|---|---|---|---|---|---|---|---|---|---|---|---|---|---|
| CCR1_g514 | CCR1 | Unnamed Lowland | 40.0 | 60.0 | 121.7 | 97.6 | 1.25 | 107.3 | 107.1 | 1.0 | 229.0 | 232.1 | 0.99 | 7967544:5reads(h1=0,h2=0,unphased=5); 7969223:10reads(h1=6,h2=0,unphased=4); 7980340:11reads(h1=0,h2=11,unphased=0); 7990856:4reads(h1=0,h2=0,unphased=4); 7991584:3reads(h1=0,h2=0,unphased=3); 7994887:3reads(h1=2,h2=0,unphased=1); 7995050:22reads(h1=16,h2=0,unphased=6); 7995591:3reads(h1=2,h2=0,unphased=1); 7996724:97reads(h1=45,h2=39,unphased=13); 8002146:8reads(h1=0,h2=0,unphased=8); 8013682:136reads(h1=38,h2=25,unphased=73); 8016670:7reads(h1=4,h2=0,unphased=3); 8018029:5reads(h1=2,h2=0,unphased=3); 8018777:3reads(h1=0,h2=0,unphased=3) | no depth signal; clip clusters present |
| CCS-like | CCS-like | Gwen | 61.0 | 39.0 | 33.6 | 29.4 | 1.14 | 29.7 | 27.2 | 1.09 | 63.3 | 63.7 | 0.99 | 58661373:7reads(h1=2,h2=1,unphased=4); 58702469:30reads(h1=0,h2=6,unphased=24); 58704967:10reads(h1=3,h2=0,unphased=7) | no depth signal; clip clusters present |
| CSU2 | CSU2 | Gwen | 40.0 | 60.0 | 33.3 | 30.1 | 1.1 | 45.6 | 41.6 | 1.1 | 79.0 | 82.0 | 0.96 | 390089:9reads(h1=5,h2=0,unphased=4); 399814:3reads(h1=0,h2=0,unphased=3); 407130:13reads(h1=13,h2=0,unphased=0); 414059:3reads(h1=1,h2=2,unphased=0); 417340:15reads(h1=5,h2=0,unphased=10); 420621:26reads(h1=6,h2=0,unphased=20); 420790:3reads(h1=0,h2=0,unphased=3) | no depth signal; clip clusters present |
| EMF1 | EMF1 | Gwen | 61.0 | 39.0 | 57.0 | 44.4 | 1.29 | 35.7 | 31.3 | 1.14 | 92.8 | 89.2 | 1.04 | 88422117:4reads(h1=0,h2=1,unphased=3); 88458256:3reads(h1=1,h2=0,unphased=2); 88459207:3reads(h1=1,h2=1,unphased=1); 88463590:4reads(h1=1,h2=2,unphased=1); 88465304:6reads(h1=3,h2=3,unphased=0) | no depth signal; clip clusters present |
| FBD-RNI-like | FBD-RNI-like | Unnamed Lowland | 38.0 | 62.0 | 31.0 | 31.6 | 0.98 | 49.8 | 42.2 | 1.18 | 80.8 | 84.4 | 0.96 | 15637899:26reads(h1=13,h2=0,unphased=13); 15638916:51reads(h1=0,h2=22,unphased=29); 15669872:22reads(h1=0,h2=8,unphased=14) | no depth signal; clip clusters present |
| ICR1 | ICR1 | Unnamed Lowland | 60.0 | 40.0 | 47.8 | 36.5 | 1.31 | 31.8 | 26.8 | 1.19 | 79.5 | 74.7 | 1.06 | 53803914:30reads(h1=14,h2=0,unphased=16) | no depth signal; clip clusters present |

| label | gene | read set | af h1 | af h2 | h1 gene depth | h1 flank depth | h1 gene over flank | h2 gene depth | h2 flank depth | h2 gene over flank | all gene depth | all flank depth | all gene over flank | clip breakpoints | call |
|---|---|---|---|---|---|---|---|---|---|---|---|---|---|---|---|
| LCYB | LCYB | Gwen | 68.0 | 32.0 | 58.6 | 47.9 | 1.22 | 33.0 | 28.9 | 1.14 | 91.6 | 86.0 | 1.06 | 70636235:10reads(h1=3,h2=2,unphased=5);<br>70654942:16reads(h1=0,h2=7,unphased=9);<br>70658817:18reads(h1=4,h2=3,unphased=11);<br>70661045:12reads(h1=4,h2=3,unphased=5);<br>70661474:4reads(h1=1,h2=0,unphased=3);<br>70663229:3reads(h1=0,h2=0,unphased=3);<br>70666212:4reads(h1=1,h2=2,unphased=1);<br>70674790:12reads(h1=0,h2=1,unphased=11);<br>70675233:14reads(h1=1,h2=1,unphased=12) | no depth signal; clip clusters present |
| LCYB | LCYB | Unnamed Low-land | 61.0 | 39.0 | 44.3 | 37.0 | 1.2 | 28.0 | 18.4 | 1.52 | 72.3 | 59.6 | 1.21 | 70658215:40reads(h1=1,h2=0,unphased=39);<br>70658817:9reads(h1=0,h2=0,unphased=9);<br>70661043:35reads(h1=0,h2=0,unphased=35);<br>70661772:50reads(h1=0,h2=13,unphased=37);<br>70664184:4reads(h1=1,h2=0,unphased=3);<br>70674753:20reads(h1=0,h2=8,unphased=12) | no depth signal; clip clusters present |
| NAC25 | NAC25 | Thille | 62.0 | 38.0 | 46.0 | 37.3 | 1.23 | 28.5 | 24.0 | 1.19 | 74.5 | 77.6 | 0.96 | 17102984:3reads(h1=3,h2=0,unphased=0);<br>17136265:13reads(h1=4,h2=0,unphased=9) | no depth signal; clip clusters present |
| NRX1 | NRX1 | Gwen | 62.0 | 38.0 | 47.8 | 41.1 | 1.16 | 39.7 | 32.0 | 1.24 | 87.5 | 89.5 | 0.98 | 78888330:12reads(h1=0,h2=4,unphased=8);<br>78891439:4reads(h1=0,h2=0,unphased=4);<br>78891650:5reads(h1=0,h2=0,unphased=5);<br>78892176:3reads(h1=0,h2=1,unphased=2);<br>78892441:15reads(h1=0,h2=3,unphased=12);<br>78893401:20reads(h1=0,h2=10,unphased=10);<br>78904443:8reads(h1=0,h2=0,unphased=8);<br>78905153:9reads(h1=1,h2=1,unphased=7);<br>78909843:22reads(h1=0,h2=0,unphased=22);<br>78911099:19reads(h1=0,h2=0,unphased=19);<br>78914991:24reads(h1=0,h2=15,unphased=9);<br>78923495:4reads(h1=0,h2=3,unphased=1) | no depth signal; clip clusters present |
| NRX1 | NRX1 | Thille | 60.0 | 40.0 | 56.8 | 42.8 | 1.33 | 32.9 | 18.4 | 1.79 | 89.8 | 77.6 | 1.16 | 78888337:29reads(h1=0,h2=15,unphased=14);<br>78892418:36reads(h1=0,h2=21,unphased=15);<br>78893401:34reads(h1=0,h2=24,unphased=10);<br>78909843:6reads(h1=0,h2=0,unphased=6);<br>78911095:7reads(h1=0,h2=0,unphased=7);<br>78914987:25reads(h1=0,h2=10,unphased=15) | no depth signal; clip clusters present |

| label | gene | read set | af h1 | af h2 | h1 gene depth | h1 flank depth | h1 gene over flank | h2 gene depth | h2 flank depth | h2 gene over flank | all gene depth | all flank depth | all gene over flank | clip breakpoints | call |
|---|---|---|---|---|---|---|---|---|---|---|---|---|---|---|---|
| NRX1 | NRX1 | Unnamed Low-land | 60.0 | 40.0 | 53.7 | 46.5 | 1.15 | 35.3 | 18.9 | 1.87 | 89.0 | 78.6 | 1.13 | 78888330:47reads(h1=9,h2=13,unphased=25); 78892418:40reads(h1=0,h2=20,unphased=20); 78893401:38reads(h1=0,h2=22,unphased=16); 78909841:70reads(h1=0,h2=0,unphased=70); 78910568:4reads(h1=0,h2=0,unphased=4); 78911095:69reads(h1=0,h2=0,unphased=69); 78914987:26reads(h1=0,h2=15,unphased=11) | no depth signal; clip clusters present |
| PAR1 | PAR1 | Gwen | 63.0 | 37.0 | 27.5 | 25.5 | 1.08 | 20.7 | 23.2 | 0.89 | 48.2 | 65.5 | 0.74 | 7894950:5reads(h1=0,h2=2,unphased=3); 7897136:7reads(h1=0,h2=6,unphased=1); 7905382:3reads(h1=0,h2=1,unphased=2); 7908405:3reads(h1=1,h2=0,unphased=2); 7909717:26reads(h1=0,h2=0,unphased=26); 7910507:46reads(h1=1,h2=32,unphased=13); 7911423:21reads(h1=1,h2=10,unphased=10); 7912183:20reads(h1=0,h2=10,unphased=10); 7912563:34reads(h1=3,h2=0,unphased=31); 7912745:39reads(h1=0,h2=28,unphased=11); 7913272:3reads(h1=2,h2=0,unphased=1); 7915055:5reads(h1=2,h2=1,unphased=2); 7917625:6reads(h1=2,h2=3,unphased=1); 7918747:3reads(h1=1,h2=2,unphased=0); 7921565:22reads(h1=0,h2=0,unphased=22); 7922539:4reads(h1=0,h2=0,unphased=4); 7924951:47reads(h1=0,h2=1,unphased=46); 7926041:3reads(h1=0,h2=0,unphased=3); 7926599:62reads(h1=0,h2=18,unphased=44); 7934872:3reads(h1=0,h2=3,unphased=0); 7935821:4reads(h1=0,h2=0,unphased=4); 7946094:3reads(h1=0,h2=0,unphased=3) | one haplotype missing over part of the gene (all-read depth <= 0.75x flanks); clip clusters present |
| PAR1 | PAR1 | Thille | 26.0 | 74.0 | 19.0 | 26.3 | 0.72 | 46.0 | 37.9 | 1.21 | 65.0 | 90.0 | 0.72 | 7894950:8reads(h1=2,h2=0,unphased=6); 7897135:3reads(h1=3,h2=0,unphased=0); 7904973:3reads(h1=0,h2=0,unphased=3); 7905396:5reads(h1=5,h2=0,unphased=0); 7909717:34reads(h1=0,h2=0,unphased=34); 7910507:43reads(h1=34,h2=0,unphased=9); 7911423:19reads(h1=10,h2=0,unphased=9); 7912187:31reads(h1=20,h2=0,unphased=11); 7912564:42reads(h1=0,h2=1,unphased=41); 7912749:26reads(h1=16,h2=0,unphased=10); 7917624:8reads(h1=8,h2=0,unphased=0); 7921565:39reads(h1=0,h2=0,unphased=39); 7924951:28reads(h1=0,h2=0,unphased=28); 7926594:24reads(h1=6,h2=0,unphased=18); 7934872:21reads(h1=7,h2=0,unphased=14) | one haplotype missing over part of the gene (all-read depth <= 0.75x flanks); clip clusters present |

| label | gene | read set | af h1 | af h2 | h1 gene depth | h1 flank depth | h1 gene over flank | h2 gene depth | h2 flank depth | h2 gene over flank | all gene depth | all flank depth | all gene over flank | clip breakpoints | call |
|---|---|---|---|---|---|---|---|---|---|---|---|---|---|---|---|
| SDMYB | SDMYB | Gwen | 70.0 | 30.0 | 40.8 | 43.6 | 0.94 | 20.5 | 7.8 | 2.62 | 61.3 | 58.6 | 1.05 | 10301790:7reads(h1=2,h2=0,unphased=5);<br>10310867:31reads(h1=14,h2=0,unphased=17);<br>10311404:33reads(h1=0,h2=21,unphased=12);<br>10316752:4reads(h1=0,h2=1,unphased=3);<br>10317551:5reads(h1=0,h2=1,unphased=4);<br>10320535:25reads(h1=0,h2=14,unphased=11);<br>10325706:19reads(h1=1,h2=6,unphased=12);<br>10326128:3reads(h1=0,h2=1,unphased=2);<br>10326589:25reads(h1=0,h2=4,unphased=21);<br>10329601:3reads(h1=0,h2=0,unphased=3);<br>10330705:3reads(h1=0,h2=0,unphased=3) | no depth signal; clip clusters present |
| SGRL | SGRL | Gwen | 38.0 | 62.0 | 27.9 | 25.4 | 1.1 | 39.1 | 37.6 | 1.04 | 67.1 | 77.9 | 0.86 | 5804045:5reads(h1=4,h2=0,unphased=1);<br>5832766:6reads(h1=0,h2=0,unphased=6);<br>5833476:37reads(h1=11,h2=0,unphased=26);<br>5848039:4reads(h1=1,h2=1,unphased=2);<br>5850176:3reads(h1=0,h2=0,unphased=3);<br>5850331:3reads(h1=0,h2=0,unphased=3);<br>5851042:3reads(h1=0,h2=0,unphased=3);<br>5851847:23reads(h1=0,h2=0,unphased=23);<br>5852114:28reads(h1=1,h2=0,unphased=27);<br>5852855:3reads(h1=1,h2=0,unphased=2);<br>5871778:3reads(h1=2,h2=0,unphased=1) | no depth signal; clip clusters present |
| SGRL | SGRL | Thille | 40.0 | 60.0 | 26.1 | 22.6 | 1.15 | 38.5 | 31.2 | 1.23 | 64.5 | 65.8 | 0.98 | 5833476:13reads(h1=3,h2=0,unphased=10);<br>5850225:3reads(h1=0,h2=0,unphased=3) | no depth signal; clip clusters present |
| SPA1 | SPA1 | Thille | 62.0 | 38.0 | 37.0 | 34.4 | 1.08 | 24.8 | 20.9 | 1.19 | 61.8 | 69.0 | 0.9 | 31244347:15reads(h1=2,h2=6,unphased=7);<br>31244725:4reads(h1=1,h2=0,unphased=3);<br>31245839:3reads(h1=0,h2=0,unphased=3);<br>31250165:25reads(h1=6,h2=6,unphased=13);<br>31253636:28reads(h1=0,h2=4,unphased=24);<br>31254846:13reads(h1=0,h2=0,unphased=13) | no depth signal; clip clusters present |
| TIC | TIC | Gwen | 71.0 | 29.0 | 38.4 | 33.9 | 1.13 | 23.9 | 23.4 | 1.02 | 62.3 | 65.3 | 0.95 | 23298976:5reads(h1=0,h2=0,unphased=5);<br>23302001:8reads(h1=1,h2=0,unphased=7);<br>23321660:41reads(h1=0,h2=9,unphased=32) | no depth signal; clip clusters present |
| TIC | TIC | Thille | 60.0 | 40.0 | 42.7 | 30.9 | 1.38 | 28.3 | 21.7 | 1.3 | 71.0 | 63.2 | 1.12 | 23298993:11reads(h1=0,h2=0,unphased=11);<br>23300619:7reads(h1=0,h2=0,unphased=7);<br>23302001:6reads(h1=0,h2=0,unphased=6);<br>23321660:40reads(h1=0,h2=4,unphased=36) | no depth signal; clip clusters present |

| label | gene | read set | af h1 | af h2 | h1 gene depth | h1 flank depth | h1 gene over flank | h2 gene depth | h2 flank depth | h2 gene over flank | all gene depth | all flank depth | all gene over flank | clip breakpoints | call |
|---|---|---|---|---|---|---|---|---|---|---|---|---|---|---|---|
| VOZ1_g26844 | VOZ1 | Gwen | 79.0 | 21.0 | 10.2 | 3.8 | 2.72 | 10.8 | 3.5 | 3.1 | 65.0 | 61.1 | 1.06 | 54328473:38reads(h1=0,h2=0,unphased=38); 54330118:17reads(h1=1,h2=0,unphased=16); 54340491:74reads(h1=8,h2=9,unphased=57); 54343905:3reads(h1=0,h2=0,unphased=3); 54344742:3reads(h1=0,h2=0,unphased=3); 54346401:81reads(h1=7,h2=11,unphased=63); 54347507:3reads(h1=0,h2=0,unphased=3); 54349233:3reads(h1=0,h2=0,unphased=3); 54349760:3reads(h1=0,h2=0,unphased=3) | no depth signal; clip clusters present |
| ZEP | ZEP | Gwen | 73.0 | 27.0 | 49.2 | 37.7 | 1.3 | 28.2 | 9.0 | 3.13 | 77.6 | 71.4 | 1.09 | 10922370:3reads(h1=0,h2=0,unphased=3); 10924702:28reads(h1=0,h2=9,unphased=19); 10932103:3reads(h1=2,h2=1,unphased=0); 10932753:3reads(h1=1,h2=0,unphased=2); 10933112:6reads(h1=0,h2=1,unphased=5); 10933275:5reads(h1=0,h2=0,unphased=5); 10936354:3reads(h1=0,h2=2,unphased=1); 10937261:3reads(h1=1,h2=0,unphased=2); 10937525:3reads(h1=1,h2=0,unphased=2); 10941340:3reads(h1=1,h2=2,unphased=0); 10944769:72reads(h1=0,h2=20,unphased=52); 10946827:5reads(h1=0,h2=0,unphased=5); 10949997:3reads(h1=0,h2=0,unphased=3) | no depth signal; clip clusters present |
| ZEP | ZEP | Thille | 40.0 | 60.0 | 36.5 | 38.1 | 0.96 | 50.4 | 41.9 | 1.2 | 86.8 | 96.3 | 0.9 | none | no depth signal |
| ZFP2 | ZFP2 | Gwen | 60.0 | 40.0 | 53.0 | 50.1 | 1.06 | 44.0 | 37.6 | 1.17 | 97.0 | 104.3 | 0.93 | 2524817:4reads(h1=0,h2=2,unphased=2); 2547828:3reads(h1=0,h2=0,unphased=3); 2549139:6reads(h1=0,h2=0,unphased=6); 2551564:3reads(h1=1,h2=0,unphased=2); 2553885:3reads(h1=0,h2=2,unphased=1); 2554240:3reads(h1=1,h2=0,unphased=2); 2558553:3reads(h1=2,h2=1,unphased=0); 2560651:7reads(h1=0,h2=3,unphased=4); 2561060:5reads(h1=0,h2=1,unphased=4); 2563902:5reads(h1=0,h2=0,unphased=5); 2564582:3reads(h1=0,h2=0,unphased=3); 2564768:7reads(h1=0,h2=0,unphased=7); 2567687:3reads(h1=0,h2=1,unphased=2); 2567946:3reads(h1=0,h2=0,unphased=3) | no depth signal; clip clusters present |
